# Accuracy of predicting chemical body composition of growing pigs using dual-energy X-ray absorptiometry

**DOI:** 10.1101/2020.09.15.286153

**Authors:** Claudia Kasper, Patrick Schlegel, Isabel Ruiz-Ascacibar, Peter Stoll, Giuseppe Bee

## Abstract

Studies in animal science assessing nutrient and energy efficiency or determining nutrient requirements necessitate gathering exact measurements of body composition or body nutrient contents. Wet chemical analysis methods or standardized dissection are commonly applied, but both are destructive. Harnessing human medical imaging techniques for animal science can enable repeated measurements of individuals over time and reduce the number of individuals required for research. Among imaging techniques, dual-energy X-ray absorptiometry (DXA) is particularly promising. However, the measurements obtained with DXA do not perfectly match dissections or chemical analyses, requiring the adjustment of the DXA via calibration equations. Several calibration regressions have been published, but comparative studies are pending. Thus, it is currently not clear whether existing regression equations can be directly used to convert DXA measurements into chemical values or whether each individual DXA device will require its own calibration. Our study builds prediction equations that relate body composition to the content of single nutrients in growing entire male pigs (body weight range 20-100 kg) as determined by both DXA and chemical analyses, with R^2^ ranging between 0.89 for ash and 0.99 for water and crude protein. Moreover, we show that the chemical composition of the empty body can be satisfactorily determined by DXA scans of carcasses, with the prediction error rCV ranging between 4.3% for crude protein and 12.6% for ash. Finally, we compare existing prediction equations for pigs of a similar range of body weights with the equations derived from our DXA measurements and evaluate their fit with our chemical analyses data. We found that existing equations for absolute contents that were built using the same DXA beam technology predicted our data more precisely than equations based on different technologies and percentages of fat and lean mass. This indicates that the creation of generic regression equations that yield reliable estimates of body composition in pigs of different growth stages, sexes and genetic breeds could be achievable in the near future. DXA may be a promising tool for high-throughput phenotyping for genetic studies, because it efficiently measures body composition in a large number and wide array of animals.

## Introduction

For studies in animal science that include measurements of nutrient and energy efficiency or the determination of protein, mineral and energy requirements, it is necessary to determine exact body composition. Two destructive methods are commonly used to determine body composition. Carcass dissection into fat, lean and bone tissue is used to determine carcass quality for nutrition and genetic selection studies. Wet chemistry analyses of the carcass are carried out to determine the nutrient content, and thus the nutrient and energy deposition rate and efficiency. The latter method certainly offers better accuracy and less operator bias, and it is commonly considered the gold standard. But both techniques are destructive, expensive and time-consuming, which precludes their application in studies requiring repeated measurements on individuals or large samples sizes. However, assessment of body composition or nutrient content in vivo could be used to evaluate the influence of feeding strategies, housing systems or environmental factors on the development of body composition or could facilitate monitoring nutrient efficiency. For genetics research, determining body composition and nutrient content with non-destructive methods could aid performance testing and the selection of parent individuals for breeding. Therefore, harnessing medical imaging techniques originally designed for humans, such as computed tomography (CT), magnetic resonance imaging (MRI) and dual-energy X-ray absorptiometry (DXA), is a promising area for animal science (Scholz et al., 2015). Imaging methods can be performed on carcasses or in vivo under anaesthesia, which enables repeated measurements of individuals through time, and can reduce the number of individuals required for research, as demanded by the 3R principles (National Centre for the Replacement, Refinement and Reduction of Animals in Research, 2019).

Medical imaging techniques such as DXA, CT and MRI not only enable repeated measurements of the same individual but also yield more reproducible results than dissections. Among medical imaging techniques, DXA has advantages that make it particularly attractive for application in animal science (Pomar et al., 2017), which mainly include financial and work security aspects. Compared to CT and MRI, DXA is easier to use, has lower instrument and operating costs, a more rapid scan speed, little to no operator bias and requires less image processing, consequently allowing for quick data analyses (Scholz et al., 2015). Moreover, ionizing radiation emitted by DXA is relatively low (Genton et al., 2002), making it a secure device for operators. The DXA provides information about lean mass, fat tissue mass and bone mineral content (BMC), the sum of which represents the total body mass. It also measures the bone surface, allowing for determination of bone mineral density (BMC divided by bone surface). However, because DXA is an indirect tool, it does not provide true values, although its values are precise and have, for example, less incidence of random error (Kipper et al., 2019a). Thus, the information from DXA devices does not fully correspond with the data from invasive techniques, and it is therefore necessary to adjust measurements to reference dissections or, ideally, chemical analyses via prediction equations (Mitchell et al., 1998b). Regression equations have been published for a range of breeds, sexes and weight classes of pigs, using different DXA beam technologies (Mitchell et al., 1998a; Mitchell et al., 1998c; Pomar et al., 2001; Mitchell et al., 2003; Suster et al., 2003; Pomar et al., 2017; Kipper et al., 2019b). The image-processing software on DXA devices used to compute tissue masses was developed for the human body. It is optimized for scanning humans in a supine position, and it might rely on human-specific assumptions about the distribution of fat, lean tissue and bone tissue (Genton et al., 2002; Hunter et al., 2011). Moreover, because algorithms are proprietary, these assumptions cannot be checked or adapted to animal species.

The results of imaging techniques should, in principle, be independent of animal size and body composition, and therefore, regression equations might not require adjustment for different growth stages, sexes and genetic breeds (Mitchell et al., 1998a; Marcoux et al., 2005; Soladoye et al., 2016). To date, no comparative study has evaluated the fit of previously published regression equations to a new set of data. Thus, it is unclear whether existing regression equations can be directly used to convert DXA measurements into chemical values or whether each individual DXA device requires its own calibration. The objectives of this study on pigs of 20 to 100 kg were 1) to estimate the chemical composition of the empty body (EB) from values determined by DXA on the live animal; 2) to estimate the chemical composition of the carcass from the values determined by DXA on carcass; 3) to estimate the chemical composition of the EB from the carcass values determined by DXA, over the complete range of body weights. Finally, we compared existing prediction equations with the equations derived from our data and evaluated their fit with our data.

## Methods

### Animals

We used 68 entire males originating from 17 litters in two farrowing series of the Agroscope Large White pig dam line. The details of housing, feeding and slaughter procedures were the same as described by Ruiz-Ascacibar et al. (2017). In brief, pigs with a mean BW of 8.9 kg (± 2.1 kg SD) were selected at weaning and and kept in 17.35 m^2^ pens in groups of a maximum of 14. The pigs had ad libitum access to a diet based on BW, with the pigs having < 20 kg BW receiving a weaning diet, those between 20 and 60 kg BW getting one of two grower diets and pigs with > 60 kg BW (20–60 kg BW) being fed one of two finisher diets. The grower and finisher diets differed by 20% in their CP content (165 vs 139 g CP per kg grower diet and 140 vs 117 g CP per kg finisher diet). Both diets were formulated to contain the same amount of digestible energy (13.2 MJ/kg feed). Pigs were either slaughtered at a live BW of 20 kg (N = 6) or kept on feed until they reached a target BW of 60 kg (N = 18) or 100 kg (N = 37 for EB, and N = 44 for carcass), as determined by weekly weighing (Grüter SST-WA-03, Eschenbach, Switzerland; ± 100 g). Three days prior to slaughter (−3), the pigs were fasted for at least 16h and then sedated using isoflurane (Attane, Piramal Critical Care, Inc., Bethlehem, PA, USA) prior to DXA scanning. Sedation was conducted to prevent movement during the scanning procedure. The pigs were scanned on the DXA and subsequently their BW was determined using a scale (BW_−3_). The awakened pigs were brought back to their respective pens. The live DXA scans under anaesthesia were carried out three days before slaughter in order to meet the required withdrawal period for isoflurane before the sale of carcasses for human consumption. A technical problem occurred on the DXA that disabled scanning for a period of two weeks. Thus, live scans of seven 100 kg category pigs could not be conducted. After 16 h of feed withdrawal, the pigs were slaughtered using the Agroscope experimental abattoir for exsanguination after CO2 stunning. Pigs were weighed (Berkel IWS IT 6000, Berkel-Obrecht, Spreitenbach, Switzerland; ±200 g) before and after exsanguination to determine blood mass. The BW prior to exsanguination corresponds to the BW at slaughter (BWslaughter). Immediately after exsanguination, hair and claws were removed and collected. Animals were then eviscerated and the carcass and head were sawed into two halves. The two half carcasses including the head without brain were weighed for correspondence with the warm carcass weight. Then, they were chilled at 2°C and a cold carcass weight was determined 24 h post mortem. The left carcass halves (including halved head without brain) were then processed into primal cuts (head, neck, shoulder, ham, loin, tail and feet) and scanned using DXA. Subsequently, the cuts were frozen for further processing to determine the carcass chemical composition as described by Ruiz-Ascacibar et al. (2017). To prevent thawing and re-freezing, the carcass halves were divided into primal cuts before scanning because grinding prior to chemical analysis is always performed in a frozen state. The carcass cuts of the seven pigs slaughtered during the DXA failure were exceptions in being thawed for the DXA scanning and then re-frozen. The gall bladder, bladder, stomach, intestine and hind gut were emptied and rinsed with clean water to remove any remaining bile, urine and digesta. The eviscerated organs and brain were then pooled and homogenized for further processing in order to determine the chemical composition of the EB as described by Ruiz-Ascacibar et al. (2017).

### DXA image acquisition

A GE Lunar DXA (i-DXA, GE Healthcare Switzerland, Glattbrugg, Switzerland) with a narrow-angle fan beam (Collimator Model 42129) was used to scan the live pigs and carcass halves. The DXA has a 100 kV generator and K-edge filtering, resulting in approximately 39 keV and 71 keV X-ray effective energies. The calibration was checked and passed before each scanning session by scanning a calibration phantom according to the manufacturer’s instructions. Scans were conducted using the ‘total body thick’ mode (0.188 mA, scan speed 80mm/s according to Carver et al., 2013) with enCORE software (version 16). It has to be noted that, according to the manufacturer, this mode is not recommended for pigs of smaller sizes. In this study, the mode with the highest level of radiation was used for all individuals regardless of size. This was done in order to avoid switching between modes, which could introduce additional variation between the measurements of pigs of different BW classes. The sedated pigs were placed on the DXA table in a standardized way in a prone position with the hind legs extended while the front legs were positioned along the side but kept away from the body by two wedges of foam plastic (Figure 1a). The left carcass cuts were placed next to each other on the DXA table with the skin side facing downwards. The cuts were arranged in a standardized manner to reflect an uncut half carcass (Figure 1b). All scans started from the head.

**Figure 1.**
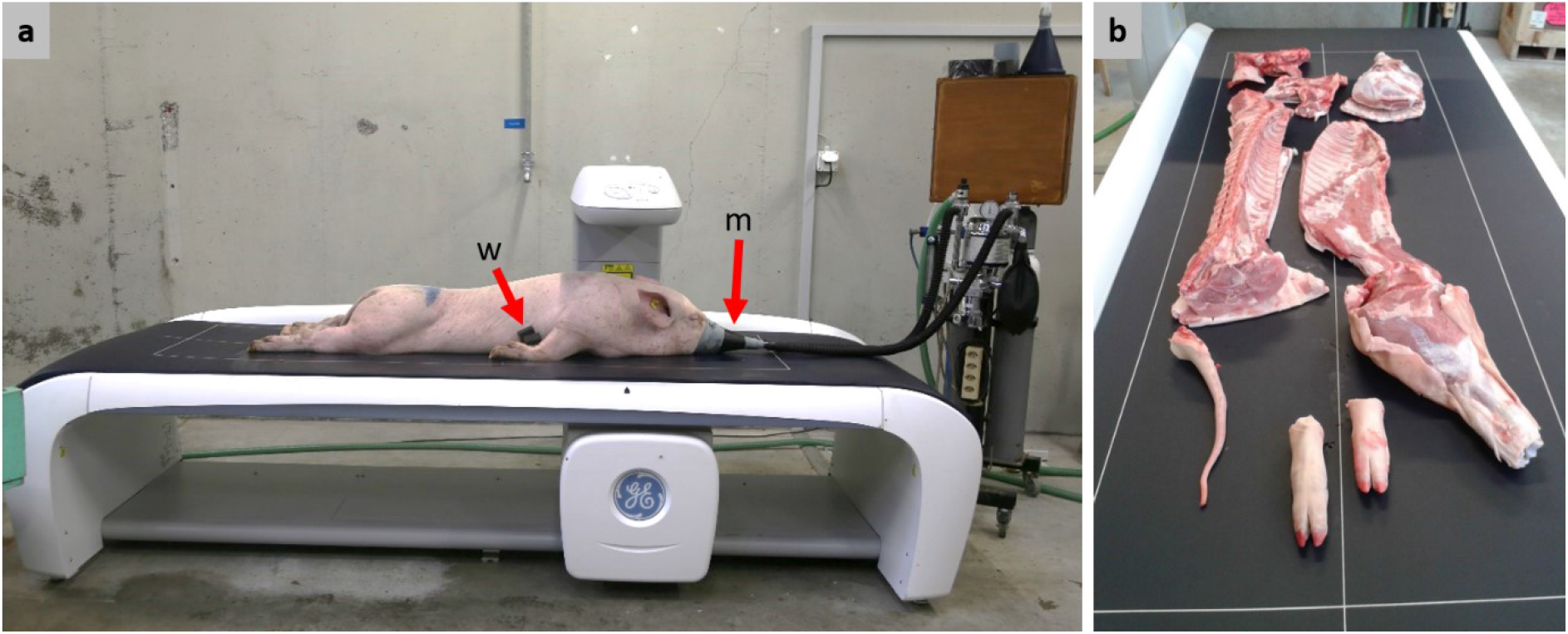
**(a)** Positioning of live animals on the DXA table for scanning under anaesthesia. **w:** Plastic foam wedge used to keep the front legs away from the body. **m:** Mask for the continuous administration of isoflurane. **(b)** Positioning of cut carcass on the DXA table.

### DXA image processing

Scan images from live animals were processed to remove artefacts (i.e. regions not belonging to the animal, such as the mask and tube on the sedating apparatus and the ear tag) and to position the regions of interest (ROI). The ROI were placed in order to be similar to the human ROI according to the supplier’s guidelines. This was done by defining reference points for the pig (Supplementary Material Section 1, Figure S1). The ROI in the scans of carcasses were set on “right arm" for all primal cuts. The variables ‘total mass’, ‘BMC’ (bone mineral content), ‘lean’ and ‘fat’ of the live whole body and of each left carcass half (listed in the upper part of Table 1) were exported from the software and used for further analysis.

**Table 1.**
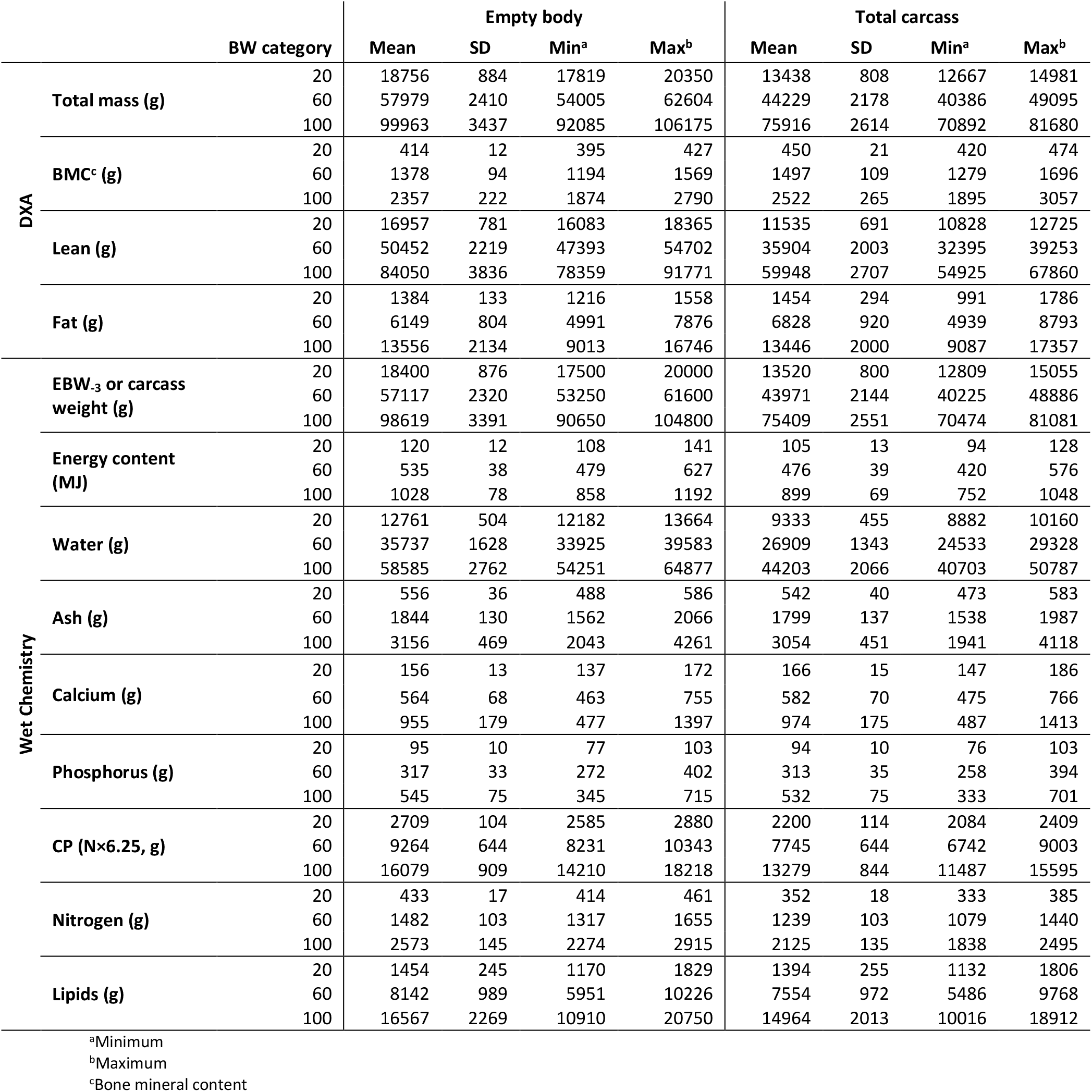
Body and carcass composition of growing pigs determined at 20, 60 and 100 kg body weight

### Laboratory analyses and calculations

Chemical analyses of energy, water, ash, Ca, P, N and lipid content (listed in the lower part of Table 1) were conducted as described in Ruiz-Ascacibar et al. (2017; 2019). In brief, all fractions of the EB_slaughter_, which were 1) the pooled left carcass cuts, including the left half of the head without brain, 2) the pooled empty digestive tract, the organs and brain, 3) the hair and claws and 4) the blood, were ground, homogenised and dried separately. For the EB_slaughter_, all calculations of contents were based on warm carcass weight (to closely mimic the situation in the live animal), and for the carcass, they were based on cold carcass weight. The contents of energy, water, ash, Ca, P, N and lipids in the left carcass half were combined with those of the right carcass half to obtain values for the full carcass. The CP content was calculated by multiplying the N content determined by wet chemistry by 6.25. Analyses were conducted in duplicate, but if results differed by more than 5%, up to four replicates were analysed. Since the live pigs were scanned by DXA three days prior to slaughter, the chemical composition of the EB at that time (EB-3) was obtained individually from the EB_slaughter_ values as follows: The chemical composition of EB_slaughter_ was divided by EBWslaughter and multiplied by EBW-3. The EBW-3 corresponded to the BW-3 multiplied by 95.3%, 96.2% or 95.6% (the average ratio of EBW slaughter to BW slaughter for the 20, 60 and 100 kg category pigs, respectively).

### Statistical analysis of results

The plausibility of the data was checked before the statistical analysis (Supplementary Material Section 2), including a check of the differences between the total levels of raw ash, CP, lipids and water and the EBWslaughter and cold carcass weight obtained by scales. Histograms of the raw data obtained by chemical analyses (Figure S2) were visually checked. In all analyses, the experimental unit was the individual pig. The code on which statistical analyses were based in R (version 3.6.3; R Core Team, 2020) can be found in the Supplementary Material. For the comparison of chemical and DXA-derived values, we conducted Welch’s two-sample t-test for unequal variances. To compute the difference, we subtracted the mean of the respective chemical value from the DXA value and expressed it as a percentage of the chemical value. Regression equations were computed with the lm() command, and figures were created using the package ggplot2 (version 3.2.1; Wickham, 2016). BW_DXA_, energy_DXA_, water_DXA_, ash_DXA_, Ca_DXA_, P_DXA_, CP_DXA_, N_DXA_ and lipid_DXA_ were used to denote the values predicted by the regression equations. To facilitate interpretation, we kept variables on their original scale in all models. The models for predicting EBW-3 and carcass weight contained total mass by DXA as the sole predictor variable. A stepwise reduction of linear models was used to select estimates of body composition provided by DXA. The full models for the prediction of all chemical parameters included BMC, lean tissue and fat tissue as estimated by DXA. In case more than one of the predictors of the full model was not significant, we first omitted the variable from which, from a biological point of view, we expected the weakest relationship to the estimated chemical content. Because the data stem from an experiment with different dietary treatments, we examined whether dietary treatment had an effect on the slopes and intercepts of the regression equations (Supplementary Material Section 3). The effect of dietary treatment was never significant and the inclusion of the dietary treatment changed slopes and intercepts only marginally (Table S1); thus, we present models without dietary treatments. Adjusted R^2^ as a measure of goodness-of-fit was extracted from the R model object, and the root mean square error (RMSE) was calculated as 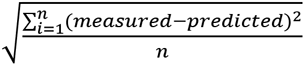 where measured is the value determined by chemical analysis, predicted is the value estimated from the DXA measurement via the regression equation and n is the sample size. To facilitate comparison among variables with a wide range of contents in the body, we computed the residual coefficient of variation (rCV), which is the percentage of the RMSE to the chemically measured value, as a standardized metric. We performed cross-validation of the regression equations via a jack-knife approach by iteratively omitting one data point from the data set and computing the prediction error, using the leave-one-out method in the caret package (version 6.0-86; Kuhn, 2020). The models did not fully comply with the assumptions for linear least squares regressions (Supplementary Material Section 4), which could possibly lead to a misestimation of the parameters. Thus, using the leave-one-out cross-validation procedure, we checked whether the estimates of intercepts and slopes were robust or if they deviated from our estimates due to a disproportionate leverage of certain data points. This also allowed us to derive non-parametric confidence intervals for slopes and intercepts (for details, see Supplementary Material Section 4). Even though our main goal was to build regression equations over the whole range of slaughter weights, we also present specific (regional) regressions for each target slaughter-weight category (20, 60 and 100 kg, see Supplementary Material Section 5). This was done to compare the slopes and intercepts of regional regressions with the ones of the global regression. However, these are probably less reliable since they are based on only a subset of the data, and the number of observations for each regression is low, in particular for the 20 kg category.

Finally, we compared the prediction equations we obtained from our data set with those published in the literature (Mitchell et al., 1998a; Mitchell et al., 1998c; Pomar et al., 2001; Mitchell et al., 2003; Pomar et al., 2017; Kipper et al., 2019b). To do this, we applied the published equation to the DXA values of our data set and computed the RMSE of the values predicted using the published equations and the chemically measured values from our study.

## Results and Discussion

### Correspondence of wet chemistry and DXA measurements

Table 1 shows the descriptive statistics of DXA values from the live pigs three days prior to slaughter and from the carcass, the scale weight of the EB-3 and carcass and the wet-chemistry values from the EB-3 and the carcass, respectively. Table 2 shows the results of testing for significant differences between DXA and chemical values. Scale weight and DXA weights of the EB and carcass were comparable (P > 0.05), with a mean difference of 1% in both EB and carcass. As noted previously, the exact correspondence of body weight measured by DXA and weigh scales does not necessarily mean that the body composition determined by the two measuring methods is equally accurate (Mitchell et al., 1998b). In contrast to body weight, the values for ash, CP and lipids obtained from chemical analyses differed (P < 0.01) from the BMC, lean tissue and fat mass obtained by DXA, except for the values of lipids and fat in the carcass (P > 0.05). Such observations were reported previously (Suster et al., 2003). The DXA readout does not provide information on water content. Therefore, the water content in the fat-free mass (sum of CP, ash and water), which was determined by chemical methods, must be taken into account. It represented 76% (± 1.46% SD; CV=1.91%) in the EB and 74% (± 1.47% SD; CV=2.00%) in the carcass. This content is in line with the reported 73% to 77% water in fat-free mass (Speakman, 1997; Bocquier et al., 1999; Lerch et al., 2015). Because the major proportion of body water is in the fat-free mass and because DXA derives lean tissue mass indirectly via body water (Hunter et al., 2011), the water may be attributed with CP to DXA lean. The sum of CP and water mass was comparable (P > 0.05) with lean mass. The higher concordance between lean mass and CP with water in the carcass in comparison to the EB could be explained by gut water content and/or a lipid-to-protein ratio in the intestines and organs that differs from that in the carcass.

**Table 2.** Means and standard deviations of variables measured by DXA and determined by chemical analyses (or scales) and the results of Welch’s two-sample t-test comparing DXA and chemical values in growing pigs

| | Item DXA | Item Chemical | DXA [g] | Chemical [g] | $\Delta\%$<sup>a</sup> | $t^b$ | df | P |
| --- | --- | --- | --- | --- | --- | --- | --- | --- |
| Empty Body | Total mass | EBW<sub>-3</sub> | 79587 ± 27838 | 78482 ± 27504 | 1 | 0.22 | 119.98 | 0.826 |
|  | BMC<sup>c</sup> | Ash | 1877 ± 680 | 2513 ± 954 | -34 | -4.22 | 108.47 | <0.001 |
|  | Lean | CP | 67537 ± 22844 | 12753 ± 4602 | 81 | 18.36 | 64.86 | <0.001 |
|  | Lean | CP + water | 67537 ± 22844 | 60089 ± 20181 | 11 | 1.91 | 118.20 | 0.059 |
|  | Fat | Lipid | 10173 ± 4749 | 12594 ± 5609 | -24 | -2.57 | 116.83 | 0.011 |
| Carcass | Total mass | Carcass weight | 62016 ± 20708 | 61626 ± 20521 | 1 | 0.11 | 133.99 | 0.913 |
|  | BMC<sup>c</sup> | Ash | 2068 ± 711 | 2500 ± 901 | -21 | -3.1 | 127.11 | 0.002 |
|  | Lean | CP | 49312 ± 16005 | 10836 ± 3705 | 78 | 19.31 | 74.16 | <0.001 |
|  | Lean | CP + water | 49312 ± 16005 | 47385 ± 15215 | 4 | 0.72 | 133.66 | 0.473 |
|  | Fat | Lipid | 10636 ± 4407 | 11806 ± 4895 | -11 | -1.46 | 132.55 | 0.146 |

<sup>a</sup>Difference between DXA and chemical values as a percentage of chemical value
<sup>b</sup>Statistic from Welch's two-sample t-test
<sup>c</sup>Bone mineral content

The BMC was 34% and 21% lower (P < 0.01) than the ash content of the EB and carcass, respectively. The skeleton is rich in minerals and contains a high proportion of certain minerals, such as Ca and P. However, other minerals found in body ash are mainly present in body tissues and fluids, so they are not considered as part of BMC. Thus, BMC is expected to be lower than body ash. When comparing BMC with ash in single bones, the correspondence of the values was greatly improved (Schlegel and Gutzwiller, 2020). Fat tissue mass was 24% lower (P < 0.05) than lipid content in the EB but comparable (P > 0.05) in the carcass. The fat content of the gastrointestinal tract, the organs and the visceral adipose tissue (see Weigand et al., 2020) may have caused this discrepancy.

Given the expected discrepancies between the values obtained by DXA and wet chemistry, the conversion of DXA estimates of body composition via prediction equations was necessary to obtain quantitative estimates of body composition. The prediction equations had similar high precision and accuracy for EB and carcass. The non-parametric confidence intervals show that estimates of slopes and intercepts were robust (Table 3) even though model diagnostics were not optimal (see Supplementary Material Section 4).

**Table 3.**
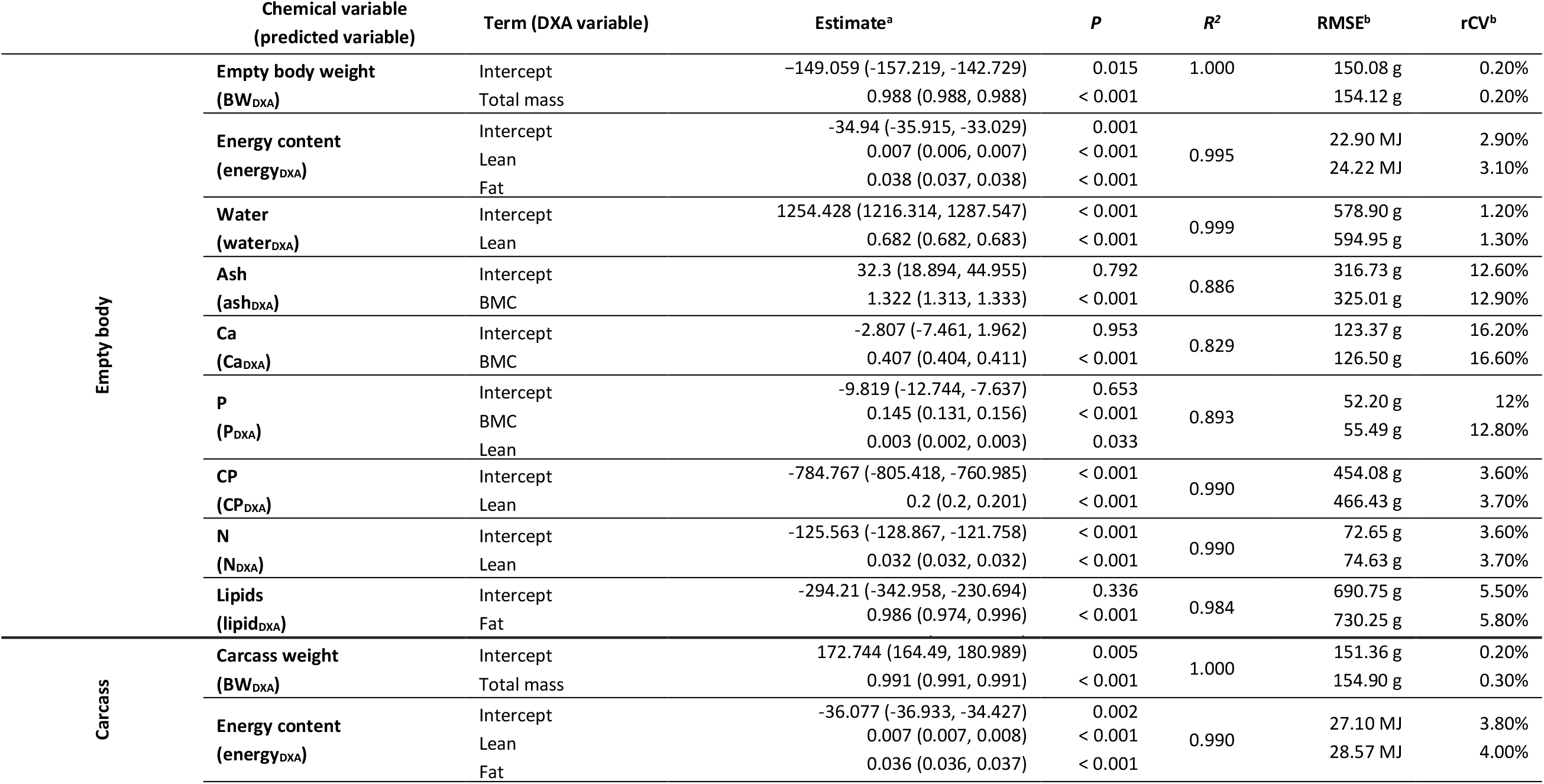
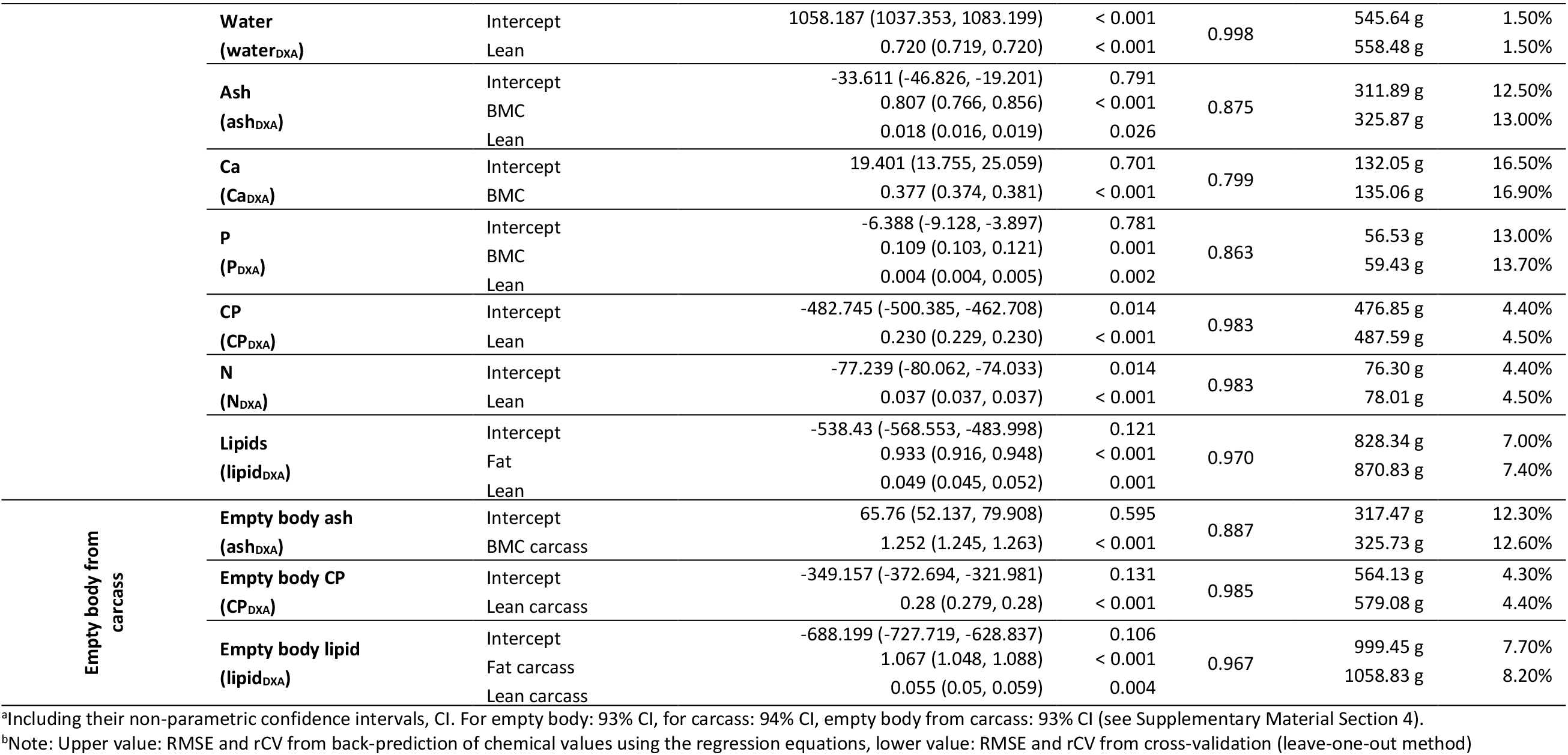
Models to estimate chemical composition of the empty body and the carcass from DXA values

#### Weight

The EBW and carcass weight could be predicted by the regression equations using DXA total mass with the highest R^2^ and lowest rCV (Table 3, Figure 2, Figure 3).This was expected given the non-significant difference between the weights obtained by scales and DXA (Table 2). The RMSEs in the EB and the carcass were very similar, and we obtained an excellent fit (R^2^ = 1 after rounding) with both EB and carcass. The intercept of the prediction equation was slightly shifted below zero for the EB, and above zero for the carcass. The slope was almost 1 in the EB and slightly below 1 in the carcass. This shows that DXA can estimate the BW with very high reliability. The coefficients of regional regressions for each target slaughter-weight category were very similar to the global regression and all obtained very high R^2^ (Table S2, Figures S3 and S4).

**Figure 2.**
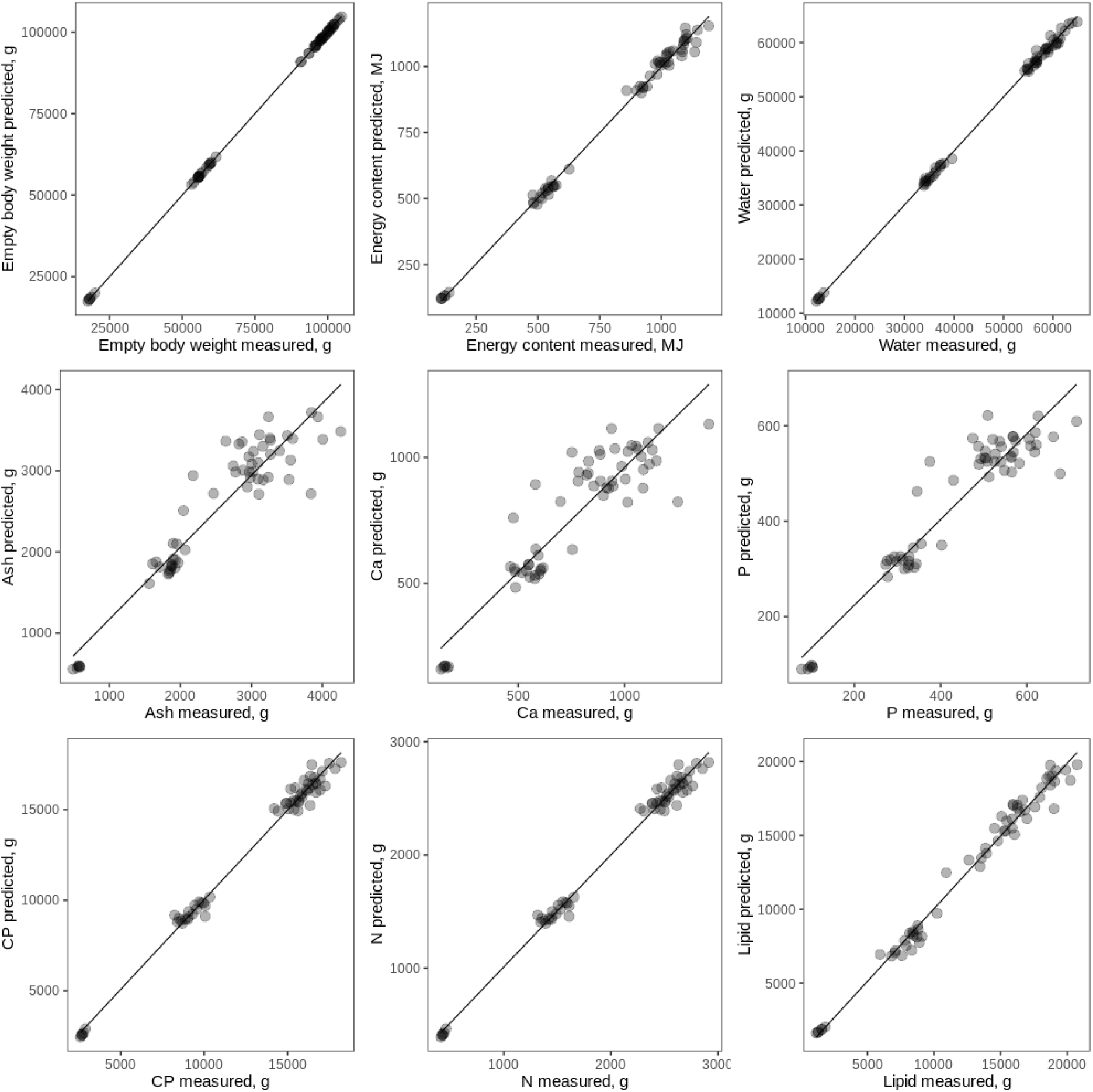
Predicted values against measured chemical values in the empty body of growing pigs. Predicted values were obtained by applying the regression equations to the DXA measurements. Dots represent individual data points.

**Figure 3.**
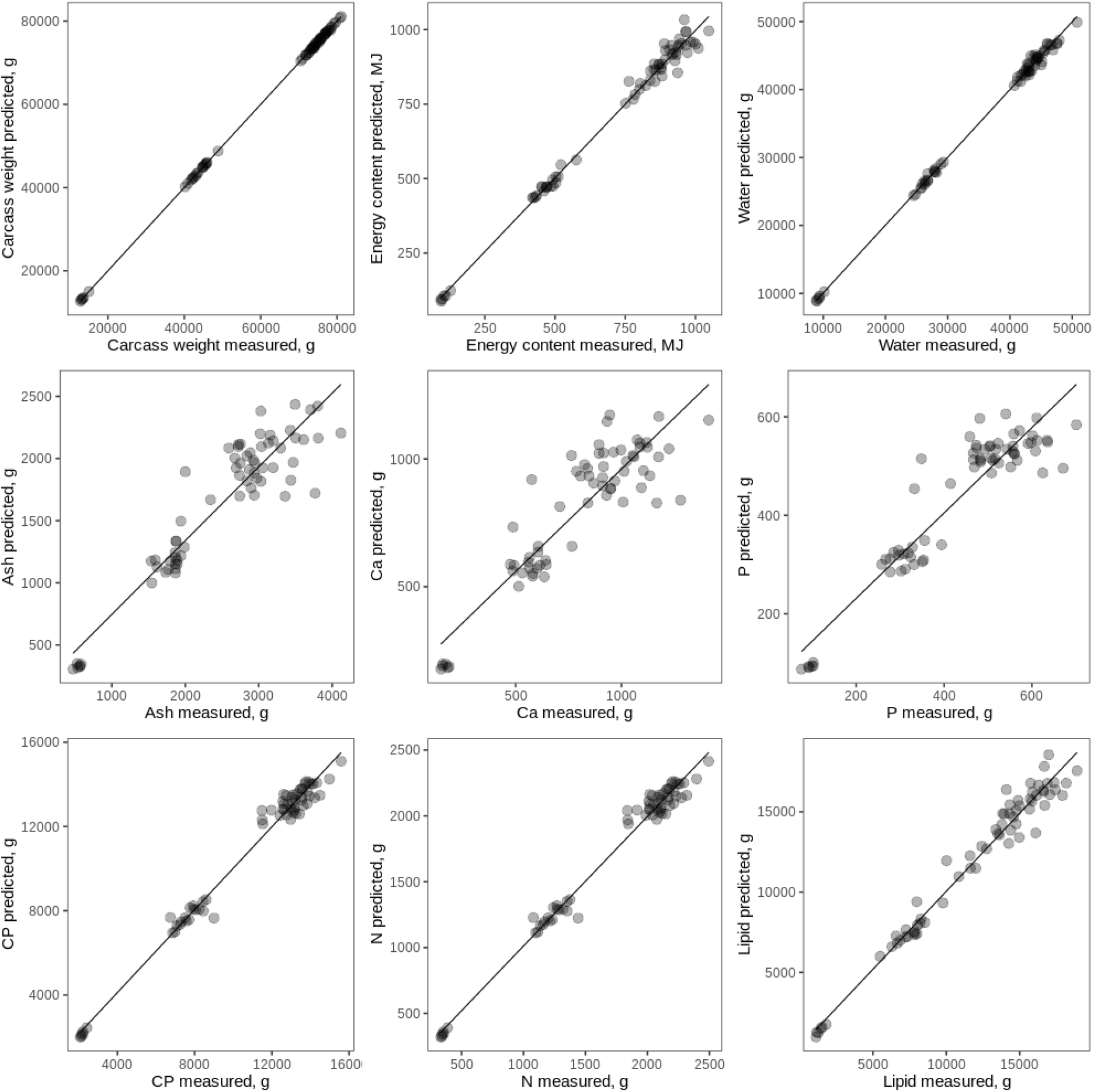
Predicted values against measured chemical values in the carcass of growing pigs. Predicted values were obtained by applying the regression equations to the DXA measurements. Dots represent individual data points.

#### CP, N, water and lipids

For energy_DXA_, CP_DXA_, N_DXA_ and water_DXA_, we obtained excellent precision and accuracy in terms of R^2^ (> 0.99) (Figure 2 and 3) and error (rCV < 3.6%) when we back-predicted the chemical values using the equations presented in Table 3. LipidDXA was slightly less precise (R^2^ > 0.97). Accuracy was better in the EB than in the carcass (rCV = 5.4% and 7%, respectively). The DXA values for fat were consistently lower than chemically measured lipid values for the pigs in the 20-kg slaughter category, which was more pronounced in the empty body than in the carcass. The same phenomenon was observed in a study comparing DXA and chemical analyses in small pigs (5 to 27 kg live weight; Mitchell et al., 1998b). It appears that DXA overestimates the fat content of pigs with a high percentage of body fat (> 20%) and underestimates the fat content in lean pigs (Mitchell et al., 1998b). Similar findings have also been reported in humans (Genton et al., 2006). Because CP in the present study was directly calculated from N, the precision and error of the prediction equations for NDXA are analogous to the ones for CP_DXA_ from lean mass. The ease with which we could predict water content from lean tissue by DXA (water_DXA_) can be related to the fact that lean tissue mass is not measured directly by DXA but is instead derived from the water content and is based on specific assumptions, such as a fixed ratio of lean tissue mass to water content (Hunter et al., 2011; but note that they used a different DXA device and software). Regional regression lines overlapped with the global one and obtained similar R^2^ and error, except for CP and N, which had reduced R^2^ (Table S2, Figures S3 and S4).

#### Minerals

Predictions of ash_DXA_, Ca_DXA_ and P_DXA_ were less precise (R^2^ between 0.80 and 0.89) and subject to major error (rCV between 12.0 and 16.5%; Table 3). Ca and P contents are certainly strongly correlated with the ash content, and one must expect that these predictions cannot be better than those of the ash content. Whereas CaDXA solely depended on BMC, PDXA also included lean mass, which is in line with the knowledge that body Ca and P are found at about 99% and 80%, respectively, in the bones (Suttle, 2010). A similar lower prediction accuracy for ash has been reported in other studies (Suster et al., 2003). The slopes of regional regressions for each target slaughter-weight category differed importantly for ash and P, but were rather similar for Ca (Table S2, Figures S3 and S4). Fitting regional regressions did not only result in reduced prediction errors, but also reduced R^2^ compared to the global regression. This suggests that it is difficult to fit reliable regional calibration models for ash, Ca and P with the present dataset, and that more data and more work is needed.

### Prediction of empty body composition at slaughter from carcass

The use of carcass DXA scans to predict the ash and CP content of the EB produced levels of accuracy and low error that were equivalent to those obtained by predicting EB ash and CP content from live scans (Table 3, Figure 4). The prediction of lipid content of the EB using carcass fat content determined by DXA was marginally less precise and had greater error (R^2^ = 0.96 vs 0.97, rCV = 8.5% vs 5.4%) than predictions from live scans (Table 3, Figure 4). With increasing BW, the regression line of the prediction models for CPDXA in the EB based on carcass scans deviated from those with the live scans. The reason for this might be that a considerable amount of lipids and CP are contained in the organs and thus missing from the carcass. Depending on the nature of the problem (e.g. when repeated measurements are not necessary) and the required degree of accuracy and prediction, invasive scanning of live animals requiring anaesthesia could, therefore, be replaced by carcass scans.

**Figure 4.**
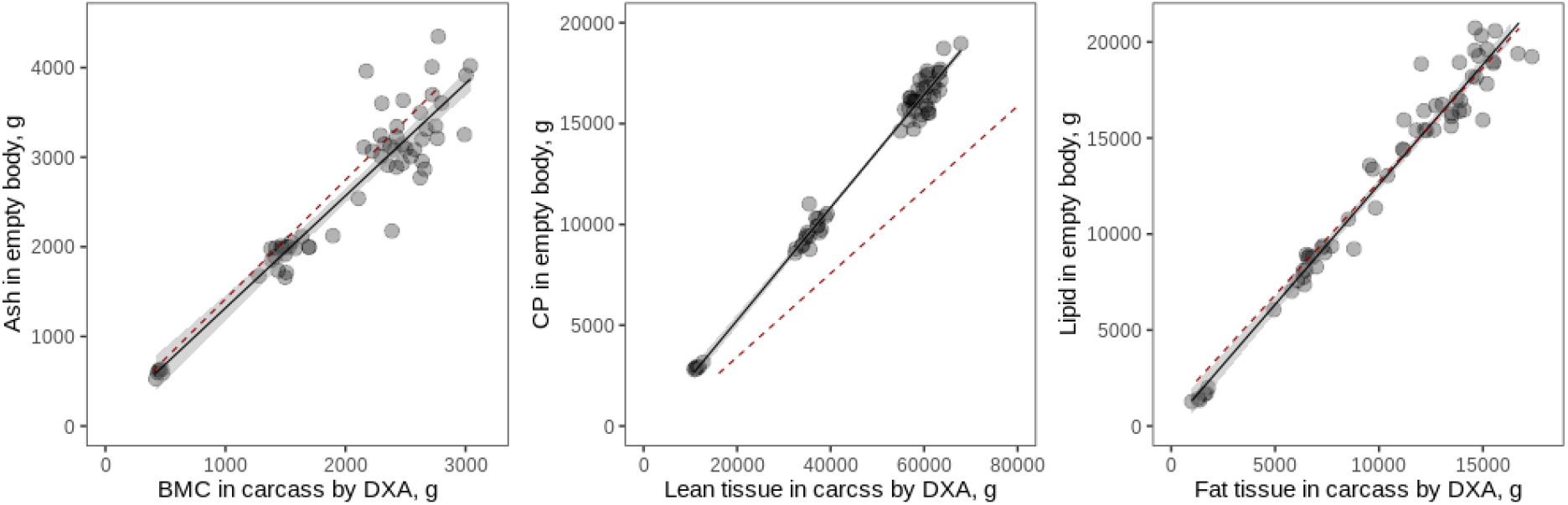
Regressions of carcass DXA values (x-axis) on empty body chemical value (y-axis) in growing pigs. The red dashed lines are regression lines from corresponding prediction equations for live DXA scans on empty body values, as presented in Table 3. For lipids, the regression line for a reduced model including only fat by DXA (instead of fat and lean by DXA) is shown.

### Comparison with published regression equations

#### Empty body

Our results are in line with published regression equations for the EB, which were built from wet chemistry analyses and live scans of pigs ranging from 5 to 97 kg live BW (Mitchell et al., 1998a), pigs with a mean live BW of 105.6 kg (Kipper et al., 2019b) and a mean warm carcass weight of 85 kg (Pomar et al., 2001, as cited in Pomar et al., 2017). Figure 5 shows the data points with regressions derived from this study and the regression lines from the literature for the EB (or live animal) for CP, water, fat and ash in the EB (or live animal). Concerning CP, the regression equations by Kipper et al. (2019b) (Figure 5, 2nd row) and Pomar et al. (2017) (Figure 5, 3rd row) fit our data quite well, whereas the one by Mitchell et al. (1998a) (Figure 5, 1st row) had a slightly worse fit. The intercepts of published equations and our study differ by 109 g to 784 g CP, which is a rather small amount given the total protein mass of the EB. Slopes were almost identical to our study (Figure 5). Using the regression by Mitchell et al. (1998a) to predict data from our study led to an overestimation of CP content in the EB throughout all weight categories. A possible explanation for the slightly worse fit of the regression equations from Mitchell et al. (1998a) than those from Kipper et al. (2019b) and Pomar et al. (2017) could be the differences in the DXA technology used. As in the present study, Kipper et al. (2019b) and Pomar et al. (2017) used a DXA scanner with a narrow fan beam, whereas Mitchell et al. (1998a) used the pencil-beam technology. It has been shown previously that different devices, and particularly different beam technologies, yield considerably variable results in terms of total BW (compared to scales), along with fat and lean mass in human subjects (Genton et al., 2002) and livestock (Scholz et al., 2015). It is conceivable that in the case of extreme proportions of body fat, such as, for example, piglets versus finishing pigs, or pigs versus small ruminants or other livestock species, the absolute values obtained by DXA could be inaccurate compared to the ‘standard’ body composition (Hunter et al., 2011). The DXA technologies and their software might differ in the way they deal with these issues, leading to the observed discrepancies. Concerning water, the published equations from Mitchell et al. (1998a) (Figure 5, 1st row) and Kipper et al. (2019b) (Figure 5, 3rd row) fit our data well (RMSE = 7.3% and 1.7%, respectively). The slopes of the lines from the data in this study and from both published equations were similar. Given the large amount of water in the body, differences in intercepts of approximately 776 g to 1 kg are not relevant, and the regression lines were very close to each other (Figure 5). However, for pigs with higher BW, Mitchell’s regressions led to a slight overestimation of water content in the EB. Concerning lipids, the regression equation by Pomar et al. (2017) had a relatively good fit to our data (Figure 5, 2nd row), with a similar slope, and it differed by 2 kg at the intercept. Thus, while this regression predicted the lipid content of pigs between 60–100 kg well, it overestimated the lipid content of piglets. Regression equations for lipids and ash by Kipper et al. (2019b) (Figure 5, 4th row) were similar to ours but consistently predicted higher values than our equation did. Kipper et al.’s (2019b) slope for lipids was similar to ours; however, their intercept was higher (by 2.7 kg), leading to a regression line almost parallel but slightly shifted upwards as compared to ours. Thus, using Kipper’s equation on our data reflected well the differences between individuals in fat content by DXA, but it overestimated the actual values.

**Figure 5.**
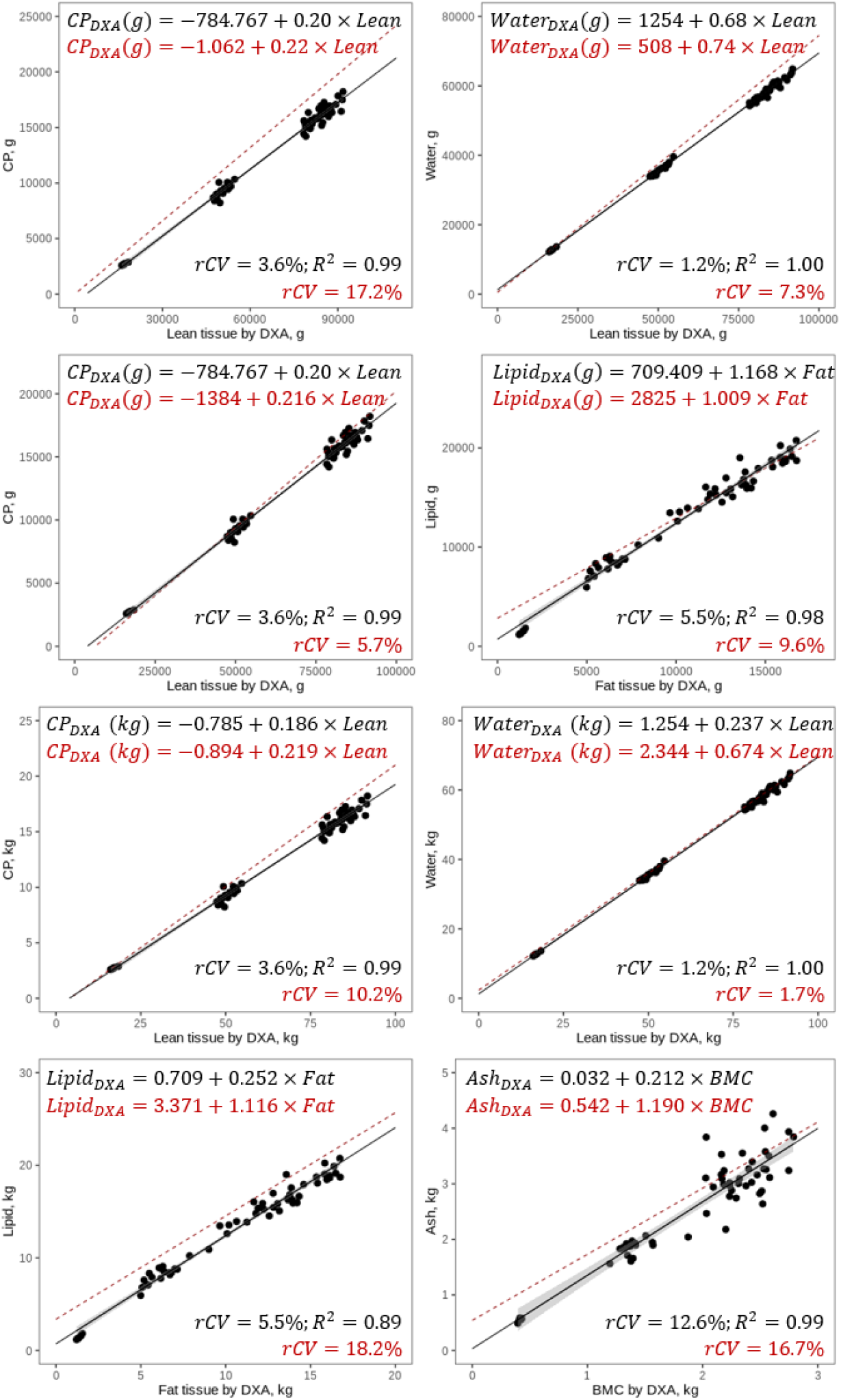
Comparison of own and published regression lines for DXA measurement on chemical value in empty body (live DXA scans) of growing pigs. The solid black lines are regression lines obtained from the data in this study, and the dashed red lines are published regression lines. Equations and rCV in black pertain to the present study; the equations in red are previously published equations. rCV in red gives the prediction error of the published equation applied to the DXA variable obtained in the present study. 1^st^ row: lean tissue by DXA on chemically determined CP and water in grams (Mitchell et al., 1998a). 2^nd^ row: lean and fat tissue by DXA on chemically determined CP and lipid in grams (Pomar et al., 2017). 3^rd^ row: Lean tissue by DXA on chemically determined CP and lean tissue by DXA on chemically determined water. 4^th^ row: fat tissue by DXA on chemically determined lipids and BMC by DXA on chemically determined ash in kilograms (Kipper et al., 2019b).

#### Carcass

The published equations for CP and water in half carcasses using absolute values (Mitchell et al., 1998c) built from wet chemistry analyses and carcass scans of pigs slaughtered at a live BW of 30 to 120 kg fit our data well, with generally lower rCVs than for the EB (Figure 6, upper row). The slopes were almost identical and the intercepts differed by 640 g for CP and less than 400 g for water. Thus, the prediction errors when using the equations by Mitchell et al. (1998c) were only slightly higher than using our own regression equations. Comparing the regression equations for percentages of lean mass and water with the ones from Mitchell et al. (2003), obtained from half carcasses with a mean weight of 43 kg, proved to be more difficult. To create our prediction equations, we calculated the percentage of lean tissue (sum of CP and water) or fat in the carcass half as the proportion of the chemical value of the sum of all chemical components (see Supplementary Material Section 6, Table S3). The equations from Mitchell et al. (2003) were dramatically different from ours, with a difference in the intercept of 13% for lean tissue and 33% for fat (Figure 6). The slopes were also very different. The resulting regression lines clearly missed our data points. This means that the equations published by Mitchell et al. (2003) are in no way suitable for predicting the chemical values for the percentage of lean mass and lipid from the present study. The reason for these striking differences could be that when the absolute values are transformed to percentages, an important amount of variation and also co-variation between DXA and wet chemistry values is lost. Consequently, fitting a regression line through data points that have such a reduced variation is difficult, and the resulting parameters of the equations might not be as robust as the ones derived from regressions on the absolute values. The pigs in this study were within a rather narrow weight range (Mitchell et al., 2003), comparable to our 100 kg BW category. Thus, these equations might not be suitable for predicting the proportion of lean tissue and lipid in the smaller pigs in our study because those differ in the proportion of lean tissue and lipids from larger pigs (Figure 6 lower row). As for lipids, Mitchell et al.’s (2003) and our regression lines seemed to converge for pigs with a lipid content of around 30%, which was outside the range in our study.

**Figure 6.**
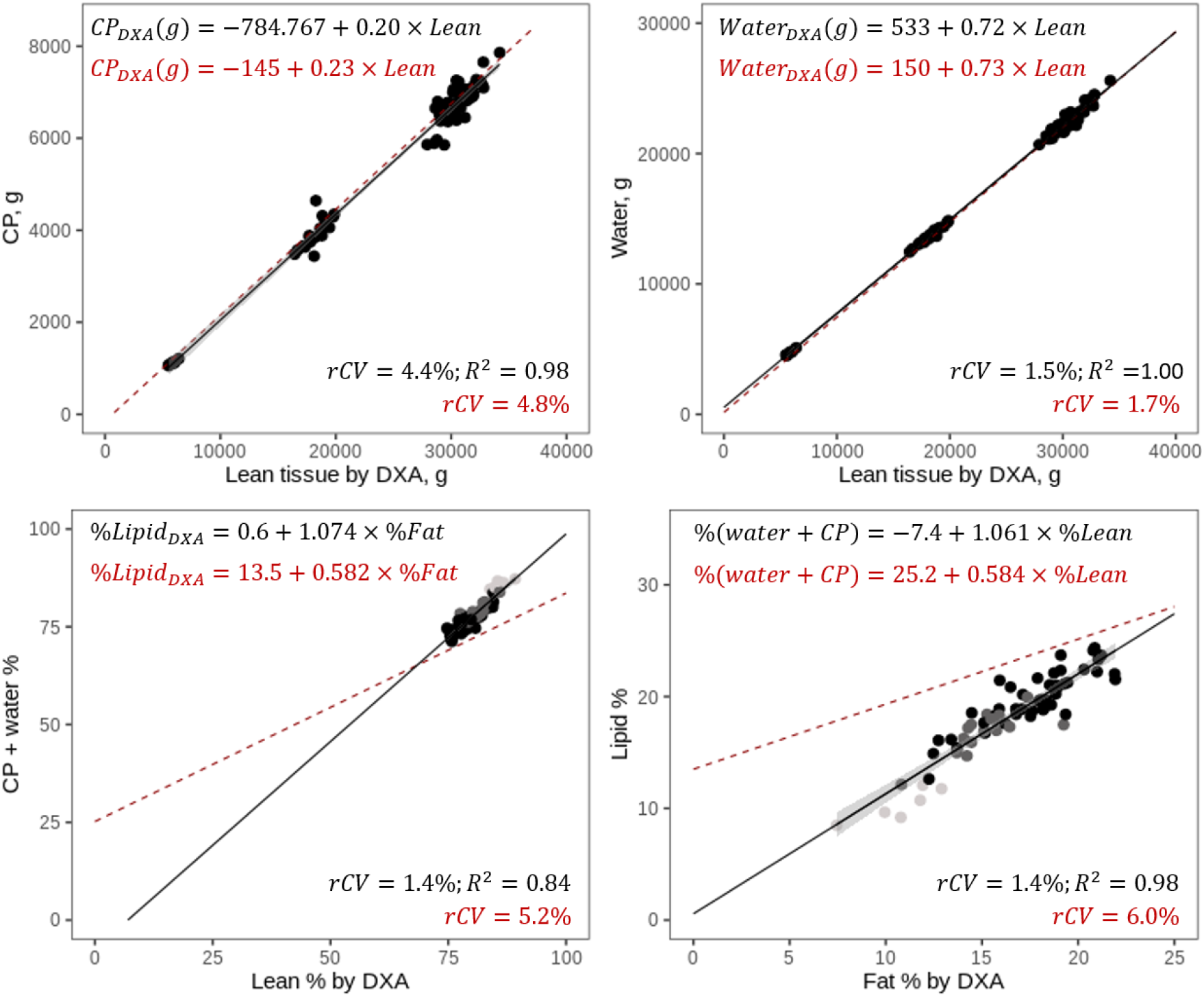
Comparison of this study and other published regression lines for DXA measurement of chemical values in carcasses (live DXA scans) of growing pigs. Solid black lines are regression lines obtained from data in this study, while dashed red lines show published regression lines. Equations, rCV and R^2^ in black pertain to the present study; the equations in red are previously published equations. rCV in red gives the prediction error of the published equation applied to the DXA variable obtained in the present study. Upper row: lean tissue DXA on chemically determined CP and water in grams (Mitchell et al., 1998c). Lower row: percentage of lean (CP + water) and fat tissue DXA on chemically determined percentage of lean and fat mass (sum of EP and water; Mitchell et al., 2003). Different categories of live weight at slaughter are illustrated by shading of the data points: light grey = 20 kg, dark grey = 60 kg, black = 100 kg.

### Conclusion

In this work, we show that although calibration regressions are usually done for each DXA device and/or working group, there is considerable agreement between the parameters of published and our own regressions in pigs. Therefore, in the near future, it might be quite realistic to create generic regression equations for body composition in pigs that yield reliable estimates for pigs of different growth stages, sexes and genetic breeds. To achieve this goal, all published results should ideally be summarized within a meta-analysis or, even better, a joint analysis. This may allow for a broader application of DXA, including for research groups that do not have the possibility to generate their own calibration equations. This endeavour would be greatly facilitated by data sharing among laboratories or, better, the existence of open data sets providing both chemical and DXA values. We illustrate the possibility of predicting the chemical content of individual nutrients such as P and Ca from DXA values. This prospect is especially important for the emerging field of precision feeding, in which the nutrient demands of single individuals have to be closely monitored. We have also shown that body composition of the EB can be inferred with rather good precision and accuracy from scanning carcass halves. Replacing the invasive, time-consuming and labour-intensive scanning of live pigs with the scanning of carcasses has great potential to improve animal welfare standards and could facilitate nutritional studies that investigate overall nutrient deposition efficiency using a large number of animals. Another promising application of DXA for measuring body composition in a relatively efficient and low-cost manner are genetic studies. Currently, the effort associated with chemical analysis certainly represents a bottleneck for those types of studies in which traits have to be measured in hundreds to thousands of individuals. DXA could be applied for high-throughput phenotyping, enabling the generation of large amounts of data for many individuals.

## Supporting information

R code for analysis

## Ethics approval

This experimental procedure was approved by the Office for Food Safety and Veterinary Affairs (2015_37_FR) and all procedures were conducted in accordance with the Ordinance on Animal Protection and the Ordinance on Animal Experimentation.

## Data accessibility

The data that support the findings of this study are publicly available in Zenodo (Kasper et al., 2020) and can be accessed at https://zenodo.org/record/3981182.

## Supplementary material

Script and codes are available online: The code used for models and the statistical analyses can be found in the Supplementary Material (accessible at https://www.biorxiv.org/content/biorxiv/early/2020/12/08/2020.09.15.286153/DC2/embed/media-2.zip?download=true).

## Acknowledgements

We are grateful to Guy Maïkoff and his team for the maintenance and slaughter of the pigs and assistance with carcass grinding and DXA scans, Sébastien Dubois and his team for carrying out the chemical analyses and Andreas Gutzwiller for assistance with DXA scans. We thank Armin Scholz and Candido Pomar for their valuable technical advice concerning the performance of DXA scans on live pigs. Florence Gondret, Arthur Francisco Araujo Fernandes, Mathieu Monziols and an anonymous reviewer provided valuable comments on a previous version of this manuscript. Version 4 of this preprint has been peer-reviewed and recommended by Peer Community In Animal Science (https://doi.org/10.24072/pci.animsci.100005).

## Appendix

Supplementary Material can be found at https://www.biorxiv.org/content/biorxiv/early/2020/12/08/2020.09.15.286153/DC1/embed/media-1.docx?download=true

