## Supplementary material for "Accuracy of predicting chemical body composition of growing pigs using dual-energy X-ray absorptiometry": R code for analysis: Kasper_etal_Rcode.Rmd

````
---
title: "Code for Accuracy of predicting chemical body composition of growing pigs by dual-energy X-ray absorptiometry"
author: "Claudia Kasper"
date: "Sept. 9, 2020"
output: word_document
---

# SETUP AND DATA LOADING

```{r initialization, results='hide', message=FALSE, warning=FALSE}
rm(list = ls(all.names = TRUE))
library(tidyverse)
library(knitr)
library(MASS)
library(car)
library(caret)

options(scipen=999) # disable scientific notation

props <- function(var, sum){
    prop <- (var/sum)*100
    return(prop)
} # calculates proportion of a component relative to the sum of all components. Input: var: specific component to compute proportion for and sum: sum of all components

CV <- function (variable) {
  100*(sd(variable)/mean(variable))
} # computes coefficient of variation of a variable

rmse <- function(measured, pred){
  return(sqrt(sum((measured - pred)^2)/(length(pred))))
} # computes square root of the sum of squared residuals divided by n. Input: measured: variable measured (with wet-chemistry methods in this case) and pred: variable predicted by regression equation

rCV <- function(measured, pred){
  paste0(round((rmse(measured, pred)*100)/mean(measured), 1), " %")
} # computes residual coefficient of variation (RMSE as % of measured value). Input: measured: variable measured (with wet-chemistry methods in this case) and pred: variable predicted by regression equation

rCV_loo <- function(measured, loo){
  paste0(round(loo$results[2]*100/mean(measured),1), " %")
} # computes residual coefficient of variation (RMSE as % of measured value) from leave-one-out procedure in package caret

round_df <- function(df, digits) {
  nums <- vapply(df, is.numeric, FUN.VALUE = logical(1))
    df[,nums] <- round(df[,nums], digits = digits)
    (df)
} # searches for numeric elements in dataframe and rounds them. Input: df: dataframe, digits: desired number of digits

looCI_1 <- function(data, lowLimit, upLimit){  
    coeffi <- data.frame(intercept = rep(NA, nrow(data)), slope1 = rep(NA, nrow(data))) # creates empty dataframe of coefficients
    for (i in 1:nrow(data)) { # loops over all cross-validation datasets, performs a linear model for each and extracts estimates of coefficients (intercept and slope) 
      a <- data[-i,]
      m <- lm(y ~ x1, data = a)  
      coeffi[i,] <- as.numeric(m$coefficients)
    }
    ciIntcpt <- sort(coeffi[,1]) # coefficient estimates are ranked and the lower und upper X% (indicated by lowLimit and upLimit)
    ciSlpe1 <- sort(coeffi[,2])
    cis <- cbind(rbind(as.data.frame(ciIntcpt[lowLimit]), ciSlpe1[lowLimit]), 
               rbind(as.data.frame(ciIntcpt[upLimit]), ciSlpe1[upLimit]))
    names(cis) <- c("lower", "upper")
    cis
} # function to construct nonparametric confidence intervals via a leave-one-out (jackknife) approach for a simple linear model (with one predictor variable). Input: data: dataframe, lowLimit/upLimit: the position in the ranked coefficient vector for the lower and the upper limit

looCI_2 <- function(data, lowLimit, upLimit){ 
    coeffi <- data.frame(intercept = rep(NA, nrow(data)), slope1 = rep(NA, nrow(data)), slope2 = rep(NA, nrow(data)))  # creates empty dataframe of coefficients
    for (i in 1:nrow(data)) { # loops over all cross-validation datasets, performs a linear model for each and extracts estimates of coefficients (intercept and slopes) 
      a <- data[-i,]
      m <- lm(y ~ x1 + x2, data = a)
      coeffi[i,] <- as.numeric(m$coefficients)
    }
    ciIntcpt <- sort(coeffi[,1]) # coefficient estimates are ranked and the lower und upper X% (indicated by lowLimit and upLimit)
    ciSlpe1 <- sort(coeffi[,2])
    ciSlpe2 <- sort(coeffi[,3])
    cis <- cbind(rbind(as.data.frame(ciIntcpt[lowLimit]), ciSlpe1[lowLimit], ciSlpe2[lowLimit]), 
               rbind(as.data.frame(ciIntcpt[upLimit]), ciSlpe1[upLimit], ciSlpe2[upLimit]))
    names(cis) <- c("lower", "upper")
    cis
} # function to construct nonparametric confidence intervals via a leave-one-out (jackknife) approach for a linear model with two predictor variables. Input: data: dataframe, lowLimit/upLimit: the position in the ranked coefficient vector for the lower and the upper limit

# Multiple plot function (copy-pasted from http://www.cookbook-r.com/Graphs/Multiple_graphs_on_one_page_(ggplot2)/)
#
# ggplot objects can be passed in ..., or to plotlist (as a list of ggplot objects)
# - cols:   Number of columns in layout
# - layout: A matrix specifying the layout. If present, 'cols' is ignored.
#
# If the layout is something like matrix(c(1,2,3,3), nrow=2, byrow=TRUE),
# then plot 1 will go in the upper left, 2 will go in the upper right, and
# 3 will go all the way across the bottom.
#

multiplot <- function(..., plotlist=NULL, file, cols=1, layout=NULL) {
  library(grid)

  # Make a list from the ... arguments and plotlist
  plots <- c(list(...), plotlist)

  numPlots = length(plots)

  # If layout is NULL, then use 'cols' to determine layout
  if (is.null(layout)) {
    # Make the panel
    # ncol: Number of columns of plots
    # nrow: Number of rows needed, calculated from # of cols
    layout <- matrix(seq(1, cols * ceiling(numPlots/cols)),
                    ncol = cols, nrow = ceiling(numPlots/cols))
  }

 if (numPlots==1) {
    print(plots[[1]])

  } else {
    # Set up the page
    grid.newpage()
    pushViewport(viewport(layout = grid.layout(nrow(layout), ncol(layout))))

    # Make each plot, in the correct location
    for (i in 1:numPlots) {
      # Get the i,j matrix positions of the regions that contain this subplot
      matchidx <- as.data.frame(which(layout == i, arr.ind = TRUE))

      print(plots[[i]], vp = viewport(layout.pos.row = matchidx$row,
                                      layout.pos.col = matchidx$col))
    }
  }
}
```

Load data sets from Zenodo 

```{r dataLoad, results='hide', message=FALSE, warning=FALSE}
eb <- read.table("https://zenodo.org/record/3981182/files/emptybody.txt?download=1", sep = "\t", header = TRUE) # load original empty body dataset (chemical contents based on empty body weight at slaughter - 3 days after DXA live scans)

carcass <- read.table("https://zenodo.org/record/3981182/files/carcass.txt?download=1", sep = "\t", header = TRUE)
```

Correct empty body weight for the 3 days between DXA scan and slaughter: 

weight definitions:

_body weight after DXA (BW_3):_ scale weight of live animal immediately after DXA scanning (3 days before slaughter)
_live body weight (BW_slaughter):_ scale weight after CO2-stunning before bleeding at slaugherhouse (3 days after DXA scans)
_empty body weight (EBW_slaughter):_ empty body weight after slaughter (BW_slaughter) minus contents of gastrointestinal tract and bladder

```{r}
eb$category <- as.factor(eb$category)
eb$ratio_ebw_bw <- eb$EBW_slaughter/eb$BW_slaughter # proportion of empty body weight : live body weight for each pig

# mean ratio for each slaughter category (20, 60 or 100 kg target weight)
ratios_ebw_bw <- as.data.frame( 
  eb %>%
  group_by(category) %>%
  summarise(
    mean = mean(ratio_ebw_bw),
    sd = sd(ratio_ebw_bw))) %>%
  ungroup()

eb$EBW_3 <- eb$BW_3 * eb$ratio_ebw_bw # estimate empty body weight at the time of DXA live scans based on the live weight at DXA and the ratio of empty body weight : live weight

correct_3d_ebw <- eb[, c("category", "EBW_slaughter", "BW_3", "chem_energy_eb", "chem_water_eb", "chem_ash_eb", "chem_Ca_eb", "chem_P_eb", "chem_CP_eb", "chem_N_eb", "chem_lipid_eb")]

correct_3d_ebw_temp <- as.data.frame(matrix(data=rep(NA, 488), ncol = 8))

# calculate the content of the empty body at the time of DXA for each chemical value in the data set by dividing the previous value for the empty body at slaughter by the empty body weight to obtain the content per g empty body weight and multiply with the product of live weight at DXA times ratio of empty body to live BW (and create new data set "emptybody")
for (j in 1:8) {
  for (i in 1:nrow(correct_3d_ebw)) {
    if (correct_3d_ebw[i, 1]=="20") {
      correct_3d_ebw_temp[i, j] <- (correct_3d_ebw[i,j+3] / correct_3d_ebw$EBW_slaughter[i]) * (correct_3d_ebw$BW_3[i]  *  ratios_ebw_bw[1,2])
    } else if (correct_3d_ebw[i,1]=="60") {
      correct_3d_ebw_temp[i, j] <- (correct_3d_ebw[i,j+3] / correct_3d_ebw$EBW_slaughter[i]) * (correct_3d_ebw$BW_3[i]  *  ratios_ebw_bw[2,2])
    } else {
      correct_3d_ebw_temp[i, j] <- (correct_3d_ebw[i,j+3] / correct_3d_ebw$EBW_slaughter[i]) * (correct_3d_ebw$BW_3[i]  *  ratios_ebw_bw[3,2])
    } 
  }
}

emptybody <- eb

emptybody[, c("chem_energy_eb", "chem_water_eb", "chem_ash_eb", "chem_Ca_eb", "chem_P_eb", "chem_CP_eb", "chem_N_eb", "chem_lipid_eb")] <- correct_3d_ebw_temp[, 1:8]
```

subsetting data to weight groups:

```{r subset}
carc_20 <- subset(carcass, subset = category == 20)
carc_60 <- subset(carcass, subset = category == 60)
carc_100 <- subset(carcass, subset = category == 100)

eb_20 <- subset(emptybody, subset = category == 20)
eb_60 <- subset(emptybody, subset = category == 60)
eb_100 <- subset(emptybody, subset = category == 100)
```

# DATA CHECKS (see Supplementary Material section 2)

### 1. Check whether sums of components in chemical analysis corresponds to empty body weight 

```{r}
sumComponents_carc <- (carcass$chem_water_carc + carcass$chem_ash_carc + carcass$chem_CP_carc + carcass$chem_lipid_carc) 
sumComponents_eb <- (emptybody$chem_water_eb + emptybody$chem_ash_eb + emptybody$chem_CP_eb + emptybody$chem_lipid_eb)
```

By computing the sum of the components, the error in empty body weight (at the time of DXA live scans) was on average `r round(mean(emptybody$emptyBW_DXA - sumComponents_eb), 2)` g $\pm$ `r round(sd(emptybody$emptyBW_DXA - sumComponents_eb), 2)` g and in carcass weight on average `r round(mean(carcass$coldCW - sumComponents_carc), 2)` g $\pm$ `r round(sd(carcass$coldCW - sumComponents_carc), 2)` g. 

### 2. Histograms of chemical body composition and coefficients of variation

```{r}
nameVector <- c("water", "ash", "CP", "lipid")
colours_eb <- c("#7e082a", "#006666", "#f6cd61", "#f37736")
colours_carc <- c("#be8394", "#7fb2b2", "#fae6b0", "#f7ad86")

df_orig_eb <- emptybody[, c("chem_water_eb", "chem_ash_eb", "chem_CP_eb", "chem_lipid_eb")]
df_orig_carc <- carcass[, c("chem_water_carc", "chem_ash_carc", "chem_CP_carc", "chem_lipid_carc")]

hist_fun <- function(df, var, colours, name){
  cvtitle <- function(prop){
    paste0("; CV = ", round(CV(prop),2), "%")
  }
  
  var <- props(var, rowSums(df))
  cv <- cvtitle(var)
  
  ggplot(df, aes(x=var)) +
    geom_histogram(binwidth=0.5, colour="black", fill=colours) +
    geom_vline(xintercept = mean(var), colour="red") +
    ylab("count") + xlab(paste0("proportion of ", name)) + 
  theme_bw(base_size = 15, base_family = "Calibri") + theme(panel.grid.major=element_blank(), panel.grid.minor=element_blank()) + 
  theme(legend.position="none") + 
  ggtitle(paste0(name,  ": mean = ", round(mean(var), 2), cv)) +
  theme(plot.title = element_text(size = 15, face = "bold"))
}

myplots_eb <- vector('list', ncol(df_orig_eb))

for (i in seq_along(df_orig_eb)) {
    myplots_eb[[i]] <- local({
        i <- i
        p1 <- hist_fun(df = df_orig_eb, var = df_orig_eb[,i], colours = colours_eb[i], name = nameVector[i])
    })
}

myplots_carc <- vector('list', ncol(df_orig_carc))

for (i in seq_along(df_orig_carc)) {
    myplots_carc[[i]] <- local({
        i <- i
        p1 <- hist_fun(df = df_orig_carc, var = df_orig_carc[,i], colours = colours_carc[i], name = nameVector[i])
    })
}
```

## Figure S1

```{r eval=FALSE, warning=FALSE, message=FALSE}
multiplot(myplots_eb[[1]], myplots_eb[[2]], myplots_eb[[3]], myplots_eb[[4]], 
          myplots_carc[[1]], myplots_carc[[2]], myplots_carc[[3]], myplots_carc[[4]],
          cols = 4)

png("FigS1.png", width = 1000, height = 660)
multiplot(myplots_eb[[1]], myplots_carc[[1]], myplots_eb[[2]], myplots_carc[[2]], myplots_eb[[3]], myplots_carc[[3]], myplots_eb[[4]], myplots_carc[[4]],
          cols = 4)
dev.off()
```

## Table 1

```{r}
stats <- c("mean", "sd", "min", "max")

table1 <- as.data.frame(matrix(c(rep(c(20, 60, 100), 13), rep(NA, 312)), ncol = 9))
names(table1) <- c("category", "mean empty body", "sd empty body", "min empty body",  "max empty body",  "mean carcass", "sd carcass",   "min carcass",  "max carcass")

by_category_eb <- emptybody[, c("category", "DXA_totalMass_eb", "DXA_BMC_eb", "DXA_lean_eb", "DXA_fat_eb", "BW_3", "chem_energy_eb", "chem_water_eb", "chem_ash_eb", "chem_Ca_eb", "chem_P_eb", "chem_CP_eb", "chem_N_eb", "chem_lipid_eb")] %>%
  group_by(category) 

for (i in 1:4){
  table1[, i+1] <- as.data.frame(by_category_eb %>%
  summarise_all(stats[i]) %>%
  pivot_longer(-category, names_to = "variable", values_to = "values") %>%
  arrange(factor(variable, levels = c("DXA_totalMass_eb", "DXA_BMC_eb", "DXA_lean_eb", "DXA_fat_eb", "BW_3", "chem_energy_eb", "chem_water_eb", "chem_ash_eb",
                                      "chem_Ca_eb", "chem_P_eb", "chem_CP_eb", "chem_N_eb", "chem_lipid_eb"))))[,3]
}

by_category_carc <- carcass[, c("category", "DXA_totalMass_carc", "DXA_BMC_carc", "DXA_lean_carc", "DXA_fat_carc", "coldCW", "chem_energy_carc", "chem_water_carc", "chem_ash_carc", "chem_Ca_carc", "chem_P_carc", "chem_CP_carc", "chem_N_carc", "chem_lipid_carc")] %>%
  group_by(category) 

for (i in 1:4){
  table1[, i+5] <- as.data.frame(by_category_carc %>%
  summarise_all(stats[i]) %>%
  pivot_longer(-category, names_to = "variable", values_to = "values") %>%
  arrange(factor(variable, levels = c("DXA_totalMass_carc", "DXA_BMC_carc", "DXA_lean_carc", "DXA_fat_carc", "coldCW", "chem_energy_carc", "chem_water_carc",
                                      "chem_ash_carc", "chem_Ca_carc", "chem_P_carc", "chem_CP_carc", "chem_N_carc", "chem_lipid_carc"))))[,3]
}

row.names(table1) <- paste0(c(rep("DXA total mass [g]",3), rep("DXA BMC [g]",3), rep("DXA lean [g]",3), rep("DXA fat [g]",3), rep("weight [g]",3), rep("chem energy [MJ]",3), rep("chem water [g]",3), rep("chem ash [g]",3), rep("chem Ca [g]",3), rep("chem P [g]",3), rep("chem CP [g]",3), rep("chem N [g]",3), rep("chem lipid [g]",3)), " ", rep(c("20", "60", "100"), 13)) 

write.table(round_df(table1), file = "table1.txt", sep = "\t")
```

## Table2 

```{r}
tests <- function(dxa, chem){
  tt <- t.test(dxa, chem)
  list <- c(mean(dxa), sd(dxa), mean(chem), sd(chem), (mean(dxa - chem)/mean(dxa))*100, tt$statistic, tt$parameter, tt$p.value)
  return(list)
}

table2 <- as.data.frame(matrix(rep(NA, 80), ncol = 8))

emptybody$chem_CPplusWater_eb <- emptybody$chem_CP_eb + emptybody$chem_water_eb
pairs_eb <- as.data.frame(matrix(as.character(c("DXA_totalMass_eb", "BW_3", "DXA_BMC_eb", "chem_ash_eb", "DXA_lean_eb", "chem_CP_eb",
                                "DXA_lean_eb", "chem_CPplusWater_eb", "DXA_fat_eb", "chem_lipid_eb")), ncol = 2, byrow = TRUE))
                  
for (i in 1:nrow(pairs_eb)) {
  table2[i,] <- tests(emptybody[, as.character(pairs_eb[i,1])], emptybody[, as.character(pairs_eb[i,2])])
}

carcass$chem_CPplusWater_carc <- carcass$chem_CP_carc + carcass$chem_water_carc
pairs_carc <- as.data.frame(matrix(as.character(c("DXA_totalMass_carc", "coldCW", "DXA_BMC_carc", "chem_ash_carc", "DXA_lean_carc", "chem_CP_carc",
                                "DXA_lean_carc", "chem_CPplusWater_carc", "DXA_fat_carc", "chem_lipid_carc")), ncol = 2, byrow = TRUE))
                  
for (i in 1:nrow(pairs_carc)) {
  table2[i+5,] <- tests(carcass[, as.character(pairs_carc[i,1])], carcass[, as.character(pairs_carc[i,2])])
}

item_chem <- c("empty body scale weight", "ash", "CP", "CP + water", "lipid", "cold carcass scale weight", "ash", "CP", "CP + water", "lipid")
table2 <- cbind(item_chem, round_df(table2[, 1:5]), round_df(table2[, 6:7], 2), round(table2[, 8], 3))
names(table2) <- c("item_chem", "mean (DXA)[g]", "sd (DXA)[g]", "mean (chem)[g]", "sd (chem)[g]", "delta%", "t", "df", "p-value")

row.names(table2) <- c("weight (empty body)", "BMC (empty body)", "lean (empty body)", "lean + water (empty body)", "fat (empty body)", 
                       "weight (full carcass)", "BMC (full carcass)", "lean (full carcass)", "lean + water (full carcass)", 
                       "fat (full carcass)")

write.table(table2, file = "Table2.txt", sep = "\t")
```

## 3. Proportion of water in fat-free mass (protein + ash + water) 

Previous publications show that it is around 75%

```{r echo=FALSE}
fatfremass_Carc <- (carcass$chem_water_carc + carcass$chem_ash_carc + carcass$chem_CP_carc) / 1000
propWFF_Carc <- (carcass$chem_water_carc/1000) / fatfremass_Carc

fatfremass_EB <- (emptybody$chem_water_eb + emptybody$chem_ash_eb + emptybody$chem_CP_eb) / 1000
propWFF_EB <- (emptybody$chem_water_eb/1000) / fatfremass_EB
```

The proportion of water in fat-free mass of the empty body is on average `r round(mean(propWFF_EB*100))` $\pm $ `r round(sd(propWFF_EB*100), 2)`% (CV = `r round(CV(propWFF_EB*100), 2)` %) and in the carcass `r round(mean(propWFF_Carc*100))`  $\pm $ `r round(sd(propWFF_Carc*100), 2)`% (CV = `r round(CV(propWFF_Carc*100), 2)` %). 

# PREDICTION EQUATIONS

## EMPTY BODY

### _A. TOTAL WEIGHT_

For annotation of code see this part (for B-C and carcass code is not annotated). Model diagnostics (supplementary material section 4) only shown for total weight

```{r BW_eb}
# model and model including treatment effect (for supplementary material section 3)
mBW_eb <- lm(BW_3 ~ DXA_totalMass_eb, data = emptybody)
mBW_eb_treat <- lm(BW_3 ~ DXA_totalMass_eb + treatment, data = emptybody)

# model diagnostics (for supplementary material section 4)
plot(mBW_eb)
outlierTest(mBW_eb) 
qqPlot(mBW_eb) 
leveragePlots(mBW_eb)
sresid <- studres(mBW_eb)
hist(sresid, freq=FALSE,  main="Distribution of Studentized Residuals")
xfit<-seq(min(sresid),max(sresid),length=40)
yfit<-dnorm(xfit)
lines(xfit, yfit) 

# data frame for calculations for table 3
df_TotWeight_EB <- data.frame(y=emptybody$BW_3, x1=emptybody$DXA_totalMass_eb, x2=NA)

# regional regressions on subsets
mBW_eb_20 <- lm(BW_3 ~ DXA_totalMass_eb, data = eb_20)
mBW_eb_60 <- lm(BW_3 ~ DXA_totalMass_eb, data = eb_60)
mBW_eb_100 <- lm(BW_3 ~ DXA_totalMass_eb, data = eb_100)
```

CI obtained by _leave-one-out procedure_:

To obtain the confidence intervals, we computed intercepts and slopes of all models by omitting one row at a time, ordered the estimates according to their value and took the 3rd and 59th and the 3rd and 66th ranked estimates for models of the empty body and the carcass, respectively, as the limits of the confidence intervals. This corresponds to a 93% CI in the empty body and a 94% CI in the carcass. 

```{r}
looCI01 <- looCI_1(df_TotWeight_EB, 3, 59)
kable(round_df(looCI01, 3))
```

### _B. ENERGY_ 

```{r energy_eb}
mENERGY_eb <- lm(chem_energy_eb ~ DXA_lean_eb + DXA_fat_eb, data = emptybody)
mENERGY_eb_treat <- lm(chem_energy_eb ~ DXA_lean_eb + DXA_fat_eb + treatment, data = emptybody)
df_Energy_EB <- data.frame(y=emptybody$chem_energy_eb, x1=emptybody$DXA_lean_eb, x2=emptybody$DXA_fat_eb)

mENERGY_eb_20 <- lm(chem_energy_eb ~ DXA_lean_eb + DXA_fat_eb, data = eb_20)
mENERGY_eb_60 <- lm(chem_energy_eb ~ DXA_lean_eb + DXA_fat_eb, data = eb_60)
mENERGY_eb_100 <- lm(chem_energy_eb ~ DXA_lean_eb + DXA_fat_eb, data = eb_100)

mENERGY_eb_s <- lm(chem_energy_eb ~ DXA_totalMass_eb, data = emptybody)
mENERGY_eb_20_s <- lm(chem_energy_eb ~ DXA_totalMass_eb, data = eb_20)
mENERGY_eb_60_s <- lm(chem_energy_eb ~ DXA_totalMass_eb, data = eb_60)
mENERGY_eb_100_s <- lm(chem_energy_eb ~ DXA_totalMass_eb, data = eb_100)
```

### _C. WATER_

```{r water_eb}
mWATER_eb <- lm(chem_water_eb ~ DXA_lean_eb, data = emptybody)
mWATER_eb_treat <- lm(chem_water_eb ~ DXA_lean_eb + treatment, data = emptybody)
df_Water_EB <- data.frame(y=emptybody$chem_water_eb, x1=emptybody$DXA_lean_eb, x2=NA)

mWATER_eb_20 <- lm(chem_water_eb ~ DXA_lean_eb, data = eb_20)
mWATER_eb_60 <- lm(chem_water_eb ~ DXA_lean_eb, data = eb_60)
mWATER_eb_100 <- lm(chem_water_eb ~ DXA_lean_eb, data = eb_100)
```

### _D. ASH_ 

```{r ash_eb}
mASH_eb <- lm(chem_ash_eb ~ DXA_BMC_eb, data = emptybody)
mASH_eb_treat <- lm(chem_ash_eb ~ DXA_BMC_eb + treatment, data = emptybody)
df_Ash_EB <- data.frame(y=emptybody$chem_ash_eb, x1=emptybody$DXA_BMC_eb, x2=NA)

mASH_eb_20 <- lm(chem_ash_eb ~ DXA_BMC_eb, data = eb_20)
mASH_eb_60 <- lm(chem_ash_eb ~ DXA_BMC_eb, data = eb_60)
mASH_eb_100 <- lm(chem_ash_eb ~ DXA_BMC_eb, data = eb_100)
```

### _E. Ca_

```{r CA_eb}
mCa_eb <- lm(chem_Ca_eb ~ DXA_BMC_eb, data = emptybody)
mCa_eb_treat <- lm(chem_Ca_eb ~ DXA_BMC_eb + treatment, data = emptybody)
df_Ca_EB <- data.frame(y=emptybody$chem_Ca_eb, x1=emptybody$DXA_BMC_eb, x2=NA)

mCa_eb_20 <- lm(chem_Ca_eb ~ DXA_BMC_eb, data = eb_20)
mCa_eb_60 <- lm(chem_Ca_eb ~ DXA_BMC_eb, data = eb_60)
mCa_eb_100 <- lm(chem_Ca_eb ~ DXA_BMC_eb, data = eb_100)
```

### _F. P_ 

```{r P_eb}
mP_eb <- lm(chem_P_eb ~ DXA_BMC_eb + DXA_lean_eb, data = emptybody)
mP_eb_treat <- lm(chem_P_eb ~ DXA_BMC_eb + DXA_lean_eb + treatment, data = emptybody)
df_P_EB <- data.frame(y=emptybody$chem_P_eb, x1=emptybody$DXA_BMC_eb, x2=emptybody$DXA_lean_eb)

mP_eb_20 <- lm(chem_P_eb ~ DXA_BMC_eb + DXA_lean_eb, data = eb_20)
mP_eb_60 <- lm(chem_P_eb ~ DXA_BMC_eb + DXA_lean_eb, data = eb_60)
mP_eb_100 <- lm(chem_P_eb ~ DXA_BMC_eb + DXA_lean_eb, data = eb_100)

mP_eb_s <- lm(chem_P_eb ~ DXA_BMC_eb, data = emptybody)
mP_eb_20_s <- lm(chem_P_eb ~ DXA_BMC_eb, data = eb_20)
mP_eb_60_s <- lm(chem_P_eb ~ DXA_BMC_eb, data = eb_60)
mP_eb_100_s <- lm(chem_P_eb ~ DXA_BMC_eb, data = eb_100)
```

### _G. CP_ 

```{r CP_eb}
mCP_eb <- lm(chem_CP_eb ~ DXA_lean_eb, data = emptybody)
mCP_eb_treat <- lm(chem_CP_eb ~ DXA_lean_eb + treatment, data = emptybody)
df_CP_EB <- data.frame(y=emptybody$chem_CP_eb, x1=emptybody$DXA_lean_eb, x2=NA)

mCP_eb_20 <- lm(chem_CP_eb ~ DXA_lean_eb, data = eb_20)
mCP_eb_60 <- lm(chem_CP_eb ~ DXA_lean_eb, data = eb_60)
mCP_eb_100 <- lm(chem_CP_eb ~ DXA_lean_eb, data = eb_100)
```

### _H. N_ 

```{r N_eb}
mN_eb <- lm(chem_N_eb ~ DXA_lean_eb, data = emptybody)
mN_eb_treat <- lm(chem_N_eb ~ DXA_lean_eb + treatment, data = emptybody)
df_N_EB <- data.frame(y=emptybody$chem_N_eb, x1=emptybody$DXA_lean_eb, x2=NA)

mN_eb_20 <- lm(chem_N_eb ~ DXA_lean_eb, data = eb_20)
mN_eb_60 <- lm(chem_N_eb ~ DXA_lean_eb, data = eb_60)
mN_eb_100 <- lm(chem_N_eb ~ DXA_lean_eb, data = eb_100)
```

### _I. LIPID_ 

```{r RLT_eb}
mLipid_eb <- lm(chem_lipid_eb ~ DXA_fat_eb + DXA_lean_eb, data = emptybody)
mLipid_eb_treat <- lm(chem_lipid_eb ~ DXA_fat_eb + DXA_lean_eb + treatment, data = emptybody)
df_Lipid_EB <- data.frame(y=emptybody$chem_lipid_eb, x1=emptybody$DXA_fat_eb, x2=emptybody$DXA_lean_eb)

mLipid_eb_20 <- lm(chem_lipid_eb ~ DXA_fat_eb + DXA_lean_eb, data = eb_20)
mLipid_eb_60 <- lm(chem_lipid_eb ~ DXA_fat_eb + DXA_lean_eb, data = eb_60)
mLipid_eb_100 <- lm(chem_lipid_eb ~ DXA_fat_eb + DXA_lean_eb, data = eb_100)

mLipid_eb_s <- lm(chem_lipid_eb ~ DXA_fat_eb, data = emptybody)
mLipid_eb_20_s <- lm(chem_lipid_eb ~ DXA_fat_eb, data = eb_20)
mLipid_eb_60_s <- lm(chem_lipid_eb ~ DXA_fat_eb, data = eb_60)
mLipid_eb_100_s <- lm(chem_lipid_eb ~ DXA_fat_eb, data = eb_100)
```


## Figure 2

define vector of indices for variables in emptybody data frame

```{r}
varList_eb <- as.data.frame(matrix(data = c(4,  15, NA, 12, 17, 18, 7, 17, NA, 8, 16, NA, 13, 16, NA, 14, 16, 17, 11, 17, NA, 9, 17, NA, 10, 18, 17), ncol = 3, byrow = TRUE))
```

create data frame for global and regional regressions

ANNOTATION OF CODE MISSING

```{r}
df_part <- function(model1, model2, model3, model4) { # for simple regression
  df <- data.frame(intercept=c(as.numeric(coefficients(model1)[1]), as.numeric(coefficients(model2)[1]),
                               as.numeric(coefficients(model3)[1]), as.numeric(coefficients(model4)[1])),
                   x1=c(as.numeric(coefficients(model1)[2]), as.numeric(coefficients(model2)[2]),
                        as.numeric(coefficients(model3)[2]), as.numeric(coefficients(model4)[2])))
  rownames(df) <- c("full data", "20 kg", "60 kg", "100 kg")
  df$x2 <- rep(NA, 4)
  df$Rsq <- c(summary(model1)$adj.r.squared, summary(model2)$adj.r.squared, summary(model3)$adj.r.squared, summary(model4)$adj.r.squared)
  return(df)
} # function that takes global model (model 1) and regional models (model 2 to 4) and returns a data frame with 

df_part_2 <- function(model1, model2, model3, model4) { # for multiple regression
  df <- data.frame(intercept=c(as.numeric(coefficients(model1)[1]), as.numeric(coefficients(model2)[1]),
                               as.numeric(coefficients(model3)[1]), as.numeric(coefficients(model4)[1])),
                   x1=c(as.numeric(coefficients(model1)[2]), as.numeric(coefficients(model2)[2]),
                        as.numeric(coefficients(model3)[2]), as.numeric(coefficients(model4)[2])),
                   x2=c(as.numeric(coefficients(model1)[3]), as.numeric(coefficients(model2)[3]),
                        as.numeric(coefficients(model3)[3]), as.numeric(coefficients(model4)[3]))                   )
  rownames(df) <- c("full data", "20 kg", "60 kg", "100 kg")
  df$Rsq <- c(summary(model1)$adj.r.squared, summary(model2)$adj.r.squared, summary(model3)$adj.r.squared, summary(model4)$adj.r.squared)
  return(df)
}
```

```{r}
eb_df <- rbind(df_part(mBW_eb, mBW_eb_20, mBW_eb_60, mBW_eb_100),df_part_2(mENERGY_eb, mENERGY_eb_20, mENERGY_eb_60, mENERGY_eb_100),
               df_part(mWATER_eb, mWATER_eb_20, mWATER_eb_60, mWATER_eb_100), df_part(mASH_eb, mASH_eb_20, mASH_eb_60, mASH_eb_100),
               df_part(mCa_eb, mCa_eb_20, mCa_eb_60, mCa_eb_100), df_part_2(mP_eb, mP_eb_20, mP_eb_60, mP_eb_100),
               df_part(mCP_eb, mCP_eb_20, mCP_eb_60, mCP_eb_100), df_part(mN_eb, mN_eb_20, mN_eb_60, mN_eb_100), 
               df_part_2(mLipid_eb, mLipid_eb_20, mLipid_eb_60, mLipid_eb_100))
```


```{r}
eb_df_full <- eb_df[seq(1, 36, 4),]
eb_df_20 <- eb_df[seq(1, 36, 4) + 1,]
eb_df_60 <- eb_df[seq(1, 36, 4) + 2,]
eb_df_100 <- eb_df[seq(1, 36, 4) + 3,]
```

add dataframe with predicted variables to original data frame

```{r}
pred_eb <- list()

for (i in 1:nrow(varList_eb)) {
  pred_eb[[i]] <- eb_df_full$intercept[i] + eb_df_full$x1[i] * emptybody[, varList_eb [i,2]]
}

# in the empty body, energy [2], phosphorus [6] and lipid [9] have 2 predictor variables

pred_eb[[2]] <- eb_df_full$intercept[2] + eb_df_full$x1[2] * emptybody[, varList_eb [2,2]] + 
  eb_df_full$x2[2] * emptybody[, varList_eb [2,3]]

pred_eb[[6]] <- eb_df_full$intercept[6] + eb_df_full$x1[6] * emptybody[, varList_eb [6,2]] + 
  eb_df_full$x2[6] * emptybody[, varList_eb [6,3]]

pred_eb[[9]] <- eb_df_full$intercept[9] + eb_df_full$x1[9] * emptybody[, varList_eb [9,2]] + 
  eb_df_full$x2[9] * emptybody[, varList_eb [9,3]]

emptybody_add <- as.data.frame(t(do.call(rbind.data.frame, pred_eb)), row.names = FALSE)
names(emptybody_add) <- c("prBW_eb", "prEnergy_bw", "prWater_eb", "prAsh_eb", "prCa_eb", "prP_eb", "prCP_eb", "prN_eb", "prLipid_eb")

emptybody <- cbind(emptybody, emptybody_add)
```

Construction of figure showing the data points and regression lines for empty body. Function is reused for Figure 3 and code is similar for Figure 3 (not annotated there).

x - measured, y - predicted (21-20)

```{r warning=FALSE, message=FALSE}
fig_Fun <- function(indata, inx, iny, xlab, ylab){
  nms <- names(indata)
  x <- nms[inx]
  y <- nms[iny]
  p <- ggplot(indata, aes(x = !!ensym(x), y = !!ensym(y))) + # see addressing x and y in aes by variable number (https://stackoverflow.com/questions/15323269/addressing-x-and-y-in-aes-by-variable-number)
    geom_point(size=4, alpha = 0.3) +
    geom_smooth(method='lm', formula= y~x, se=FALSE, color="black", linetype=1, size=.5) +
    ylab(ylab) + xlab(xlab) + 
    theme_bw(base_size = 15, base_family = "Arial") +
    theme(panel.grid.major = element_blank(), panel.grid.minor = element_blank()) + 
    theme(legend.position="none")
  p
}  # creates a figure with DXA values on x-axis and chemical values on y-axis, including raw data points and simple linear regression line with 95% CI. Input: indata: data frame, inx: vector with numbers indicating columns of explanatory variables, iny: vector with numbers indicating columns of outcome variables, xlab/ylab: vector with axes labels

ylabs <- c("Empty body weight predicted, g", "Energy content predicted, MJ", "Water predicted, g", "Ash predicted, g", "Ca predicted, g", "P predicted, g", "CP predicted, g", "N predicted, g", "Lipid predicted, g")

xlabs <- c("Empty body weight measured, g", "Energy content measured, MJ", "Water measured, g", "Ash measured, g", "Ca measured, g", "P measured, g", "CP measured, g", "N measured, g", "Lipid measured, g")

inx <- c(4, 12, 7, 8, 13, 14, 11, 9, 10)
iny <- 21:29

plots_eb <- vector('list', 9)

# loop over all variables for the empty body to create figure for each and store it in list object
for (i in 1:9) {
  plots_eb[[i]] <- fig_Fun(indata=emptybody, inx=inx[i], iny=iny[i], xlab = xlabs[i], ylab = ylabs[i])
}
```

print 3x3 plot for all nine variables for the empty body

```{r eval=FALSE, warning=FALSE, message=FALSE}
multiplot(plots_eb[[1]], plots_eb[[4]], plots_eb[[7]], plots_eb[[2]], plots_eb[[5]], plots_eb[[8]], plots_eb[[3]], plots_eb[[6]], plots_eb[[9]], cols = 3)

png("Fig2.png", width = 1000, height = 1000)
multiplot(plots_eb[[1]], plots_eb[[4]], plots_eb[[7]], plots_eb[[2]], plots_eb[[5]], plots_eb[[8]], plots_eb[[3]], plots_eb[[6]], plots_eb[[9]], cols = 3)
dev.off()
```

## CARCASS

### _A. TOTAL WEIGHT_

```{r BW_carc}
mBW_carc <- lm(coldCW ~ DXA_totalMass_carc, data = carcass)
mBW_carc_treat <- lm(coldCW ~ DXA_totalMass_carc + treatment, data = carcass)
df_TotWeight_carc <- data.frame(y=carcass$coldCW, x1=carcass$DXA_totalMass_carc, x2=NA)

mBW_carc_20 <- lm(coldCW ~ DXA_totalMass_carc, data = carc_20)
mBW_carc_60 <- lm(coldCW ~ DXA_totalMass_carc, data = carc_60)
mBW_carc_100 <- lm(coldCW ~ DXA_totalMass_carc, data = carc_100)
```

### _B. ENERGY_ 

```{r energy_carc}
mENERGY_carc <- lm(chem_energy_carc ~ DXA_lean_carc + DXA_fat_carc, data = carcass)
mENERGY_carc_treat <- lm(chem_energy_carc ~ DXA_lean_carc + DXA_fat_carc + treatment, data = carcass)
df_Energy_carc <- data.frame(y=carcass$chem_energy_carc, x1=carcass$DXA_lean_carc, x2=carcass$DXA_fat_carc)

mENERGY_carc_20 <- lm(chem_energy_carc ~ DXA_lean_carc + DXA_fat_carc, data = carc_20)
mENERGY_carc_60 <- lm(chem_energy_carc ~ DXA_lean_carc + DXA_fat_carc, data = carc_60)
mENERGY_carc_100 <- lm(chem_energy_carc ~ DXA_lean_carc + DXA_fat_carc, data = carc_100)

mENERGY_carc_s <- lm(chem_energy_carc ~ DXA_totalMass_carc, data = carcass)
mENERGY_carc_20_s <- lm(chem_energy_carc ~ DXA_totalMass_carc, data = carc_20)
mENERGY_carc_60_s <- lm(chem_energy_carc ~ DXA_totalMass_carc, data = carc_60)
mENERGY_carc_100_s <- lm(chem_energy_carc ~ DXA_totalMass_carc, data = carc_100)
```

### _C. WATER_

```{r water_carc}
mWATER_carc <- lm(chem_water_carc ~ DXA_lean_carc, data = carcass)
mWATER_carc_treat <- lm(chem_water_carc ~ DXA_lean_carc + treatment, data = carcass)
df_Water_carc <- data.frame(y=carcass$chem_water_carc, x1=carcass$DXA_lean_carc, x2=NA)

mWATER_carc_20 <- lm(chem_water_carc ~ DXA_lean_carc, data = carc_20)
mWATER_carc_60 <- lm(chem_water_carc ~ DXA_lean_carc, data = carc_60)
mWATER_carc_100 <- lm(chem_water_carc ~ DXA_lean_carc, data = carc_100)
```

### _D. ASH_ 

```{r ash_carc}
mASH_carc <- lm(chem_ash_carc ~ DXA_BMC_carc + DXA_lean_carc, data = carcass)
mASH_carc_treat <- lm(chem_ash_carc ~ DXA_BMC_carc + DXA_lean_carc + treatment, data = carcass)
df_Ash_carc <- data.frame(y=carcass$chem_ash_carc, x1=carcass$DXA_BMC_carc, x2=carcass$DXA_lean_carc)

mASH_carc_20 <- lm(chem_ash_carc ~ DXA_BMC_carc + DXA_lean_carc, data = carc_20)
mASH_carc_60 <- lm(chem_ash_carc ~ DXA_BMC_carc + DXA_lean_carc, data = carc_60)
mASH_carc_100 <- lm(chem_ash_carc ~ DXA_BMC_carc + DXA_lean_carc, data = carc_100)

mASH_carc_s <- lm(chem_ash_carc ~ DXA_BMC_carc, data = carcass)
mASH_carc_20_s <- lm(chem_ash_carc ~ DXA_BMC_carc, data = carc_20)
mASH_carc_60_s <- lm(chem_ash_carc ~ DXA_BMC_carc, data = carc_60)
mASH_carc_100_s <- lm(chem_ash_carc ~ DXA_BMC_carc, data = carc_100)
```

### _E. Ca_ 

```{r CA_carc}
mCa_carc <- lm(chem_Ca_carc ~ DXA_BMC_carc, data = carcass)
mCa_carc_treat <- lm(chem_Ca_carc ~ DXA_BMC_carc + treatment, data = carcass)
df_Ca_carc <- data.frame(y=carcass$chem_Ca_carc, x1=carcass$DXA_BMC_carc, x2=NA)

mCa_carc_20 <- lm(chem_Ca_carc ~ DXA_BMC_carc, data = carc_20)
mCa_carc_60 <- lm(chem_Ca_carc ~ DXA_BMC_carc, data = carc_60)
mCa_carc_100 <- lm(chem_Ca_carc ~ DXA_BMC_carc, data = carc_100)
```

### _F. P_ 

```{r P_carc}
mP_carc <- lm(chem_P_carc ~ DXA_BMC_carc + DXA_lean_carc, data = carcass)
mP_carc_treat <- lm(chem_P_carc ~ DXA_BMC_carc + DXA_lean_carc + treatment, data = carcass)
df_P_carc <- data.frame(y=carcass$chem_P_carc, x1=carcass$DXA_BMC_carc, x2=carcass$DXA_lean_carc)

mP_carc_20 <- lm(chem_P_carc ~ DXA_BMC_carc + DXA_lean_carc, data = carc_20)
mP_carc_60 <- lm(chem_P_carc ~ DXA_BMC_carc + DXA_lean_carc, data = carc_60)
mP_carc_100 <- lm(chem_P_carc ~ DXA_BMC_carc + DXA_lean_carc, data = carc_100)

mP_carc_s <- lm(chem_P_carc ~ DXA_BMC_carc, data = carcass)
mP_carc_20_s <- lm(chem_P_carc ~ DXA_BMC_carc, data = carc_20)
mP_carc_60_s <- lm(chem_P_carc ~ DXA_BMC_carc, data = carc_60)
mP_carc_100_s <- lm(chem_P_carc ~ DXA_BMC_carc, data = carc_100)
```

### _G. CP_ 

```{r CP_carc}
mCP_carc <- lm(chem_CP_carc ~ DXA_lean_carc, data = carcass)
mCP_carc_treat <- lm(chem_CP_carc ~ DXA_lean_carc + treatment, data = carcass)
df_CP_carc <- data.frame(y=carcass$chem_CP_carc, x1=carcass$DXA_lean_carc, x2=NA)

mCP_carc_20 <- lm(chem_CP_carc ~ DXA_lean_carc, data = carc_20)
mCP_carc_60 <- lm(chem_CP_carc ~ DXA_lean_carc, data = carc_60)
mCP_carc_100 <- lm(chem_CP_carc ~ DXA_lean_carc, data = carc_100)
```

### _H. N_ 

```{r N_carc}
mN_carc <- lm(chem_N_carc ~ DXA_lean_carc, data = carcass)
mN_carc_treat <- lm(chem_N_carc ~ DXA_lean_carc + treatment, data = carcass)
df_N_carc <- data.frame(y=carcass$chem_N_carc, x1=carcass$DXA_lean_carc, x2=NA)

mN_carc_20 <- lm(chem_N_carc ~ DXA_lean_carc, data = carc_20)
mN_carc_60 <- lm(chem_N_carc ~ DXA_lean_carc, data = carc_60)
mN_carc_100 <- lm(chem_N_carc ~ DXA_lean_carc, data = carc_100)
```

### _I. LIPID_ 

```{r RLT_carc}
mLipid_carc <- lm(chem_lipid_carc ~ DXA_fat_carc + DXA_lean_carc, data = carcass)
mLipid_carc_treat <- lm(chem_lipid_carc ~ DXA_fat_carc + DXA_lean_carc + treatment, data = carcass)
df_Lipid_carc <- data.frame(y=carcass$chem_lipid_carc, x1=carcass$DXA_fat_carc, x2=carcass$DXA_lean_carc)

mLipid_carc_20 <- lm(chem_lipid_carc ~ DXA_fat_carc + DXA_lean_carc, data = carc_20)
mLipid_carc_60 <- lm(chem_lipid_carc ~ DXA_fat_carc + DXA_lean_carc, data = carc_60)
mLipid_carc_100 <- lm(chem_lipid_carc ~ DXA_fat_carc + DXA_lean_carc, data = carc_100)

mLipid_carc_s <- lm(chem_lipid_carc ~ DXA_fat_carc, data = carcass)
mLipid_carc_20_s <- lm(chem_lipid_carc ~ DXA_fat_carc, data = carc_20)
mLipid_carc_60_s <- lm(chem_lipid_carc ~ DXA_fat_carc, data = carc_60)
mLipid_carc_100_s <- lm(chem_lipid_carc ~ DXA_fat_carc, data = carc_100)
```


## Figure 3

define vector of indices for variables in emptybody data frame

```{r}
varList_carc <- as.data.frame(matrix(data = c(5,  22, NA, 11, 24, 25, 6, 24, NA, 7, 23, 24, 12, 23, NA, 13, 23, 24, 10, 24, NA, 8, 24, NA, 9, 25, 24), ncol = 3, byrow = TRUE))
```

create data frame for global and regional regressions

```{r}
carc_df <- rbind(df_part(mBW_carc, mBW_carc_20, mBW_carc_60, mBW_carc_100), 
                 df_part_2(mENERGY_carc, mENERGY_carc_20, mENERGY_carc_60, mENERGY_carc_100),
                 df_part(mWATER_carc, mWATER_carc_20, mWATER_carc_60, mWATER_carc_100), 
                 df_part_2(mASH_carc, mASH_carc_20, mASH_carc_60, mASH_carc_100),
                 df_part(mCa_carc, mCa_carc_20, mCa_carc_60, mCa_carc_100), 
                 df_part_2(mP_carc, mP_carc_20, mP_carc_60, mP_carc_100),
                 df_part(mCP_carc, mCP_carc_20, mCP_carc_60, mCP_carc_100), 
                 df_part(mN_carc, mN_carc_20, mN_carc_60, mN_carc_100), 
                 df_part_2(mLipid_carc, mLipid_carc_20, mLipid_carc_60, mLipid_carc_100))
```

```{r}
carc_df_full <- carc_df[seq(1, 36, 4),]
carc_df_20 <- carc_df[seq(1, 36, 4) + 1,]
carc_df_60 <- carc_df[seq(1, 36, 4) + 2,]
carc_df_100 <- carc_df[seq(1, 36, 4) + 3,]
```

add dataframe with predicted variables to original data frame

```{r}
pred_carc <- list()

for (i in 1:nrow(varList_carc)) {
  pred_carc[[i]] <- carc_df_full$intercept[i] + carc_df_full$x1[i] * carcass[, varList_carc [i,2]]
}

# in the empty body, energy [2], phosphorus [6] and lipid [9] have 2 predictor variables

pred_carc[[2]] <- carc_df_full$intercept[2] + carc_df_full$x1[2] * carcass[, varList_carc [2,2]] + 
  carc_df_full$x2[2] * carcass[, varList_carc [2,3]]

pred_carc[[6]] <- carc_df_full$intercept[6] + carc_df_full$x1[6] * carcass[, varList_carc [6,2]] + 
  carc_df_full$x2[6] * carcass[, varList_carc [6,3]]

pred_carc[[9]] <- carc_df_full$intercept[9] + carc_df_full$x1[9] * carcass[, varList_carc [9,2]] + 
  carc_df_full$x2[9] * carcass[, varList_carc [9,3]]

carcass_add <- as.data.frame(t(do.call(rbind.data.frame, pred_carc)), row.names = FALSE)
names(carcass_add) <- c("prBW_carc", "prEnergy_bw", "prWater_carc", "prAsh_carc", "prCa_carc", "prP_carc", "prCP_carc", "prN_carc", "prLipid_carc")

carcass <- cbind(carcass, carcass_add)
```

Construction of figure showing the data points and regression lines for empty body

x - measured, y - predicted (32-40)

```{r warning=FALSE, message=FALSE}
rm(ylabs, xlabs)

ylabs <- c("Carcass weight predicted, g", "Energy content predicted, MJ", "Water predicted, g", "Ash predicted, g", "Ca predicted, g", "P predicted, g", "CP predicted, g", "N predicted, g", "Lipid predicted, g")

xlabs <- c("Carcass weight measured, g", "Energy content measured, MJ", "Water measured, g", "Ash measured, g", "Ca measured, g", "P measured, g", "CP measured, g", "N measured, g", "Lipid measured, g")

rm(inx, iny)

inx <- c(5, 11, 6, 7, 12, 13, 10, 8, 9)
iny <- 32:40
plots_carc <- vector('list', 9)

for (i in 1:9) {
  plots_carc[[i]] <- fig_Fun(indata=carcass, inx=inx[i], iny=iny[i], xlab = xlabs[i], ylab = ylabs[i])
}
```

```{r eval=FALSE, warning=FALSE, message=FALSE}
multiplot(plots_carc[[1]], plots_carc[[4]], plots_carc[[7]], plots_carc[[2]], plots_carc[[5]], plots_carc[[8]], plots_carc[[3]], plots_carc[[6]], plots_carc[[9]], cols = 3)

png("Fig3.png", width = 1000, height = 1000)
multiplot(plots_carc[[1]], plots_carc[[4]], plots_carc[[7]], plots_carc[[2]], plots_carc[[5]], plots_carc[[8]], plots_carc[[3]], plots_carc[[6]], plots_carc[[9]], cols = 3)
dev.off()
```

## EB from CARCASS

Here the original empty body data set is used

### _J. EB ASH ~ Carcass DXA_BMC_

```{r}
c1 <- carcass[which(carcass$ID %in% eb$ID),]

#append values for carcass to dataset for empty body at slaughter
eb$DXA_BMC_carc <- c1$DXA_BMC_carc
eb$DXA_lean_carc <- c1$DXA_lean_carc
eb$DXA_fat_carc <- c1$DXA_fat_carc

mAsh_ceb <- lm(chem_ash_eb ~ DXA_BMC_carc, data = eb)
df_Ash_ceb <- data.frame(y=eb$chem_ash_eb, x1=eb$DXA_BMC_carc, x2=NA)
```

### _K. EB CP ~ Carcass DXALean_

```{r}
mCP_ceb <- lm(chem_CP_eb ~ DXA_lean_carc, data = eb)
df_CP_ceb <- data.frame(y=eb$chem_CP_eb, x1=eb$DXA_lean_carc, x2=NA)
```

### _L. EB LIPID ~ Carcass DXAFat_

```{r}
mLipid_ceb <- lm(chem_lipid_eb ~ DXA_fat_carc + DXA_lean_carc, data = eb)
df_Lipid_ceb <- data.frame(y=eb$chem_lipid_eb, x1=eb$DXA_fat_carc, x2=eb$DXA_lean_carc)
```


## Figure 4

In addition to the regression lines of carcass DXA on empty body chemical values, this figure contains simple linear regression lines of empty body DXA on empty body chemical values (darkred, dashed line) 

```{r warning=FALSE, message=FALSE}
p1a <- ggplot(eb, aes(x = DXA_BMC_carc, y = chem_ash_eb)) +
  geom_point(size=4, alpha = 0.3) +
  geom_smooth(method='lm', formula= y~x, color="black", linetype=1, size=.5) +
  geom_smooth(aes(x = DXA_BMC_eb, y = chem_ash_eb), method='lm', formula= y~x,  # add regression line from empty body model
              color="darkred", linetype=2, size=.5, se=FALSE) +
  ylim(0, 4500) + xlim(0, 3200) +
  ylab("Ash in empty body, g") + xlab("BMC in carcass by DXA, g") +
  theme_bw(base_size = 15, base_family = "Arial") +
  theme(panel.grid.major = element_blank(), panel.grid.minor = element_blank()) + 
  theme(legend.position="none")

p2a <- ggplot(eb, aes(x = DXA_lean_carc, y = chem_CP_eb)) +
  geom_point(size=4, alpha = 0.3) +
  geom_smooth(method='lm', formula= y~x, color="black", linetype=1, size=.5) +
  geom_smooth(aes(x = DXA_lean_eb, y = chem_CP_eb), method='lm', formula= y~x, 
              color="darkred", linetype=2, size=.5, se=FALSE) +
  ylim(0, 20000) + xlim(0, 80000) +
  ylab("CP in empty body, g") + xlab("Lean tissue in carcss by DXA, g") +
  theme_bw(base_size = 15, base_family = "Arial") +
  theme(panel.grid.major = element_blank(), panel.grid.minor = element_blank()) + 
  theme(legend.position="none")

p3a <- ggplot(eb, aes(x = DXA_fat_carc, y = chem_lipid_eb)) +
  geom_point(size=4, alpha = 0.3) +
  geom_smooth(method='lm', formula= y~x, color="black", linetype=1, size=.5) +
  geom_smooth(aes(x = DXA_fat_eb, y = chem_lipid_eb), method='lm', formula= y~x, 
              color="darkred", linetype=2, size=.5, se=FALSE) +
  ylim(0, 21000) + xlim(0, 18000) +
  ylab("Lipid in empty body, g") + xlab("Fat tissue in carcass by DXA, g") +
  theme_bw(base_size = 15, base_family = "Arial") +
  theme(panel.grid.major = element_blank(), panel.grid.minor = element_blank()) + 
  theme(legend.position="none")
```

```{r eval=FALSE, warning=FALSE, message=FALSE}
multiplot(p1a, p2a, p3a, cols = 3)

png("Fig4.png", width = 1000, height = 330)
multiplot(p1a, p2a, p3a, cols = 3)
dev.off()
```

## Table 3

### Calculations for Table 3

```{r}
df_list_carcEB <- list(df_Ash_ceb, df_CP_ceb, df_Lipid_ceb)
loos_carcEB <- list()

for (i in 1:length(df_list_carcEB)) {
  if (unique(is.na(df_list_carcEB[[i]]$x2)) == TRUE) {
    loos_carcEB[[i]] <- train(y ~ x1, method = "lm", data = df_list_carcEB[[i]], trControl = trainControl(method = "LOOCV"))
  } else {
    loos_carcEB[[i]] <- train(y ~ x1 + x2, method = "lm", data = df_list_carcEB[[i]], trControl = trainControl(method = "LOOCV"))
  }
}

looCIs_carcEB <- list()
for (i in 1:length(df_list_carcEB)) {
  if (unique(is.na(df_list_carcEB[[i]]$x2)) == TRUE) {
    looCIs_carcEB[[i]] <- looCI_1(df_list_carcEB[[i]], 3, 59)
  } else {
    looCIs_carcEB[[i]] <- looCI_2(df_list_carcEB[[i]], 3, 59)
  }
}
```

Leave-one-out cross validation and CIs for empty body and carcass

```{r}
df_list_eb <- list(df_TotWeight_EB, df_Energy_EB, df_Water_EB, df_Ash_EB, df_Ca_EB, df_P_EB, df_CP_EB, df_N_EB, df_Lipid_EB)
loos_eb <- list()

for (i in 1:length(df_list_eb)) {
  if (unique(is.na(df_list_eb[[i]]$x2)) == TRUE) {
    loos_eb[[i]] <- train(y ~ x1, method = "lm", data = df_list_eb[[i]], trControl = trainControl(method = "LOOCV"))
  } else {
    loos_eb[[i]] <- train(y ~ x1 + x2, method = "lm", data = df_list_eb[[i]], trControl = trainControl(method = "LOOCV"))
  }
}

looCIs_eb <- list()
for (i in 1:length(df_list_eb)) {
  if (unique(is.na(df_list_eb[[i]]$x2)) == TRUE) {
    looCIs_eb[[i]] <- looCI_1(df_list_eb[[i]], 3, 59)
  } else {
    looCIs_eb[[i]] <- looCI_2(df_list_eb[[i]], 3, 59)
  }
}
```

```{r}
df_list_carc <- list(df_TotWeight_carc, df_Energy_carc, df_Water_carc, df_Ash_carc, df_Ca_carc, df_P_carc, df_CP_carc, df_N_carc, df_Lipid_carc)
loos_carc <- list()

for (i in 1:length(df_list_carc)) {
  if (unique(is.na(df_list_carc[[i]]$x2)) == TRUE) {
    loos_carc[[i]] <- train(y ~ x1, method = "lm", data = df_list_carc[[i]], trControl = trainControl(method = "LOOCV"))
  } else {
    loos_carc[[i]] <- train(y ~ x1 + x2, method = "lm", data = df_list_carc[[i]], trControl = trainControl(method = "LOOCV"))
  }
}

looCIs_carc <- list()
for (i in 1:length(df_list_carc)) {
  if (unique(is.na(df_list_carc[[i]]$x2)) == TRUE) {
    looCIs_carc[[i]] <- looCI_1(df_list_carc[[i]], 3, 66)
  } else {
    looCIs_carc[[i]] <- looCI_2(df_list_carc[[i]], 3, 66)
  }
}
```

### Table 3

```{r}
model_names_eb = c("mBW_eb","mENERGY_eb", "mWATER_eb", "mASH_eb", "mCa_eb", "mP_eb", "mCP_eb", "mN_eb", "mLipid_eb")
modelList_eb = lapply(model_names_eb, get)
names(modelList_eb) = model_names_eb

model_names_carc = c("mBW_carc","mENERGY_carc", "mWATER_carc", "mASH_carc", "mCa_carc", "mP_carc", "mCP_carc", "mN_carc", "mLipid_carc")
modelList_carc = lapply(model_names_carc, get)
names(modelList_carc) = model_names_carc

model_names_carcEB = c("mAsh_ceb","mCP_ceb", "mLipid_ceb")
modelList_carcEB = lapply(model_names_carcEB, get)
names(modelList_carcEB) = model_names_carcEB

table3 <- as.data.frame(matrix(rep(NA, 300), ncol = 6))
names(table3) <- c("termDXA", "estimate", "P", "Rsq", "RMSE", "rCV")
table3$termDXA <- c("Intercept", "total mass", "Intercept", "lean", "fat", "Intercept", "lean", "Intercept", "BMC", "Intercept", "BMC", "Intercept", "BMC", "lean", 
           "Intercept", "lean", "Intercept", "lean", "Intercept", "fat", "lean", 
           
           "Intercept", "total mass", "Intercept", "lean", "fat", "Intercept", "lean", "Intercept", "BMC", "lean", "Intercept", "BMC", "Intercept", "BMC", "lean", 
           "Intercept", "lean", "Intercept", "lean", "Intercept", "fat", "lean", 
           
           "Intercept", "BMC carcass", "Intercept", "lean carcass", "Intercept", "fat carcass", "lean carcass")

# estimates

estim <- function(model, CI) {
  est <- NULL
  est[1] <- paste0(round(summary(model)$coefficients[1,1], 3), " (", round(CI$lower[1], 3), ", ", round(CI$upper[1], 3), ")")
  est[2] <- paste0(round(summary(model)$coefficients[2,1], 3), " (", round(CI$lower[2], 3), ", ", round(CI$upper[2], 3), ")")
  if (is.na(model$coefficients[3]) == TRUE) {
    est[3] <- NA
  } else {
    est[3] <- paste0(round(summary(model)$coefficients[3,1], 3), " (", round(CI$lower[3], 3), ", ", round(CI$upper[3], 3), ")")
  }
  return (est)
}

estimates_eb <- as.data.frame(matrix(rep(NA, 27), ncol = 3))

for (i in 1:length(modelList_eb)) {
  estimates_eb[i,] <- estim(modelList_eb[[i]], looCIs_eb[[i]]) 
}

estimates_eb <- as.vector(t(estimates_eb))
table3$estimate[1:21] <- estimates_eb[!is.na(estimates_eb)]

estimates_carc <- as.data.frame(matrix(rep(NA, 27), ncol = 3))

for (i in 1:length(modelList_carc)) {
  estimates_carc[i,] <- estim(modelList_carc[[i]], looCIs_carc[[i]]) 
}

estimates_carc <- as.vector(t(estimates_carc))
table3$estimate[22:43] <- estimates_carc[!is.na(estimates_carc)]

estimates_carcEB <- as.data.frame(matrix(rep(NA, 9), ncol = 3))

for (i in 1:length(modelList_carcEB)) {
  estimates_carcEB[i,] <- estim(modelList_carcEB[[i]], looCIs_carcEB[[i]]) 
}

estimates_carcEB <- as.vector(t(estimates_carcEB))
table3$estimate[44:50] <- estimates_carcEB[!is.na(estimates_carcEB)]

# p-values

p_fun <- function(model) {
  p <- NULL
  p[1] <- round(summary(model)$coefficients[1,4], 3)
  p[2] <- round(summary(model)$coefficients[2,4], 3)
  if (is.na(model$coefficients[3]) == TRUE) {
    p[3] <- NA
  } else {
    p[3] <- round(summary(model)$coefficients[3,4], 3)
  }
  return (p)
}

pvals_eb <-  as.data.frame(matrix(rep(NA, 27), ncol = 3))

for (i in 1:length(modelList_eb)) {
  pvals_eb[i, ] <- p_fun(modelList_eb[[i]])
}

pvals_eb <- as.vector(t(pvals_eb))
pvals_eb[pvals_eb == 0] <- "< 0.001"
table3$P[1:21] <- pvals_eb[!is.na(pvals_eb)]

pvals_carc <-  as.data.frame(matrix(rep(NA, 27), ncol = 3))

for (i in 1:length(modelList_carc)) {
  pvals_carc[i, ] <- p_fun(modelList_carc[[i]])
}

pvals_carc <- as.vector(t(pvals_carc))
pvals_carc[pvals_carc == 0] <- "< 0.001"
table3$P[22:43] <- pvals_carc[!is.na(pvals_carc)]


pvals_carcEB <-  as.data.frame(matrix(rep(NA, 9), ncol = 3))

for (i in 1:length(modelList_carcEB)) {
  pvals_carcEB[i, ] <- p_fun(modelList_carcEB[[i]])
}

pvals_carcEB <- as.vector(t(pvals_carcEB))
pvals_carcEB[pvals_carcEB == 0] <- "< 0.001"
table3$P[44:50] <- pvals_carcEB[!is.na(pvals_carcEB)]

# R squared

rsquared_fun <- function(model) {
  if (is.na(model$coefficients[3]) == TRUE) {
    rsq <- c(round(summary(model)$adj.r.sq, 3), NA)
  } else {
    rsq <- c(round(summary(model)$adj.r.sq, 3), NA, NA)
  }
  return (rsq)
}

rsq_eb <-  list()

for (i in 1:length(modelList_eb)) {
  rsq_eb[[i]] <- rsquared_fun(modelList_eb[[i]])
}

table3$Rsq[1:21] <- unlist(rsq_eb)

rsq_carc <-  list()

for (i in 1:length(modelList_carc)) {
  rsq_carc[[i]] <- rsquared_fun(modelList_carc[[i]])
}

table3$Rsq[22:43] <- unlist(rsq_carc)

rsq_carcEB <-  list()

for (i in 1:length(modelList_carcEB)) {
  rsq_carcEB[[i]] <- rsquared_fun(modelList_carcEB[[i]])
}

table3$Rsq[44:50] <- unlist(rsq_carcEB)

# RMSE & rCV

rmse_table <- function(chemvar, dxavar1, dxavar2, model, loos) {
  rmseList <- NULL
  if(is.na(dxavar2) == TRUE){
    rmseList <- c(rmse(chemvar, summary(model)$coefficients[1,1] + summary(model)$coefficients[2,1] * dxavar1), as.numeric(loos$results[2]))
  } else {
    rmseList <- c(rmse(chemvar, summary(model)$coefficients[1,1] + summary(model)$coefficients[2,1] * dxavar1 + summary(model)$coefficients[3,1] * dxavar2), as.numeric(loos$results[2]), NA)
  }
  return(rmseList)
}

rCV_table <- function(chemvar, dxavar1, dxavar2, model, loos) {
  rCVList <- NULL
  if(is.na(dxavar2) == TRUE){
    rCVList <- c(rCV(chemvar, summary(model)$coefficients[1,1] + summary(model)$coefficients[2,1] * dxavar1), rCV_loo(chemvar, loos))
  } else {
    rCVList <- c(rCV(chemvar, summary(model)$coefficients[1,1] + summary(model)$coefficients[2,1] * dxavar1 + summary(model)$coefficients[3,1] * dxavar2),
                 rCV_loo(chemvar, loos), NA)
  }
  return(rCVList)
}

rmseList_eb <- list()

for (i in 1:9) {
  if (is.na(varList_eb[i,3]) == TRUE) {
    rmseList_eb[[i]] <- round(rmse_table(emptybody[, varList_eb[i, 1]], emptybody[, varList_eb[i, 2]], NA, modelList_eb[[i]], loos_eb[[i]]), 2)
  } else {
      rmseList_eb[[i]] <- round(rmse_table(emptybody[, varList_eb[i, 1]], emptybody[, varList_eb[i, 2]], emptybody[, varList_eb[i, 3]], modelList_eb[[i]], loos_eb[[i]]), 2)
    }
}
 
table3$RMSE[1:21] <- unlist(rmseList_eb) 

rcvList_eb <- list()

for (i in 1:9) {
  if (is.na(varList_eb[i,3]) == TRUE) {
    rcvList_eb[[i]] <- rCV_table(emptybody[, varList_eb[i, 1]], emptybody[, varList_eb[i, 2]], NA, modelList_eb[[i]], loos_eb[[i]])
  } else {
      rcvList_eb[[i]] <- rCV_table(emptybody[, varList_eb[i, 1]], emptybody[, varList_eb[i, 2]], emptybody[, varList_eb[i, 3]], modelList_eb[[i]], loos_eb[[i]])
    }
}

table3$rCV[1:21] <- unlist(rcvList_eb) 

varList_carc <- as.data.frame(matrix(data = c(5,  22, NA, 11, 24, 25, 6, 24, NA, 7, 23, 24, 12, 23, NA, 13, 23, 24, 10, 24, NA, 8, 24, NA, 9, 25, 24), 
                                   ncol = 3, byrow = TRUE))

rmseList_carc <- list()

for (i in 1:9) {
  if (is.na(varList_carc[i,3]) == TRUE) {
    rmseList_carc[[i]] <- round(rmse_table(carcass[, varList_carc[i, 1]], carcass[, varList_carc[i, 2]], NA, modelList_carc[[i]], loos_carc[[i]]), 2)
  } else {
      rmseList_carc[[i]] <- round(rmse_table(carcass[, varList_carc[i, 1]], carcass[, varList_carc[i, 2]], carcass[, varList_carc[i, 3]], modelList_carc[[i]], loos_carc[[i]]), 2)
    }
}
 
table3$RMSE[22:43] <- unlist(rmseList_carc)  

rcvList_carc <- list()

for (i in 1:9) {
  if (is.na(varList_carc[i,3]) == TRUE) {
    rcvList_carc[[i]] <- rCV_table(carcass[, varList_carc[i, 1]], carcass[, varList_carc[i, 2]], NA, modelList_carc[[i]], loos_carc[[i]])
  } else {
      rcvList_carc[[i]] <- rCV_table(carcass[, varList_carc[i, 1]], carcass[, varList_carc[i, 2]], carcass[, varList_carc[i, 3]], modelList_carc[[i]], loos_carc[[i]])
    }
}

table3$rCV[22:43] <- unlist(rcvList_carc) 

varList_carcEB <- as.data.frame(matrix(data = c(8, 21, NA, 11, 22, NA, 10, 23, 22), 
                                   ncol = 3, byrow = TRUE))

rmseList_carcEB <- list()

for (i in 1:3) {
  if (is.na(varList_carcEB[i,3]) == TRUE) {
    rmseList_carcEB[[i]] <- round(rmse_table(eb[, varList_carcEB[i, 1]], eb[, varList_carcEB[i, 2]], NA, modelList_carcEB[[i]], loos_carcEB[[i]]), 2)
  } else {
      rmseList_carcEB[[i]] <- round(rmse_table(eb[, varList_carcEB[i, 1]], eb[, varList_carcEB[i, 2]], eb[, varList_carcEB[i, 3]], modelList_carcEB[[i]], loos_carcEB[[i]]), 2)
    }
}
 
table3$RMSE[44:50] <- unlist(rmseList_carcEB) 

rcvList_carcEB <- list()

for (i in 1:3) {
  if (is.na(varList_carcEB[i,3]) == TRUE) {
    rcvList_carcEB[[i]] <- rCV_table(eb[, varList_carcEB[i, 1]], eb[, varList_carcEB[i, 2]], NA, modelList_carcEB[[i]], loos_carcEB[[i]])
  } else {
      rcvList_carcEB[[i]] <- rCV_table(eb[, varList_carcEB[i, 1]], eb[, varList_carcEB[i, 2]], eb[, varList_carcEB[i, 3]], modelList_carcEB[[i]], loos_carcEB[[i]])
    }
}

table3$rCV[44:50] <- unlist(rcvList_carcEB) 

write.table(table3, file = "table3.txt", sep = "\t")
```


## Table S1: Effects of dietary treatment on slopes and intercepts

```{r}
model_names_eb = c("mBW_eb","mENERGY_eb", "mWATER_eb", "mASH_eb", "mCa_eb", "mP_eb", "mCP_eb", "mN_eb", "mLipid_eb")
modelList_eb = lapply(model_names_eb, get)
names(modelList_eb) = model_names_eb

model_names_eb_treat = c("mBW_eb_treat","mENERGY_eb_treat", "mWATER_eb_treat", "mASH_eb_treat", "mCa_eb_treat", "mP_eb_treat", 
                         "mCP_eb_treat", "mN_eb_treat", "mLipid_eb_treat")
modelList_eb_treat = lapply(model_names_eb_treat, get)
names(modelList_eb_treat) = model_names_eb_treat

model_names_carc = c("mBW_carc","mENERGY_carc", "mWATER_carc", "mASH_carc", "mCa_carc", "mP_carc", "mCP_carc", "mN_carc", "mLipid_carc")
modelList_carc = lapply(model_names_carc, get)
names(modelList_carc) = model_names_carc

model_names_carc_treat = c("mBW_carc_treat","mENERGY_carc_treat", "mWATER_carc_treat", "mASH_carc_treat", "mCa_carc_treat", "mP_carc_treat",
                           "mCP_carc_treat", "mN_carc_treat", "mLipid_carc_treat")
modelList_carc_treat = lapply(model_names_carc_treat, get)
names(modelList_carc_treat) = model_names_carc_treat

tableS1 <- as.data.frame(matrix(data = rep(NA, 104), ncol = 8))
names(tableS1) <- rep(c("deltaIntercept", "deltaSlope", "deltaRsq", "p-value"), 2)
row.names(tableS1) <- c("weight (g)", "energy content (MJ)", "x", "water (g)", "ash (g)", "y", "Ca (g)", "P (g)", "q", "CP (g)", "N (g)", "lipid (g)","z")

# delta intercept

deltaI_eb <- NULL
deltaI_carc <- NULL
for (i in 1:length(model_names_eb)) {
  deltaI_eb[i] <- round(summary(modelList_eb_treat[[i]])$coefficients[1,1] - summary(modelList_eb[[i]])$coefficients[1,1], 3)
  deltaI_carc[i] <- round(summary(modelList_carc_treat[[i]])$coefficients[1,1] - summary(modelList_carc[[i]])$coefficients[1,1], 3)
}

tableS1[c(1, 2, 4, 5, 7, 8, 10, 11, 12), 1] <- deltaI_eb
tableS1[c(1, 2, 4, 5, 7, 8, 10, 11, 12), 5] <- deltaI_carc

# delta slope

deltaS_eb <- list()  

for (i in 1:length(model_names_eb)) {
  coefs <- as.data.frame(summary(modelList_eb[[i]])$coefficients)
  coefs_treat <- as.data.frame(summary(modelList_eb_treat[[i]])$coefficients)
  if (is.na(coefs[3,1]) == TRUE) {
    deltaS_eb[[i]] <- c(round(coefs_treat[2,1] - coefs[2,1], 3), NA)
  } else {
    deltaS_eb[[i]] <- c(round(coefs_treat[2,1] - coefs[2,1], 3), round(coefs_treat[3,1] - coefs[3,1], 3))
  }
}

deltaS_eb <- unlist(deltaS_eb)
tableS1[c(1:5), 2] <- deltaS_eb[!is.na(deltaS_eb)][c(1:5)]
tableS1[6, 2] <- NA
tableS1[c(7:13), 2] <- deltaS_eb[!is.na(deltaS_eb)][c(6:12)]

deltaS_carc <- list()  

for (i in 1:length(model_names_carc)) {
  coefs <- as.data.frame(summary(modelList_carc[[i]])$coefficients)
  coefs_treat <- as.data.frame(summary(modelList_carc_treat[[i]])$coefficients)
  if (is.na(coefs[3,1]) == TRUE) {
    deltaS_carc[[i]] <- c(round(coefs_treat[2,1] - coefs[2,1], 3), NA)
  } else {
    deltaS_carc[[i]] <- c(round(coefs_treat[2,1] - coefs[2,1], 3), round(coefs_treat[3,1] - coefs[3,1], 3))
  }
}

deltaS_carc <- unlist(deltaS_carc)
tableS1[ ,6] <- deltaS_carc[!is.na(deltaS_carc)]

# delta rsq

deltaR_eb <- NULL
deltaR_carc <- NULL

for (i in 1:length(model_names_eb)) {
  deltaR_eb[i] <- round(summary(modelList_eb_treat[[i]])$adj.r.squared - summary(modelList_eb[[i]])$adj.r.squared, 3)
  deltaR_carc[i] <- round(summary(modelList_carc_treat[[i]])$adj.r.squared - summary(modelList_carc[[i]])$adj.r.squared, 3)
}

tableS1[c(1, 2, 4, 5, 7, 8, 10, 11, 12), 3] <- deltaR_eb
tableS1[c(1, 2, 4, 5, 7, 8, 10, 11, 12), 7] <- deltaR_carc

# p-values

p_eb <- NULL
p_carc <- NULL

for (i in 1:length(model_names_eb)) {
  coefs_eb_treat <- as.data.frame(summary(modelList_eb_treat[[i]])$coefficients)
  coefs_carc_treat <- as.data.frame(summary(modelList_carc_treat[[i]])$coefficients)
  p_eb[i] <- round(coefs_eb_treat[nrow(coefs_eb_treat-1), 4], 3)
  p_carc[i] <- round(coefs_carc_treat[nrow(coefs_carc_treat-1), 4], 3)
}

tableS1[c(1, 2, 4, 5, 7, 8, 10, 11, 12), 4] <- p_eb
tableS1[c(1, 2, 4, 5, 7, 8, 10, 11, 12), 8] <- p_carc

write.table(tableS1, file = "tableS1.txt", sep = "\t")
```


## Table S2: Intercepts and slopes for global and regional regressions

Dataframes used for Figures 2 and 3 used, RMSE and rCV added

```{r}
rmse_eb <- list()

for (i in 1:nrow(varList_eb)) {
  rmse_eb[[i]] <- c(rmse(emptybody[, varList_eb[i, 1]], eb_df_full$intercept[i] + eb_df_full$x1[i] * emptybody[, varList_eb [i,2]]),
                    rmse(eb_20[, varList_eb[i, 1]], eb_df_20$intercept[i] + eb_df_20$x1[i] * eb_20[, varList_eb [i,2]]),
                    rmse(eb_60[, varList_eb[i, 1]], eb_df_60$intercept[i] + eb_df_60$x1[i] * eb_60[, varList_eb [i,2]]),
                    rmse(eb_100[, varList_eb[i, 1]], eb_df_100$intercept[i] + eb_df_100$x1[i] * eb_100[, varList_eb [i,2]]))
}

# energy [2], phosphorus [6] and lipid [9] have 2 predictor variables

rmse_eb[[2]] <- 
c(rmse(emptybody[, varList_eb[2, 1]], 
       eb_df_full$intercept[2] + eb_df_full$x1[2] * emptybody[, varList_eb [2,2]] + eb_df_full$x2[2] * emptybody[, varList_eb [2,3]]),
  rmse(eb_20[, varList_eb[2, 1]], 
       eb_df_20$intercept[2] + eb_df_20$x1[2] * eb_20[, varList_eb [2,2]] + eb_df_20$x2[2] * eb_20[, varList_eb [2,3]]),
  rmse(eb_60[, varList_eb[2, 1]], 
       eb_df_60$intercept[2] + eb_df_60$x1[2] * eb_60[, varList_eb [2,2]] + eb_df_60$x2[2] * eb_60[, varList_eb [2,3]]),
  rmse(eb_100[, varList_eb[2, 1]], 
       eb_df_100$intercept[2] + eb_df_100$x1[2] * eb_100[, varList_eb [2,2]] + eb_df_100$x2[2] * eb_100[, varList_eb [2,3]]))

rmse_eb[[6]] <- 
  c(rmse(emptybody[, varList_eb[6, 1]], 
         eb_df_full$intercept[6] + eb_df_full$x1[6] * emptybody[, varList_eb [6,2]] + eb_df_full$x2[6] * emptybody[, varList_eb [6,3]]),
    rmse(eb_20[, varList_eb[6, 1]], 
         eb_df_20$intercept[6] + eb_df_20$x1[6] * eb_20[, varList_eb [6,2]] + eb_df_20$x2[6] * eb_20[, varList_eb [6,3]]),
    rmse(eb_60[, varList_eb[6, 1]], 
         eb_df_60$intercept[6] + eb_df_60$x1[6] * eb_60[, varList_eb [6,2]] + eb_df_60$x2[6] * eb_60[, varList_eb [6,3]]),
    rmse(eb_100[, varList_eb[6, 1]], 
         eb_df_100$intercept[6] + eb_df_100$x1[6] * eb_100[, varList_eb [6,2]] + eb_df_100$x2[6] * eb_100[, varList_eb [6,3]]))

rmse_eb[[9]] <- 
  c(rmse(emptybody[, varList_eb[9, 1]], 
         eb_df_full$intercept[9] + eb_df_full$x1[9] * emptybody[, varList_eb [9,2]] + eb_df_full$x2[9] * emptybody[, varList_eb [9,3]]),
    rmse(eb_20[, varList_eb[9, 1]], 
         eb_df_20$intercept[9] + eb_df_20$x1[9] * eb_20[, varList_eb [9,2]] + eb_df_20$x2[9] * eb_20[, varList_eb [9,3]]),
    rmse(eb_60[, varList_eb[9, 1]], 
         eb_df_60$intercept[9] + eb_df_60$x1[9] * eb_60[, varList_eb [9,2]] + eb_df_60$x2[9] * eb_60[, varList_eb [9,3]]),
    rmse(eb_100[, varList_eb[9, 1]], 
         eb_df_100$intercept[9] + eb_df_100$x1[9] * eb_100[, varList_eb [9,2]] + eb_df_100$x2[9] * eb_100[, varList_eb [9,3]]))

rCV_eb <- list()

for (i in 1:nrow(varList_eb)) {
  rCV_eb[[i]] <- c(rCV(emptybody[, varList_eb[i, 1]], eb_df_full$intercept[i] + eb_df_full$x1[i] * emptybody[, varList_eb [i,2]]),
                    rCV(eb_20[, varList_eb[i, 1]], eb_df_20$intercept[i] + eb_df_20$x1[i] * eb_20[, varList_eb [i,2]]),
                    rCV(eb_60[, varList_eb[i, 1]], eb_df_60$intercept[i] + eb_df_60$x1[i] * eb_60[, varList_eb [i,2]]),
                    rCV(eb_100[, varList_eb[i, 1]], eb_df_100$intercept[i] + eb_df_100$x1[i] * eb_100[, varList_eb [i,2]]))
}

# energy [2], phosphorus [6] and lipid [9] have 2 predictor variables

rCV_eb[[2]] <- 
  c(rCV(emptybody[, varList_eb[2, 1]], 
         eb_df_full$intercept[2] + eb_df_full$x1[2] * emptybody[, varList_eb [2,2]] + eb_df_full$x2[2] * emptybody[, varList_eb [2,3]]),
    rCV(eb_20[, varList_eb[2, 1]], 
         eb_df_20$intercept[2] + eb_df_20$x1[2] * eb_20[, varList_eb [2,2]] + eb_df_20$x2[2] * eb_20[, varList_eb [2,3]]),
    rCV(eb_60[, varList_eb[2, 1]], 
         eb_df_60$intercept[2] + eb_df_60$x1[2] * eb_60[, varList_eb [2,2]] + eb_df_60$x2[2] * eb_60[, varList_eb [2,3]]),
    rCV(eb_100[, varList_eb[2, 1]], 
         eb_df_100$intercept[2] + eb_df_100$x1[2] * eb_100[, varList_eb [2,2]] + eb_df_100$x2[2] * eb_100[, varList_eb [2,3]]))

rCV_eb[[6]] <- 
  c(rCV(emptybody[, varList_eb[6, 1]], 
         eb_df_full$intercept[6] + eb_df_full$x1[6] * emptybody[, varList_eb [6,2]] + eb_df_full$x2[6] * emptybody[, varList_eb [6,3]]),
    rCV(eb_20[, varList_eb[6, 1]], 
         eb_df_20$intercept[6] + eb_df_20$x1[6] * eb_20[, varList_eb [6,2]] + eb_df_20$x2[6] * eb_20[, varList_eb [6,3]]),
    rCV(eb_60[, varList_eb[6, 1]], 
         eb_df_60$intercept[6] + eb_df_60$x1[6] * eb_60[, varList_eb [6,2]] + eb_df_60$x2[6] * eb_60[, varList_eb [6,3]]),
    rCV(eb_100[, varList_eb[6, 1]], 
         eb_df_100$intercept[6] + eb_df_100$x1[6] * eb_100[, varList_eb [6,2]] + eb_df_100$x2[6] * eb_100[, varList_eb [6,3]]))

rCV_eb[[9]] <- 
  c(rCV(emptybody[, varList_eb[9, 1]], 
         eb_df_full$intercept[9] + eb_df_full$x1[9] * emptybody[, varList_eb [9,2]] + eb_df_full$x2[9] * emptybody[, varList_eb [9,3]]),
    rCV(eb_20[, varList_eb[9, 1]], 
         eb_df_20$intercept[9] + eb_df_20$x1[9] * eb_20[, varList_eb [9,2]] + eb_df_20$x2[9] * eb_20[, varList_eb [9,3]]),
    rCV(eb_60[, varList_eb[9, 1]], 
         eb_df_60$intercept[9] + eb_df_60$x1[9] * eb_60[, varList_eb [9,2]] + eb_df_60$x2[9] * eb_60[, varList_eb [9,3]]),
    rCV(eb_100[, varList_eb[9, 1]], 
         eb_df_100$intercept[9] + eb_df_100$x1[9] * eb_100[, varList_eb [9,2]] + eb_df_100$x2[9] * eb_100[, varList_eb [9,3]]))
```

```{r}
eb_df$rmse <- unlist(rmse_eb)
eb_df$rCV <- unlist(rCV_eb)
```

```{r}
rmse_carc <- list()

for (i in 1:nrow(varList_carc)) {
  rmse_carc[[i]] <- c(
    rmse(carcass[, varList_carc[i, 1]], carc_df_full$intercept[i] + carc_df_full$x1[i] * carcass[, varList_carc [i,2]]),
    rmse(carc_20[, varList_carc[i, 1]], carc_df_20$intercept[i] + carc_df_20$x1[i] * carc_20[, varList_carc [i,2]]),
    rmse(carc_60[, varList_carc[i, 1]], carc_df_60$intercept[i] + carc_df_60$x1[i] * carc_60[, varList_carc [i,2]]),
    rmse(carc_100[, varList_carc[i, 1]], carc_df_100$intercept[i] + carc_df_100$x1[i] * carc_100[, varList_carc [i,2]]))
}

# energy [2], ash [4], phosphorus [6] and lipid [9] have 2 predictor variables

rmse_carc[[2]] <- 
  c(rmse(carcass[, varList_carc[2, 1]], 
         carc_df_full$intercept[2] + carc_df_full$x1[2] * carcass[, varList_carc [2,2]] + carc_df_full$x2[2] * carcass[, varList_carc [2,3]]),
    rmse(carc_20[, varList_carc[2, 1]], 
         carc_df_20$intercept[2] + carc_df_20$x1[2] * carc_20[, varList_carc [2,2]] + carc_df_20$x2[2] * carc_20[, varList_carc [2,3]]),
    rmse(carc_60[, varList_carc[2, 1]], 
         carc_df_60$intercept[2] + carc_df_60$x1[2] * carc_60[, varList_carc [2,2]] + carc_df_60$x2[2] * carc_60[, varList_carc [2,3]]),
    rmse(carc_100[, varList_carc[2, 1]], 
         carc_df_100$intercept[2] + carc_df_100$x1[2] * carc_100[, varList_carc [2,2]] + carc_df_100$x2[2] * carc_100[, varList_carc [2,3]]))

rmse_carc[[4]] <- 
  c(rmse(carcass[, varList_carc[4, 1]], 
         carc_df_full$intercept[4] + carc_df_full$x1[4] * carcass[, varList_carc [4,2]] + carc_df_full$x2[4] * carcass[, varList_carc [4,3]]),
    rmse(carc_20[, varList_carc[4, 1]], 
         carc_df_20$intercept[4] + carc_df_20$x1[4] * carc_20[, varList_carc [4,2]] + carc_df_20$x2[4] * carc_20[, varList_carc [4,3]]),
    rmse(carc_60[, varList_carc[4, 1]], 
         carc_df_60$intercept[4] + carc_df_60$x1[4] * carc_60[, varList_carc [4,2]] + carc_df_60$x2[4] * carc_60[, varList_carc [4,3]]),
    rmse(carc_100[, varList_carc[4, 1]], 
         carc_df_100$intercept[4] + carc_df_100$x1[4] * carc_100[, varList_carc [4,2]] + carc_df_100$x2[4] * carc_100[, varList_carc [4,3]]))

rmse_carc[[6]] <- 
  c(rmse(carcass[, varList_carc[6, 1]], 
         carc_df_full$intercept[6] + carc_df_full$x1[6] * carcass[, varList_carc [6,2]] + carc_df_full$x2[6] * carcass[, varList_carc [6,3]]),
    rmse(carc_20[, varList_carc[6, 1]], 
         carc_df_20$intercept[6] + carc_df_20$x1[6] * carc_20[, varList_carc [6,2]] + carc_df_20$x2[6] * carc_20[, varList_carc [6,3]]),
    rmse(carc_60[, varList_carc[6, 1]], 
         carc_df_60$intercept[6] + carc_df_60$x1[6] * carc_60[, varList_carc [6,2]] + carc_df_60$x2[6] * carc_60[, varList_carc [6,3]]),
    rmse(carc_100[, varList_carc[6, 1]], 
         carc_df_100$intercept[6] + carc_df_100$x1[6] * carc_100[, varList_carc [6,2]] + carc_df_100$x2[6] * carc_100[, varList_carc [6,3]]))

rmse_carc[[9]] <- 
  c(rmse(carcass[, varList_carc[9, 1]], 
         carc_df_full$intercept[9] + carc_df_full$x1[9] * carcass[, varList_carc [9,2]] + carc_df_full$x2[9] * carcass[, varList_carc [9,3]]),
    rmse(carc_20[, varList_carc[9, 1]], 
         carc_df_20$intercept[9] + carc_df_20$x1[9] * carc_20[, varList_carc [9,2]] + carc_df_20$x2[9] * carc_20[, varList_carc [9,3]]),
    rmse(carc_60[, varList_carc[9, 1]], 
         carc_df_60$intercept[9] + carc_df_60$x1[9] * carc_60[, varList_carc [9,2]] + carc_df_60$x2[9] * carc_60[, varList_carc [9,3]]),
    rmse(carc_100[, varList_carc[9, 1]], 
         carc_df_100$intercept[9] + carc_df_100$x1[9] * carc_100[, varList_carc [9,2]] + carc_df_100$x2[9] * carc_100[, varList_carc [9,3]]))

rCV_carc <- list()

for (i in 1:nrow(varList_carc)) {
  rCV_carc[[i]] <- c(rCV(carcass[, varList_carc[i, 1]], carc_df_full$intercept[i] + carc_df_full$x1[i] * carcass[, varList_carc [i,2]]),
                   rCV(carc_20[, varList_carc[i, 1]], carc_df_20$intercept[i] + carc_df_20$x1[i] * carc_20[, varList_carc [i,2]]),
                   rCV(carc_60[, varList_carc[i, 1]], carc_df_60$intercept[i] + carc_df_60$x1[i] * carc_60[, varList_carc [i,2]]),
                   rCV(carc_100[, varList_carc[i, 1]], carc_df_100$intercept[i] + carc_df_100$x1[i] * carc_100[, varList_carc [i,2]]))
}

# energy [2], phosphorus [6] and lipid [9] have 2 predictor variables

rCV_carc[[2]] <- 
  c(rCV(carcass[, varList_carc[2, 1]], 
        carc_df_full$intercept[2] + carc_df_full$x1[2] * carcass[, varList_carc [2,2]] + carc_df_full$x2[2] * carcass[, varList_carc [2,3]]),
    rCV(carc_20[, varList_carc[2, 1]], 
        carc_df_20$intercept[2] + carc_df_20$x1[2] * carc_20[, varList_carc [2,2]] + carc_df_20$x2[2] * carc_20[, varList_carc [2,3]]),
    rCV(carc_60[, varList_carc[2, 1]], 
        carc_df_60$intercept[2] + carc_df_60$x1[2] * carc_60[, varList_carc [2,2]] + carc_df_60$x2[2] * carc_60[, varList_carc [2,3]]),
    rCV(carc_100[, varList_carc[2, 1]], 
        carc_df_100$intercept[2] + carc_df_100$x1[2] * carc_100[, varList_carc [2,2]] + carc_df_100$x2[2] * carc_100[, varList_carc [2,3]]))

rCV_carc[[4]] <- 
  c(rCV(carcass[, varList_carc[4, 1]], 
        carc_df_full$intercept[4] + carc_df_full$x1[4] * carcass[, varList_carc [4,2]] + carc_df_full$x2[4] * carcass[, varList_carc [4,3]]),
    rCV(carc_20[, varList_carc[4, 1]], 
        carc_df_20$intercept[4] + carc_df_20$x1[4] * carc_20[, varList_carc [4,2]] + carc_df_20$x2[4] * carc_20[, varList_carc [4,3]]),
    rCV(carc_60[, varList_carc[4, 1]], 
        carc_df_60$intercept[4] + carc_df_60$x1[4] * carc_60[, varList_carc [4,2]] + carc_df_60$x2[4] * carc_60[, varList_carc [4,3]]),
    rCV(carc_100[, varList_carc[4, 1]], 
        carc_df_100$intercept[4] + carc_df_100$x1[4] * carc_100[, varList_carc [4,2]] + carc_df_100$x2[4] * carc_100[, varList_carc [4,3]]))

rCV_carc[[6]] <- 
  c(rCV(carcass[, varList_carc[6, 1]], 
        carc_df_full$intercept[6] + carc_df_full$x1[6] * carcass[, varList_carc [6,2]] + carc_df_full$x2[6] * carcass[, varList_carc [6,3]]),
    rCV(carc_20[, varList_carc[6, 1]], 
        carc_df_20$intercept[6] + carc_df_20$x1[6] * carc_20[, varList_carc [6,2]] + carc_df_20$x2[6] * carc_20[, varList_carc [6,3]]),
    rCV(carc_60[, varList_carc[6, 1]], 
        carc_df_60$intercept[6] + carc_df_60$x1[6] * carc_60[, varList_carc [6,2]] + carc_df_60$x2[6] * carc_60[, varList_carc [6,3]]),
    rCV(carc_100[, varList_carc[6, 1]], 
        carc_df_100$intercept[6] + carc_df_100$x1[6] * carc_100[, varList_carc [6,2]] + carc_df_100$x2[6] * carc_100[, varList_carc [6,3]]))

rCV_carc[[9]] <- 
  c(rCV(carcass[, varList_carc[9, 1]], 
        carc_df_full$intercept[9] + carc_df_full$x1[9] * carcass[, varList_carc [9,2]] + carc_df_full$x2[9] * carcass[, varList_carc [9,3]]),
    rCV(carc_20[, varList_carc[9, 1]], 
        carc_df_20$intercept[9] + carc_df_20$x1[9] * carc_20[, varList_carc [9,2]] + carc_df_20$x2[9] * carc_20[, varList_carc [9,3]]),
    rCV(carc_60[, varList_carc[9, 1]], 
        carc_df_60$intercept[9] + carc_df_60$x1[9] * carc_60[, varList_carc [9,2]] + carc_df_60$x2[9] * carc_60[, varList_carc [9,3]]),
    rCV(carc_100[, varList_carc[9, 1]], 
        carc_df_100$intercept[9] + carc_df_100$x1[9] * carc_100[, varList_carc [9,2]] + carc_df_100$x2[9] * carc_100[, varList_carc [9,3]]))

```

```{r}
carc_df$rmse <- unlist(rmse_carc)
carc_df$rCV <- unlist(rCV_carc)
```

Table S2:

```{r}
tableS2 <- cbind(eb_df, carc_df)

write.table(tableS2, file = "tableS2.txt", sep = "\t")
```


  
  


## Figure S3

Figures showing the data points and overall regression lines for empty body, including regression lines for each slaughter weight category (20, 60 and 100 kg). In the case of more than one predictor variable the simple regression is shown.

###Body weight

```{r figEB_bw}
pEBw <- ggplot(emptybody, aes(x = DXA_totalMass_eb, y = BW_3)) + 
  geom_point(size=4, alpha = 0.3) +
  geom_smooth(method='lm', formula= y~x, color="gray", linetype=2, size=0.5) +
  geom_segment(aes(x = 15000, xend = 23000, y = as.numeric(coefficients(mBW_eb_20)[1]) + as.numeric(coefficients(mBW_eb_20)[2])*15000, 
                   yend = as.numeric(coefficients(mBW_eb_20)[1]) + as.numeric(coefficients(mBW_eb_20)[2])*23000), color="#E69F00") +
  geom_segment(aes(x = 50000, xend = 65000, y = as.numeric(coefficients(mBW_eb_60)[1]) + as.numeric(coefficients(mBW_eb_60)[2])*50000, 
                   yend = as.numeric(coefficients(mBW_eb_60)[1]) + as.numeric(coefficients(mBW_eb_60)[2])*65000), color="#56B4E9") +
  geom_segment(aes(x = 90000, xend = 110000, y = as.numeric(coefficients(mBW_eb_100)[1]) + as.numeric(coefficients(mBW_eb_100)[2])*90000, 
                   yend = as.numeric(coefficients(mBW_eb_100)[1]) + as.numeric(coefficients(mBW_eb_100)[2])*110000), color="#009E73") +
  ylab("Empty body weight, g") + xlab("Total mass by DXA, g") +
  theme_bw(base_size = 15, base_family = "Arial") +
  theme(panel.grid.major = element_blank(), panel.grid.minor = element_blank()) + 
  theme(legend.position="none")
```

###Energy - needs to be adapted - simple regression with DXA body weight

```{r figEB_energy}
pEBenergy <- ggplot(emptybody, aes(x = DXA_totalMass_eb, y = chem_energy_eb)) + 
  geom_point(size=4, alpha = 0.3) +
  geom_smooth(method='lm', formula= y~x, color="gray", linetype=2, size=0.5) +
  geom_segment(aes(x = 15000, xend = 23000, y = as.numeric(coefficients(mENERGY_eb_20_s)[1]) + as.numeric(coefficients(mENERGY_eb_20_s)[2])*15000, 
                   yend = as.numeric(coefficients(mENERGY_eb_20_s)[1]) + as.numeric(coefficients(mENERGY_eb_20_s)[2])*23000), color="#E69F00") +
  geom_segment(aes(x = 50000, xend = 65000, y = as.numeric(coefficients(mENERGY_eb_60_s)[1]) + as.numeric(coefficients(mENERGY_eb_60_s)[2])*50000, 
                   yend = as.numeric(coefficients(mENERGY_eb_60_s)[1]) + as.numeric(coefficients(mENERGY_eb_60_s)[2])*65000), color="#56B4E9") +
  geom_segment(aes(x = 90000, xend = 110000, y = as.numeric(coefficients(mENERGY_eb_100_s)[1]) + as.numeric(coefficients(mENERGY_eb_100_s)[2])*90000, 
                   yend = as.numeric(coefficients(mENERGY_eb_100_s)[1]) + as.numeric(coefficients(mENERGY_eb_100_s)[2])*110000), color="#009E73") +
  ylab("Energy content, MJ") + xlab("Total mass by DXA, g") +
  theme_bw(base_size = 15, base_family = "Arial") +
  theme(panel.grid.major = element_blank(), panel.grid.minor = element_blank()) + 
  theme(legend.position="none")
```

###Water

```{r figEB_water}
pEBwater <- ggplot(emptybody, aes(x = DXA_lean_eb, y = chem_water_eb)) + 
  geom_point(size=4, alpha = 0.3) +
  geom_smooth(method='lm', formula= y~x, color="gray", linetype=2, size=0.5) +
  geom_segment(aes(x = 15000, xend = 20000, y = as.numeric(coefficients(mWATER_eb_20)[1]) + as.numeric(coefficients(mWATER_eb_20)[2])*15000, 
                   yend = as.numeric(coefficients(mWATER_eb_20)[1]) + as.numeric(coefficients(mWATER_eb_20)[2])*20000), color="#E69F00") +
  geom_segment(aes(x = 45000, xend = 57000, y = as.numeric(coefficients(mWATER_eb_60)[1]) + as.numeric(coefficients(mWATER_eb_60)[2])*45000, 
                   yend = as.numeric(coefficients(mWATER_eb_60)[1]) + as.numeric(coefficients(mWATER_eb_60)[2])*57000), color="#56B4E9") +
  geom_segment(aes(x = 75000, xend = 95000, y = as.numeric(coefficients(mWATER_eb_100)[1]) + as.numeric(coefficients(mWATER_eb_100)[2])*75000, 
                   yend = as.numeric(coefficients(mWATER_eb_100)[1]) + as.numeric(coefficients(mWATER_eb_100)[2])*95000), color="#009E73") +
  ylab("Water, g") + xlab("Lean tissue by DXA, g") +
  theme_bw(base_size = 15, base_family = "Arial") +
  theme(panel.grid.major = element_blank(), panel.grid.minor = element_blank()) + 
  theme(legend.position="none")
```

###Ash

```{r figEB_ash}
pEBash <- ggplot(emptybody, aes(x = DXA_BMC_eb, y = chem_ash_eb)) + 
  geom_point(size=4, alpha = 0.3) +
  geom_smooth(method='lm', formula= y~x, color="gray", linetype=2, size=0.5) +
  geom_segment(aes(x = 350, xend = 470, y = as.numeric(coefficients(mASH_eb_20)[1]) + as.numeric(coefficients(mASH_eb_20)[2])*350, 
                   yend = as.numeric(coefficients(mASH_eb_20)[1]) + as.numeric(coefficients(mASH_eb_20)[2])*470), color="#E69F00") +
  geom_segment(aes(x = 1100, xend = 2200, y = as.numeric(coefficients(mASH_eb_60)[1]) + as.numeric(coefficients(mASH_eb_60)[2])*1100, 
                   yend = as.numeric(coefficients(mASH_eb_60)[1]) + as.numeric(coefficients(mASH_eb_60)[2])*2200), color="#56B4E9") +
  geom_segment(aes(x = 1900, xend = 3000, y = as.numeric(coefficients(mASH_eb_100)[1]) + as.numeric(coefficients(mASH_eb_100)[2])*1900, 
                   yend = as.numeric(coefficients(mASH_eb_100)[1]) + as.numeric(coefficients(mASH_eb_100)[2])*3000), color="#009E73") +
  ylab("Ash, g") + xlab("BMC by DXA, g") +
  theme_bw(base_size = 15, base_family = "Arial") +
  theme(panel.grid.major = element_blank(), panel.grid.minor = element_blank()) + 
  theme(legend.position="none")
```

###Ca

```{r figEB_ca}
pEBca <- ggplot(emptybody, aes(x = DXA_BMC_eb, y = chem_Ca_eb)) + 
  geom_point(size=4, alpha = 0.3) +
  geom_smooth(method='lm', formula= y~x, color="gray", linetype=2, size=0.5) +
  geom_segment(aes(x = 350, xend = 470, y = as.numeric(coefficients(mCa_eb_20)[1]) + as.numeric(coefficients(mCa_eb_20)[2])*350, 
                   yend = as.numeric(coefficients(mCa_eb_20)[1]) + as.numeric(coefficients(mCa_eb_20)[2])*470), color="#E69F00") +
  geom_segment(aes(x = 1100, xend = 1700, y = as.numeric(coefficients(mCa_eb_60)[1]) + as.numeric(coefficients(mCa_eb_60)[2])*1100, 
                   yend = as.numeric(coefficients(mCa_eb_60)[1]) + as.numeric(coefficients(mCa_eb_60)[2])*1700), color="#56B4E9") +
  geom_segment(aes(x = 1900, xend = 2800, y = as.numeric(coefficients(mCa_eb_100)[1]) + as.numeric(coefficients(mCa_eb_100)[2])*1900, 
                   yend = as.numeric(coefficients(mCa_eb_100)[1]) + as.numeric(coefficients(mCa_eb_100)[2])*2800), color="#009E73") +
  ylab("Ca, g") + xlab("BMC by DXA, g") +
  theme_bw(base_size = 15, base_family = "Arial") +
  theme(panel.grid.major = element_blank(), panel.grid.minor = element_blank()) + 
  theme(legend.position="none")
```

###P - needs to be adapted - simple regression with DXA BMC

```{r figEB_p}
pEBp <- ggplot(emptybody, aes(x = DXA_BMC_eb, y = chem_P_eb)) + 
  geom_point(size=4, alpha = 0.3) +
  geom_smooth(method='lm', formula= y~x, color="gray", linetype=2, size=0.5) +
  geom_segment(aes(x = 350, xend = 470, y = as.numeric(coefficients(mP_eb_20_s)[1]) + as.numeric(coefficients(mP_eb_20_s)[2])*350, 
                   yend = as.numeric(coefficients(mP_eb_20_s)[1]) + as.numeric(coefficients(mP_eb_20_s)[2])*470), color="#E69F00") +
  geom_segment(aes(x = 1100, xend = 1700, y = as.numeric(coefficients(mP_eb_60_s)[1]) + as.numeric(coefficients(mP_eb_60_s)[2])*1100, 
                   yend = as.numeric(coefficients(mP_eb_60_s)[1]) + as.numeric(coefficients(mP_eb_60_s)[2])*1700), color="#56B4E9") +
  geom_segment(aes(x = 1900, xend = 2800, y = as.numeric(coefficients(mP_eb_100_s)[1]) + as.numeric(coefficients(mP_eb_100_s)[2])*1900, 
                   yend = as.numeric(coefficients(mP_eb_100_s)[1]) + as.numeric(coefficients(mP_eb_100_s)[2])*2800), color="#009E73") +
  ylab("P, g") + xlab("BMC by DXA, g") +
  theme_bw(base_size = 15, base_family = "Arial") +
  theme(panel.grid.major = element_blank(), panel.grid.minor = element_blank()) + 
  theme(legend.position="none")
```

###CP

```{r figEB_cp}
pEBcp <- ggplot(emptybody, aes(x = DXA_lean_eb, y = chem_CP_eb)) + 
  geom_point(size=4, alpha = 0.3) +
  geom_smooth(method='lm', formula= y~x, color="gray", linetype=2, size=0.5) +
  geom_segment(aes(x = 15000, xend = 20000, y = as.numeric(coefficients(mCP_eb_20)[1]) + as.numeric(coefficients(mCP_eb_20)[2])*15000, 
                   yend = as.numeric(coefficients(mCP_eb_20)[1]) + as.numeric(coefficients(mCP_eb_20)[2])*20000), color="#E69F00") +
  geom_segment(aes(x = 45000, xend = 57000, y = as.numeric(coefficients(mCP_eb_60)[1]) + as.numeric(coefficients(mCP_eb_60)[2])*45000, 
                   yend = as.numeric(coefficients(mCP_eb_60)[1]) + as.numeric(coefficients(mCP_eb_60)[2])*57000), color="#56B4E9") +
  geom_segment(aes(x = 75000, xend = 95000, y = as.numeric(coefficients(mCP_eb_100)[1]) + as.numeric(coefficients(mCP_eb_100)[2])*75000, 
                   yend = as.numeric(coefficients(mCP_eb_100)[1]) + as.numeric(coefficients(mCP_eb_100)[2])*95000), color="#009E73") +
  ylab("CP, g") + xlab("Lean tissue by DXA, g") +
  theme_bw(base_size = 15, base_family = "Arial") +
  theme(panel.grid.major = element_blank(), panel.grid.minor = element_blank()) + 
  theme(legend.position="none")
```

###N 

```{r figEB_n}
pEBn <- ggplot(emptybody, aes(x = DXA_lean_eb, y = chem_N_eb)) + 
  geom_point(size=4, alpha = 0.3) +
  geom_smooth(method='lm', formula= y~x, color="gray", linetype=2, size=0.5) +
  geom_segment(aes(x = 15000, xend = 20000, y = as.numeric(coefficients(mN_eb_20)[1]) + as.numeric(coefficients(mN_eb_20)[2])*15000, 
                   yend = as.numeric(coefficients(mN_eb_20)[1]) + as.numeric(coefficients(mN_eb_20)[2])*20000), color="#E69F00") +
  geom_segment(aes(x = 45000, xend = 57000, y = as.numeric(coefficients(mN_eb_60)[1]) + as.numeric(coefficients(mN_eb_60)[2])*45000, 
                   yend = as.numeric(coefficients(mN_eb_60)[1]) + as.numeric(coefficients(mN_eb_60)[2])*57000), color="#56B4E9") +
  geom_segment(aes(x = 75000, xend = 95000, y = as.numeric(coefficients(mN_eb_100)[1]) + as.numeric(coefficients(mN_eb_100)[2])*75000, 
                   yend = as.numeric(coefficients(mN_eb_100)[1]) + as.numeric(coefficients(mN_eb_100)[2])*95000), color="#009E73") +
  ylab("N, g") + xlab("Lean tissue by DXA, g") +
  theme_bw(base_size = 15, base_family = "Arial") +
  theme(panel.grid.major = element_blank(), panel.grid.minor = element_blank()) + 
  theme(legend.position="none")
```

###Lipid

```{r figEB_lip}
pEBlipid <- ggplot(emptybody, aes(x = DXA_fat_eb, y = chem_lipid_eb)) + 
  geom_point(size=4, alpha = 0.3) +
  geom_smooth(method='lm', formula= y~x, color="gray", linetype=2, size=0.5) +
  geom_segment(aes(x = 1000, xend = 2000, y = as.numeric(coefficients(mLipid_eb_20_s)[1]) + as.numeric(coefficients(mLipid_eb_20_s)[2])*1000, 
                   yend = as.numeric(coefficients(mLipid_eb_20_s)[1]) + as.numeric(coefficients(mLipid_eb_20_s)[2])*2000), color="#E69F00") +
  geom_segment(aes(x = 4500, xend = 9500, y = as.numeric(coefficients(mLipid_eb_60_s)[1]) + as.numeric(coefficients(mLipid_eb_60_s)[2])*4500, 
                   yend = as.numeric(coefficients(mLipid_eb_60_s)[1]) + as.numeric(coefficients(mLipid_eb_60_s)[2])*9500), color="#56B4E9") +
  geom_segment(aes(x = 9500, xend = 17000, y = as.numeric(coefficients(mLipid_eb_100_s)[1]) + as.numeric(coefficients(mLipid_eb_100_s)[2])*9500, 
                   yend = as.numeric(coefficients(mLipid_eb_100_s)[1]) + as.numeric(coefficients(mLipid_eb_100_s)[2])*17000), color="#009E73") +
  ylab("Lipid, g") + xlab("Fat tissue by DXA, g") +
  theme_bw(base_size = 15, base_family = "Arial") +
  theme(panel.grid.major = element_blank(), panel.grid.minor = element_blank()) + 
  theme(legend.position="none")
```

###Figure S3

```{r}
multiplot(pEBw, pEBash, pEBcp, pEBenergy, pEBca, pEBn, pEBwater, pEBp, pEBlipid, cols = 3)

png("FigS3.png", width = 1000, height = 1000)
multiplot(pEBw, pEBash, pEBcp, pEBenergy, pEBca, pEBn, pEBwater, pEBp, pEBlipid, cols = 3)
dev.off()
```

## Figure S4

Figures showing the data points and overall regression lines for carcass, including regression lines for each slaughter weight category (20, 60 and 100 kg). In the case of more than one predictor variable the simple regression is shown.

###Body weight

```{r figCarc_bw}
pCARCw <- ggplot(carcass, aes(x = DXA_totalMass_carc, y = coldCW)) + 
  geom_point(size=4, alpha = 0.3) +
  geom_smooth(method='lm', formula= y~x, color="gray", linetype=2, size=0.5) +
  geom_segment(aes(x = 10000, xend = 17000, y = as.numeric(coefficients(mBW_carc_20)[1]) + as.numeric(coefficients(mBW_carc_20)[2])*10000, 
                   yend = as.numeric(coefficients(mBW_carc_20)[1]) + as.numeric(coefficients(mBW_carc_20)[2])*17000), color="#E69F00") +
  geom_segment(aes(x = 39000, xend = 51000, y = as.numeric(coefficients(mBW_carc_60)[1]) + as.numeric(coefficients(mBW_carc_60)[2])*39000, 
                   yend = as.numeric(coefficients(mBW_carc_60)[1]) + as.numeric(coefficients(mBW_carc_60)[2])*51000), color="#56B4E9") +
  geom_segment(aes(x = 69000, xend = 83000, y = as.numeric(coefficients(mBW_carc_100)[1]) + as.numeric(coefficients(mBW_carc_100)[2])*69000, 
                   yend = as.numeric(coefficients(mBW_carc_100)[1]) + as.numeric(coefficients(mBW_carc_100)[2])*83000), color="#009E73") +
  ylab("Empty body weight, g") + xlab("Total mass by DXA, g") +
  theme_bw(base_size = 15, base_family = "Arial") +
  theme(panel.grid.major = element_blank(), panel.grid.minor = element_blank()) + 
  theme(legend.position="none")
```

###Energy - needs to be adapted - simple regression with DXA body weight

```{r figCarc_energy}
pCARCenergy <- ggplot(carcass, aes(x = DXA_totalMass_carc, y = chem_energy_carc)) + 
  geom_point(size=4, alpha = 0.3) +
  geom_smooth(method='lm', formula= y~x, color="gray", linetype=2, size=0.5) +
  geom_segment(aes(x = 10000, xend = 17000, y = as.numeric(coefficients(mENERGY_carc_20_s)[1]) + 
                     as.numeric(coefficients(mENERGY_carc_20_s)[2])*10000, yend = as.numeric(coefficients(mENERGY_carc_20_s)[1]) +
                     as.numeric(coefficients(mENERGY_carc_20_s)[2])*17000), color="#E69F00") +
  geom_segment(aes(x = 38000, xend = 52000, y = as.numeric(coefficients(mENERGY_carc_60_s)[1]) +
                     as.numeric(coefficients(mENERGY_carc_60_s)[2])*38000, yend = as.numeric(coefficients(mENERGY_carc_60_s)[1]) +
                     as.numeric(coefficients(mENERGY_carc_60_s)[2])*52000), color="#56B4E9") +
  geom_segment(aes(x = 67000, xend = 84000, y = as.numeric(coefficients(mENERGY_carc_100_s)[1]) +
                     as.numeric(coefficients(mENERGY_carc_100_s)[2])*67000, yend = as.numeric(coefficients(mENERGY_carc_100_s)[1]) +
                     as.numeric(coefficients(mENERGY_carc_100_s)[2])*84000), color="#009E73") +
  ylab("Energy content, MJ") + xlab("Total mass by DXA, g") +
  theme_bw(base_size = 15, base_family = "Arial") +
  theme(panel.grid.major = element_blank(), panel.grid.minor = element_blank()) + 
  theme(legend.position="none")
```

###Water

```{r figCarc_water}
pCARCwater <- ggplot(carcass, aes(x = DXA_lean_carc, y = chem_water_carc)) + 
  geom_point(size=4, alpha = 0.3) +
  geom_smooth(method='lm', formula= y~x, color="gray", linetype=2, size=0.5) +
  geom_segment(aes(x = 9000, xend = 15000, y = as.numeric(coefficients(mWATER_carc_20)[1]) + as.numeric(coefficients(mWATER_carc_20)[2])*9000, 
                   yend = as.numeric(coefficients(mWATER_carc_20)[1]) + as.numeric(coefficients(mWATER_carc_20)[2])*15000), color="#E69F00") +
  geom_segment(aes(x = 30000, xend = 42000, y = as.numeric(coefficients(mWATER_carc_60)[1]) + as.numeric(coefficients(mWATER_carc_60)[2])*30000, 
                   yend = as.numeric(coefficients(mWATER_carc_60)[1]) + as.numeric(coefficients(mWATER_carc_60)[2])*42000), color="#56B4E9") +
  geom_segment(aes(x = 52000, xend = 70000, y = as.numeric(coefficients(mWATER_carc_100)[1]) + as.numeric(coefficients(mWATER_carc_100)[2])*52000,
                   yend = as.numeric(coefficients(mWATER_carc_100)[1]) + as.numeric(coefficients(mWATER_carc_100)[2])*70000), color="#009E73") +
  ylab("Water, g") + xlab("Lean tissue by DXA, g") +
  theme_bw(base_size = 15, base_family = "Arial") +
  theme(panel.grid.major = element_blank(), panel.grid.minor = element_blank()) + 
  theme(legend.position="none")
```

###Ash needs to be adapted - simple regression with DXA BMC

```{r figCarc_ash}
pCARCash <- ggplot(carcass, aes(x = DXA_BMC_carc, y = chem_ash_carc)) + 
  geom_point(size=4, alpha = 0.3) +
  geom_smooth(method='lm', formula= y~x, color="gray", linetype=2, size=0.5) +
  geom_segment(aes(x = 350, xend = 550, y = as.numeric(coefficients(mASH_carc_20_s)[1]) + as.numeric(coefficients(mASH_carc_20_s)[2])*350, 
                   yend = as.numeric(coefficients(mASH_carc_20_s)[1]) + as.numeric(coefficients(mASH_carc_20_s)[2])*550), color="#E69F00") +
  geom_segment(aes(x = 1200, xend = 2100, y = as.numeric(coefficients(mASH_carc_60_s)[1]) + as.numeric(coefficients(mASH_carc_60_s)[2])*1200, 
                   yend = as.numeric(coefficients(mASH_carc_60_s)[1]) + as.numeric(coefficients(mASH_carc_60_s)[2])*2100), color="#56B4E9") +
  geom_segment(aes(x = 1900, xend = 3200, y = as.numeric(coefficients(mASH_carc_100_s)[1]) + as.numeric(coefficients(mASH_carc_100_s)[2])*1900, 
                   yend = as.numeric(coefficients(mASH_carc_100_s)[1]) + as.numeric(coefficients(mASH_carc_100_s)[2])*3200), color="#009E73") +
  ylab("Ash, g") + xlab("BMC by DXA, g") +
  theme_bw(base_size = 15, base_family = "Arial") +
  theme(panel.grid.major = element_blank(), panel.grid.minor = element_blank()) + 
  theme(legend.position="none")
```

###Ca

```{r figCarc_ca}
pCARCca <- ggplot(carcass, aes(x = DXA_BMC_carc, y = chem_Ca_carc)) + 
  geom_point(size=4, alpha = 0.3) +
  geom_smooth(method='lm', formula= y~x, color="gray", linetype=2, size=0.5) +
  geom_segment(aes(x = 350, xend = 500, y = as.numeric(coefficients(mCa_carc_20)[1]) + as.numeric(coefficients(mCa_carc_20)[2])*350, 
                   yend = as.numeric(coefficients(mCa_carc_20)[1]) + as.numeric(coefficients(mCa_carc_20)[2])*500), color="#E69F00") +
  geom_segment(aes(x = 1100, xend = 2000, y = as.numeric(coefficients(mCa_carc_60)[1]) + as.numeric(coefficients(mCa_carc_60)[2])*1100, 
                   yend = as.numeric(coefficients(mCa_carc_60)[1]) + as.numeric(coefficients(mCa_carc_60)[2])*2000), color="#56B4E9") +
  geom_segment(aes(x = 2000, xend = 3100, y = as.numeric(coefficients(mCa_carc_100)[1]) + as.numeric(coefficients(mCa_carc_100)[2])*2000, 
                   yend = as.numeric(coefficients(mCa_carc_100)[1]) + as.numeric(coefficients(mCa_carc_100)[2])*3100), color="#009E73") +
  ylab("Ca, g") + xlab("BMC by DXA, g") +
  theme_bw(base_size = 15, base_family = "Arial") +
  theme(panel.grid.major = element_blank(), panel.grid.minor = element_blank()) + 
  theme(legend.position="none")
```

###P - needs to be adapted - simple regression with DXA BMC

```{r figCarc_p}
pCARCp <- ggplot(carcass, aes(x = DXA_BMC_carc, y = chem_P_carc)) + 
  geom_point(size=4, alpha = 0.3) +
  geom_smooth(method='lm', formula= y~x, color="gray", linetype=2, size=0.5) +
  geom_segment(aes(x = 350, xend = 500, y = as.numeric(coefficients(mP_carc_20_s)[1]) + as.numeric(coefficients(mP_carc_20_s)[2])*350, 
                   yend = as.numeric(coefficients(mP_carc_20_s)[1]) + as.numeric(coefficients(mP_carc_20_s)[2])*500), color="#E69F00") +
  geom_segment(aes(x = 1200, xend = 1900, y = as.numeric(coefficients(mP_carc_60_s)[1]) + as.numeric(coefficients(mP_carc_60_s)[2])*1200, 
                   yend = as.numeric(coefficients(mP_carc_60_s)[1]) + as.numeric(coefficients(mP_carc_60_s)[2])*1900), color="#56B4E9") +
  geom_segment(aes(x = 2000, xend = 3200, y = as.numeric(coefficients(mP_carc_100_s)[1]) + as.numeric(coefficients(mP_carc_100_s)[2])*2000, 
                   yend = as.numeric(coefficients(mP_carc_100_s)[1]) + as.numeric(coefficients(mP_carc_100_s)[2])*3200), color="#009E73") +
  ylab("P, g") + xlab("BMC by DXA, g") +
  theme_bw(base_size = 15, base_family = "Arial") +
  theme(panel.grid.major = element_blank(), panel.grid.minor = element_blank()) + 
  theme(legend.position="none")
```

###CP

```{r figCarc_CP}
pCARCcp <- ggplot(carcass, aes(x = DXA_lean_carc, y = chem_CP_carc)) + 
  geom_point(size=4, alpha = 0.3) +
  geom_smooth(method='lm', formula= y~x, color="gray", linetype=2, size=0.5) +
  geom_segment(aes(x = 10000, xend = 14000, y = as.numeric(coefficients(mCP_carc_20)[1]) + as.numeric(coefficients(mCP_carc_20)[2])*10000, 
                   yend = as.numeric(coefficients(mCP_carc_20)[1]) + as.numeric(coefficients(mCP_carc_20)[2])*14000), color="#E69F00") +
  geom_segment(aes(x = 30000, xend = 41000, y = as.numeric(coefficients(mCP_carc_60)[1]) + as.numeric(coefficients(mCP_carc_60)[2])*30000, 
                   yend = as.numeric(coefficients(mCP_carc_60)[1]) + as.numeric(coefficients(mCP_carc_60)[2])*41000), color="#56B4E9") +
  geom_segment(aes(x = 52000, xend = 70000, y = as.numeric(coefficients(mCP_carc_100)[1]) + as.numeric(coefficients(mCP_carc_100)[2])*52000, 
                   yend = as.numeric(coefficients(mCP_carc_100)[1]) + as.numeric(coefficients(mCP_carc_100)[2])*70000), color="#009E73") +
  ylab("CP, g") + xlab("Lean tissue by DXA, g") +
  theme_bw(base_size = 15, base_family = "Arial") +
  theme(panel.grid.major = element_blank(), panel.grid.minor = element_blank()) + 
  theme(legend.position="none")
```

###N 

```{r figCarc_n}
pCARCn <- ggplot(carcass, aes(x = DXA_lean_carc, y = chem_N_carc)) + 
  geom_point(size=4, alpha = 0.3) +
  geom_smooth(method='lm', formula= y~x, color="gray", linetype=2, size=0.5) +
  geom_segment(aes(x = 10000, xend = 14000, y = as.numeric(coefficients(mN_carc_20)[1]) + as.numeric(coefficients(mN_carc_20)[2])*10000, 
                   yend = as.numeric(coefficients(mN_carc_20)[1]) + as.numeric(coefficients(mN_carc_20)[2])*14000), color="#E69F00") +
  geom_segment(aes(x = 30000, xend = 41000, y = as.numeric(coefficients(mN_carc_60)[1]) + as.numeric(coefficients(mN_carc_60)[2])*30000, 
                   yend = as.numeric(coefficients(mN_carc_60)[1]) + as.numeric(coefficients(mN_carc_60)[2])*41000), color="#56B4E9") +
  geom_segment(aes(x = 52000, xend = 70000, y = as.numeric(coefficients(mN_carc_100)[1]) + as.numeric(coefficients(mN_carc_100)[2])*52000, 
                   yend = as.numeric(coefficients(mN_carc_100)[1]) + as.numeric(coefficients(mN_carc_100)[2])*70000), color="#009E73") +
  ylab("N, g") + xlab("Lean tissue by DXA, g") +
  theme_bw(base_size = 15, base_family = "Arial") +
  theme(panel.grid.major = element_blank(), panel.grid.minor = element_blank()) + 
  theme(legend.position="none")
```

###Lipid needs to be adapted - simple regression with DXA fat

```{r figCarc_lipid}
pCARClipid <- ggplot(carcass, aes(x = DXA_fat_carc, y = chem_lipid_carc)) + 
  geom_point(size=4, alpha = 0.3) +
  geom_smooth(method='lm', formula= y~x, color="gray", linetype=2, size=0.5) +
  geom_segment(aes(x = 700, xend = 2200, y = as.numeric(coefficients(mLipid_carc_20_s)[1]) + as.numeric(coefficients(mLipid_carc_20_s)[2])*700, 
                   yend = as.numeric(coefficients(mLipid_carc_20_s)[1]) + as.numeric(coefficients(mLipid_carc_20_s)[2])*2200), color="#E69F00") +
  geom_segment(aes(x = 4600, xend = 10200, y = as.numeric(coefficients(mLipid_carc_60_s)[1]) + as.numeric(coefficients(mLipid_carc_60_s)[2])*4600,
                   yend = as.numeric(coefficients(mLipid_carc_60_s)[1]) + as.numeric(coefficients(mLipid_carc_60_s)[2])*10200), color="#56B4E9") +
  geom_segment(aes(x = 9500, xend = 18000, y = as.numeric(coefficients(mLipid_carc_100_s)[1]) +
                     as.numeric(coefficients(mLipid_carc_100_s)[2])*9500, yend = as.numeric(coefficients(mLipid_carc_100_s)[1]) +
                     as.numeric(coefficients(mLipid_carc_100_s)[2])*18000), color="#009E73") +
  ylab("Lipid, g") + xlab("Fat tissue by DXA, g") +
  theme_bw(base_size = 15, base_family = "Arial") +
  theme(panel.grid.major = element_blank(), panel.grid.minor = element_blank()) + 
  theme(legend.position="none")
```

### Figure S4

```{r}
multiplot(pCARCw, pCARCash, pCARCcp, pCARCenergy, pCARCca, pCARCn, pCARCwater, pCARCp, pCARClipid, cols = 3)

png("FigS4.png", width = 1000, height = 1000)
multiplot(pCARCw, pCARCash, pCARCcp, pCARCenergy, pCARCca, pCARCn, pCARCwater, pCARCp, pCARClipid, cols = 3)
dev.off()
```

# COMPARISON WITH PUBLISHED EQUATIONS

## EMPTY BODY

### a) ...Mitchell, Scholz & Conway 1998 _Appl. Radiat. Isot._ 

+ empty body (live pigs 5 - 97 kg)
+ sex not specified
+ chem. analys.
+ gramm protein/water - gramm DXA lean
+ Lunar DPXL instrument -  pencil-beam

#### 1. CP

```{r}
Mit98_pred_CP_eb.a <- -1.062 + 0.22 * emptybody$DXA_lean_eb
```

__published equation:__

$$CP (g) = -1.062 + 0.22 \times Lean$$

$RMSE=$ `r round(rmse(emptybody$chem_CP_eb, Mit98_pred_CP_eb.a))` g

$rCV =$ `r round((rmse(emptybody$chem_CP_eb, Mit98_pred_CP_eb.a)*100)/mean(emptybody$chem_CP_eb), 1)`% of the mean CP content of the live pig by chemical analyses

__own equation:__

$\hat{CP_{EB}= $ `r round(summary(mCP_eb)$coefficients[1,1], 3)` + `r round(summary(mCP_eb)$coefficients[2,1], 3)` $ \times Lean $ 

$RMSE=$ `r round(rmse(emptybody$chem_CP_eb, summary(mCP_eb)$coefficients[1,1] + summary(mCP_eb)$coefficients[2,1] * emptybody$DXA_lean_eb))` g

$rCV =$ `r round((rmse(emptybody$chem_CP_eb, summary(mCP_eb)$coefficients[1,1] + summary(mCP_eb)$coefficients[2,1] * emptybody$DXA_lean_eb)*100)/mean(emptybody$chem_CP_eb), 1)`% of the mean CP content of the live pig by chemical analyses 

#### 2. Water

```{r}
Mit98_pred_Water_eb.a <- 508 + 0.74 * emptybody$DXA_lean_eb
```

__published equation:__

$$Water (g) = 508 + 0.74 \times Lean$$

$RMSE=$ `r round(rmse(emptybody$chem_water_eb, Mit98_pred_Water_eb.a))` g

$rCV =$ `r round((rmse(emptybody$chem_water_eb, Mit98_pred_Water_eb.a)*100)/mean(emptybody$chem_water_eb), 1)`% of the mean water content of the live pig by chemical analyses 

__own equation:__

$\hat{Water_{EB}= $ `r round(summary(mWATER_eb)$coefficients[1,1], 3)` + `r round(summary(mWATER_eb)$coefficients[2,1], 3)` $ \times Lean $ 

$RMSE=$ `r round(rmse(emptybody$chem_water_eb, summary(mWATER_eb)$coefficients[1,1] + summary(mWATER_eb)$coefficients[2,1] * emptybody$DXA_lean_eb))` g

$rCV =$ `r round((rmse(emptybody$chem_water_eb, summary(mWATER_eb)$coefficients[1,1] + summary(mWATER_eb)$coefficients[2,1] * emptybody$DXA_lean_eb)*100)/mean(emptybody$chem_water_eb), 1)`% of the mean water content of the live pig by chemical analyses 

### e) ...Pomar & Rivest 1996 _Can. J. Anim. Sci._ or rather Pomar et al 2007 _R. Bras. Zootec._ 

+ empty body (live pigs avg weight 84.4 (F) - 85 (M) kg)
+ barrows and gilts
+ chem. analys.
+ gramm protein/fat - gramm DXA lean/fat
+ GE Lunar Prodigy -  fan-beam

#### 1. CP

```{r}
Pom98_pred_RP_eb.e <- -1384 + 0.216 * emptybody$DXA_lean_eb
```

__published equation:__

$$CP (g) = −1384 + 0.216 \times Lean$$

$RMSE=$ `r round(rmse(emptybody$chem_CP_eb, Pom98_pred_RP_eb.e))` g

$rCV =$ `r round((rmse(emptybody$chem_CP_eb, Pom98_pred_RP_eb.e)*100)/mean(emptybody$chem_CP_eb), 1)`% of the mean CP content of the live pig by chemical analyses  

__own equation:__

$\hat{CP_{EB}= $ `r round(summary(mCP_eb)$coefficients[1,1], 3)` + `r round(summary(mCP_eb)$coefficients[2,1], 3)` $ \times Lean $ 

$RMSE=$ `r round(rmse(emptybody$chem_CP_eb, summary(mCP_eb)$coefficients[1,1] + summary(mCP_eb)$coefficients[2,1] * emptybody$DXA_lean_eb))` g

$rCV =$ `r round((rmse(emptybody$chem_CP_eb, summary(mCP_eb)$coefficients[1,1] + summary(mCP_eb)$coefficients[2,1] * emptybody$DXA_lean_eb)*100)/mean(emptybody$chem_CP_eb), 1)`% of the mean CP content of the live pig by chemical analyses 

#### 2. Lipid

```{r}
Pom98_pred_Fat_eb.e <- 2825 + 1.009 * emptybody$DXA_fat_eb
mLipid_ebx <- lm(chem_lipid_eb ~ DXA_fat_eb, data = emptybody)
```

__published equation:__

$$Lipid (g) = 2825 + 1.009 \times Fat$$

$RMSE=$ `r round(rmse(emptybody$chem_lipid_eb, Pom98_pred_Fat_eb.e))` g 

$rCV =$ `r round((rmse(emptybody$chem_lipid_eb, Pom98_pred_Fat_eb.e)*100)/mean(emptybody$chem_lipid_eb), 1)`% of the mean lipid content of the live pig by chemical analyses 

__own equation:__

$\hat{Lipid_{EB}} =$ `r round(summary(mLipid_ebx)$coefficients[1,1], 3)` + `r round(summary(mLipid_ebx)$coefficients[2,1], 3)` $ \times Fat $ 

$RMSE=$ `r round(rmse(emptybody$chem_lipid_eb, summary(mLipid_eb)$coefficients[1,1] + summary(mLipid_eb)$coefficients[2,1] * emptybody$DXA_fat_eb + summary(mLipid_eb)$coefficients[3,1] * emptybody$DXA_lean_eb))` g

$rCV =$ `r round((rmse(emptybody$chem_lipid_eb, summary(mLipid_eb)$coefficients[1,1] + summary(mLipid_eb)$coefficients[2,1] * emptybody$DXA_fat_eb + summary(mLipid_eb)$coefficients[3,1] * emptybody$DXA_lean_eb)*100)/mean(emptybody$chem_lipid_eb), 1)`% of the mean lipid content of the live pig by chemical analyses

### d) ...Kipper, Marcoux, Andretta & Pomar 2019 _EAAP No 138_ 

+ empty body (live pigs different weight)
+ barrows
+ chem. analys.
+ kg chem - kg DXA 
+ GE Lunar Prodigy Advance - fan beam

```{r}
Kip19_pred_Fat_eb.f <- 3.371 + 1.116 * (emptybody$DXA_fat_eb/1000)
Kip19_pred_Ash_eb.f <- 0.542 + 1.190 * (emptybody$DXA_BMC_eb/1000)
Kip19_pred_CP_eb.f <- -0.894 + 0.219 * (emptybody$DXA_lean_eb/1000)
Kip19_pred_Water_eb.f <- 2.344 + 0.674 * (emptybody$DXA_lean_eb/1000)
Kip19_pred_ProtWat_eb.f <- 1.450 + 0.893 * (emptybody$DXA_lean_eb/1000)
protWatChem_eb.f <- (emptybody$chem_CP_eb/1000) + (emptybody$chem_water_eb/1000)
emptybody$protWaterChem_kg <- protWatChem_eb.f

emptybody$chem_lipidKG_eb <- emptybody$chem_lipid_eb/1000
emptybody$DXA_fatKG_eb <- emptybody$DXA_fat_eb/1000
emptybody$chem_CPkg_eb <- emptybody$chem_CP_eb/1000
emptybody$DXA_leanKG_eb <- emptybody$DXA_lean_eb/1000
emptybody$chem_ashKG_eb <- emptybody$chem_ash_eb/1000
emptybody$DXA_BMCkg_eb <- emptybody$DXA_BMC_eb/1000
emptybody$chem_waterKG_eb <- emptybody$chem_water_eb/1000
```

#### 1. Lipid

```{r}
mLipidkg_eb <- lm(chem_lipidKG_eb ~ DXA_fatKG_eb + DXA_leanKG_eb, data = emptybody)
mLipidkg_eb.1 <- lm(chem_lipidKG_eb ~ DXA_fatKG_eb, data = emptybody)
```

__published equation:__

$$ Lipid = 3.371 + 1.116 \times DXAfat $$

$RMSE=$ `r round(rmse(emptybody$chem_lipidKG_eb, Kip19_pred_Fat_eb.f), 2)` kg

$rCV =$ `r round((rmse(emptybody$chem_lipidKG_eb, Kip19_pred_Fat_eb.f)*100)/mean(emptybody$chem_lipidKG_eb), 1)`% of the mean fat content of the live pig by chemical analyses 

__own equation:__

$ Lipid \; (kg) = $ `r summary(mLipidkg_eb.1)$coefficients[1,1]` + `r summary(mLipidkg_eb.1)$coefficients[1,2]` $ \times DXAfat $  

$ Lipid \; (kg) = $ `r summary(mLipidkg_eb)$coefficients[1,1]` + `r summary(mLipidkg_eb)$coefficients[1,2]` $ \times DXAfat $  + `r summary(mLipidkg_eb)$coefficients[1,3]` $ \times DXAlean $

$RMSE=$ `r round(rmse(emptybody$chem_lipidKG_eb, summary(mLipidkg_eb)$coefficients[1,1] + summary(mLipidkg_eb)$coefficients[2,1] * emptybody$DXA_fatKG_eb + summary(mLipidkg_eb)$coefficients[3,1] * emptybody$DXA_leanKG_eb), 2)` kg

$rCV =$ `r round((rmse(emptybody$chem_lipidKG_eb, summary(mLipidkg_eb)$coefficients[1,1] + summary(mLipidkg_eb)$coefficients[2,1] * emptybody$DXA_fatKG_eb + summary(mLipidkg_eb)$coefficients[3,1] * emptybody$DXA_leanKG_eb)*100)/mean(emptybody$chem_lipidKG_eb), 1)`% of the mean fat content of the live pig by chemical analyses

$R^2 = $ `r round(summary(mLipidkg_eb)$adj, 2)`

#### 2. Ash

```{r}
mAshkg_eb <- lm(chem_ashKG_eb ~ DXA_BMCkg_eb, data = emptybody)
```

__published equation:__

$$ Ash = 0.542 + 1.190 \times DXABMC $$

$RMSE=$ `r round(rmse(emptybody$chem_ashKG_eb, Kip19_pred_Ash_eb.f), 2)` kg

$rCV =$ `r round((rmse(emptybody$chem_ashKG_eb, Kip19_pred_Ash_eb.f)*100)/mean(emptybody$chem_ashKG_eb), 1)`% of the mean ash content of the live pig by chemical analyses

__own equation:__

$ Ash = $ `r summary(mAshkg_eb)$coefficients[1,1]` + `r summary(mAshkg_eb)$coefficients[1,2]` $ \times DXABMC $  

$RMSE=$ `r round(rmse(emptybody$chem_ashKG_eb, summary(mAshkg_eb)$coefficients[1,1] + summary(mAshkg_eb)$coefficients[2,1] * emptybody$DXA_BMCkg_eb), 2)` kg

$rCV =$ `r round((rmse(emptybody$chem_ashKG_eb, summary(mAshkg_eb)$coefficients[1,1] + summary(mAshkg_eb)$coefficients[2,1] * emptybody$DXA_BMCkg_eb)*100)/mean(emptybody$chem_ashKG_eb), 1)`% of the mean ash content of the live pig by chemical analyses

$R^2 = $ `r round(summary(mAshkg_eb)$adj, 2)`

#### 3. CP

```{r}
mCPkg_eb <- lm(chem_CPkg_eb ~ DXA_leanKG_eb, data = emptybody)
```

__published equation:__

$$ CP \; (kg) = -0.894 + 0.219 \times DXAlean $$

$RMSE=$ `r round(rmse(emptybody$chem_CPkg_eb, Kip19_pred_CP_eb.f), 2)` kg

$rCV =$ `r round((rmse(emptybody$chem_CPkg_eb, Kip19_pred_CP_eb.f)*100)/mean(emptybody$chem_CPkg_eb), 1)`% of the mean CP content of the live pig by chemical analyses

__own equation:__

$ CP \; (kg) = $ `r summary(mCPkg_eb)$coefficients[1,1]` + `r summary(mCPkg_eb)$coefficients[1,2]` $ \times DXAlean $  

$RMSE=$ `r round(rmse(emptybody$chem_CPkg_eb, summary(mCPkg_eb)$coefficients[1,1] + summary(mCPkg_eb)$coefficients[2,1] * emptybody$DXA_leanKG_eb), 2)` kg

$rCV =$ `r round((rmse(emptybody$chem_CPkg_eb, summary(mCPkg_eb)$coefficients[1,1] + summary(mCPkg_eb)$coefficients[2,1] * emptybody$DXA_leanKG_eb)*100)/mean(emptybody$chem_CPkg_eb), 1)`% of the mean CP content of the live pig by chemical analyses 

$R^2 = $ `r round(summary(mCPkg_eb)$adj, 2)`

#### 4. Water

```{r}
mWaterkg_eb <- lm(chem_waterKG_eb ~ DXA_leanKG_eb, data = emptybody)
```

__published equation:__

$$ Water \; (kg)= 2.344 + 0.674 \times DXAlean $$

$RMSE=$ `r round(rmse(emptybody$chem_waterKG_eb, Kip19_pred_Water_eb.f), 2)` kg

$rCV =$ `r round((rmse(emptybody$chem_waterKG_eb, Kip19_pred_Water_eb.f)*100)/mean(emptybody$chem_waterKG_eb), 1)`% of the mean water content of the live pig by chemical analyses 

__own equation:__

$ Water \; (kg)= $ `r summary(mWaterkg_eb)$coefficients[1,1]` + `r summary(mWaterkg_eb)$coefficients[1,2]` $ \times DXAlean $ 

$RMSE=$ `r round(rmse(emptybody$chem_waterKG_eb, summary(mWaterkg_eb)$coefficients[1,1] + summary(mWaterkg_eb)$coefficients[2,1] * emptybody$DXA_leanKG_eb), 2)` kg

$rCV =$ `r round((rmse(emptybody$chem_waterKG_eb, summary(mWaterkg_eb)$coefficients[1,1] + summary(mWaterkg_eb)$coefficients[2,1] * emptybody$DXA_leanKG_eb)*100)/mean(emptybody$chem_waterKG_eb), 1)`% of the mean water content of the live pig by chemical analyses

$R^2 = $ `r round(summary(mWaterkg_eb)$adj, 3)`

#### 5. CP + water

```{r}
mProtWaterkg <- lm(protWaterChem_kg ~ DXA_leanKG_eb, data = emptybody)
```

__published equation:__

$$ CP + Water \; (kg) = 1.450 + 0.893 \times DXAlean $$
$RMSE=$ `r round(rmse(protWatChem_eb.f, Kip19_pred_ProtWat_eb.f), 2)` kg

$rCV =$  `r round((rmse(protWatChem_eb.f, Kip19_pred_ProtWat_eb.f)*100)/mean(protWatChem_eb.f), 1)`% of the mean protein + water content of the live pig by chemical analyses

__own equation:__

$ CP + Water \; (kg) = $ `r summary(mProtWaterkg)$coefficients[1,1]` + `r summary(mProtWaterkg)$coefficients[1,2]` $ \times DXALean $ 

$RMSE=$ `r round(rmse(protWatChem_eb.f, summary(mProtWaterkg)$coefficients[1,1] + summary(mProtWaterkg)$coefficients[2,1] * emptybody$protWaterChem_kg), 2)` kg

$rCV =$ `r round((rmse(protWatChem_eb.f, summary(mProtWaterkg)$coefficients[1,1] + summary(mProtWaterkg)$coefficients[2,1] * emptybody$protWaterChem_kg)*100)/mean(protWatChem_eb.f), 1)`% of the mean protein + water content of the live pig by chemical analyses

$R^2 = $ `r round(summary(mProtWaterkg)$adj, 2)`

## Figure 5

```{r warning=FALSE, message=FALSE}
equation.a1=function(x){-1.062+0.22*x} # published
equation.a2=function(x){coef(mCP_eb)[1]+coef(mCP_eb)[2]*x} # own

pa1 <- ggplot(emptybody, aes(x=DXA_lean_eb, y=chem_CP_eb)) +
  geom_point(shape=19, size=3, col="black") + 
  stat_function(fun=equation.a1,geom="line", color="darkred", linetype=2) + # published
  geom_smooth(method='lm', formula= y~x, color="black", linetype=1, size=.5) + # own
  stat_function(fun=equation.a2,geom="line", color="black", linetype=1) + # own (to get CI and make regression line longer)
  ylim(0, 25000) + xlim(0, 110000) +
  ylab("CP, g") + xlab("Lean tissue by DXA, g") + 
  theme_bw(base_size = 15, base_family = "Arial") +
  theme(panel.grid.major = element_blank(), panel.grid.minor = element_blank()) +
  theme(legend.position="none")

equation.a3=function(x){508+0.74*x} # published 
equation.a4=function(x){coef(mWATER_eb)[1]+coef(mWATER_eb)[2]*x} # own

pa2 <- ggplot(emptybody, aes(x=DXA_lean_eb, y=chem_water_eb)) +
  geom_point(shape=19, size=3, col="black") + 
  stat_function(fun=equation.a3,geom="line", color="darkred", linetype=2) + # published
  geom_smooth(method='lm', formula= y~x, color="black", linetype=1, size=.5) + # own
  stat_function(fun=equation.a4,geom="line", color="black", linetype=1) + # own
  ylim(0, 80000) + xlim(0, 100000) +
  ylab("Water, g") + xlab("Lean tissue by DXA, g") + 
  theme_bw(base_size = 15, base_family = "Arial") +
  theme(panel.grid.major = element_blank(), panel.grid.minor = element_blank()) +
  theme(legend.position="none")

equation.e1=function(x){-1384 + 0.216*x} # published
equation.e2=function(x){coef(mCP_eb)[1]+coef(mCP_eb)[2]*x} # own

pe1 <- ggplot(emptybody, aes(x=DXA_lean_eb, y=chem_CP_eb)) +
  geom_point(shape=19, size=3, col="black") + 
  stat_function(fun=equation.e1,geom="line", color="darkred", linetype=2) + # published
  geom_smooth(method='lm', formula= y~x, color="black", linetype=1, size=.5) + # own
  stat_function(fun=equation.e2,geom="line", color="black", linetype=1) + # own
  ylim(0, 22000) + xlim(0, 100000) +
  ylab("CP, g") + xlab("Lean tissue by DXA, g") + 
  theme_bw(base_size = 15, base_family = "Arial") +
  theme(panel.grid.major = element_blank(), panel.grid.minor = element_blank()) + 
  theme(legend.position="none")

equation.e3=function(x){2825+1.009*x} # published 
equation.e4=function(x){coef(mLipid_ebx)[1]+coef(mLipid_ebx)[2]*x} # own

pe2 <- ggplot(emptybody, aes(x=DXA_fat_eb, y=chem_lipid_eb)) +
  geom_point(shape=19, size=3, col="black") + 
  stat_function(fun=equation.e3,geom="line", color="darkred", linetype=2) + # published
  geom_smooth(method='lm', formula= y~x, color="black", linetype=1, size=.5) + # own
  stat_function(fun=equation.e4,geom="line", color="black", linetype=1) + # own
  ylim(0, 28000) + xlim(0, 18000) +
  ylab("Lipid, g") + xlab("Fat tissue by DXA, g") + 
  theme_bw(base_size = 15, base_family = "Arial") +
  theme(panel.grid.major = element_blank(), panel.grid.minor = element_blank()) +
  theme(legend.position="none")

equation.d1=function(x){3.371+1.116*x} # published
equation.d2=function(x){coef(mLipidkg_eb.1)[1]+coef(mLipidkg_eb.1)[2]*x} # own

pd1 <- ggplot(emptybody, aes(x=DXA_fatKG_eb, y=chem_lipidKG_eb)) +
  geom_point(shape=19, size=3, col="black") + 
  stat_function(fun=equation.d1,geom="line", color="darkred", linetype=2) + # published
  geom_smooth(method='lm', formula= y~x, color="black", linetype=1, size=.5) + # own
  stat_function(fun=equation.d2,geom="line", color="black", linetype=1) + # own
  ylim(0, 30) + xlim(0, 20) +
  ylab("Lipid, kg") + xlab("Fat tissue by DXA, kg") + 
  theme_bw(base_size = 15, base_family = "Arial") +
  theme(panel.grid.major = element_blank(), panel.grid.minor = element_blank()) + 
  theme(legend.position="none")

equation.d3=function(x){0.542+1.190*x} # published
equation.d4=function(x){coef(mAshkg_eb)[1]+coef(mAshkg_eb)[2]*x} # own

pd2 <- ggplot(emptybody, aes(x=DXA_BMCkg_eb, y=chem_ashKG_eb)) +
  geom_point(shape=19, size=3, col="black") + 
  stat_function(fun=equation.d3,geom="line", color="darkred", linetype=2) + # published
  geom_smooth(method='lm', formula= y~x, color="black", linetype=1, size=.5) + # own
  stat_function(fun=equation.d4,geom="line", color="black", linetype=1) + # own
  ylim(0, 5) + xlim(0, 3) +
  ylab("Ash, kg") + xlab("BMC by DXA, kg") + 
  theme_bw(base_size = 15, base_family = "Arial") +
  theme(panel.grid.major = element_blank(), panel.grid.minor = element_blank()) +
  theme(legend.position="none")

equation.d5=function(x){-0.894+0.219*x} # published
equation.d6=function(x){coef(mCPkg_eb)[1]+coef(mCPkg_eb)[2]*x} # own

pd3 <- ggplot(emptybody, aes(x=DXA_leanKG_eb, y=chem_CPkg_eb)) +
  geom_point(shape=19, size=3, col="black") + 
  stat_function(fun=equation.d5,geom="line", color="darkred", linetype=2) + # published
  geom_smooth(method='lm', formula= y~x, color="black", linetype=1, size=.5) + # own
  stat_function(fun=equation.d6,geom="line", color="black", linetype=1) + # own
  ylim(0, 25) + xlim(0, 100) +
  ylab("CP, kg") + xlab("Lean tissue by DXA, kg") + 
  theme_bw(base_size = 15, base_family = "Arial") +
  theme(panel.grid.major = element_blank(), panel.grid.minor = element_blank()) +
  theme(legend.position="none")

equation.d7=function(x){2.344+0.674*x} # published
equation.d8=function(x){coef(mWaterkg_eb)[1]+coef(mWaterkg_eb)[2]*x} # own

pd4 <- ggplot(emptybody, aes(x=DXA_leanKG_eb, y=chem_waterKG_eb)) +
  geom_point(shape=19, size=3, col="black") + 
  stat_function(fun=equation.d7,geom="line", color="darkred", linetype=2) + # published
  geom_smooth(method='lm', formula= y~x, color="black", linetype=1, size=.5) + # own
  stat_function(fun=equation.d8,geom="line", color="black", linetype=1) + # own
  ylim(0, 80) + xlim(0, 100) +
  ylab("Water, kg") + xlab("Lean tissue by DXA, kg") + 
  theme_bw(base_size = 15, base_family = "Arial") +
  theme(panel.grid.major = element_blank(), panel.grid.minor = element_blank()) + 
  theme(legend.position="none")

equation.d9=function(x){1.450+0.893*x} # published
equation.d10=function(x){coef(mProtWaterkg)[1]+coef(mProtWaterkg)[2]*x} # own

pd5 <- ggplot(emptybody, aes(x=DXA_leanKG_eb, y=protWaterChem_kg)) +
  geom_point(shape=19, size=3, col="black") + 
  stat_function(fun=equation.d9,geom="line", color="darkred", linetype=2) + # published
  geom_smooth(method='lm', formula= y~x, color="black", linetype=1, size=.5) + # own
  stat_function(fun=equation.d10,geom="line", color="black", linetype=1) + # own
  ylim(0, 100) + xlim(0, 100) +
  ylab("CP + water, kg") + xlab("Lean tissue by DXA, kg") + 
  theme_bw(base_size = 15, base_family = "Arial") +
  theme(panel.grid.major = element_blank(), panel.grid.minor = element_blank()) +
  theme(legend.position="none")
```

```{r eval=FALSE, warning=FALSE, message=FALSE}
multiplot(pa1, pe1, pd3, pd1, pa2, pe2, pd4, pd2, cols = 2)

png("Fig5.png", width = 900, height = 1500)
multiplot(pa1, pe1, pd3, pd1, pa2, pe2, pd4, pd2, cols = 2)
dev.off()
```

## CARCASS

### b) ...Mitchell, Scholz, Pursel & Evock-Clover 1998 _J. Anim. Sci._ 

+ half-carcasses (10 - 51 kg from pigs slaughtered at approximately 30, 60, 90, and 120 kg; split at midline, head removed, right half scanned)
+ sex not specified
+ chem. analys.
+ g chem - g DXA 
+ Lunar DPXL instrument -  pencil-beam 

#### 1. CP

```{r}
Mit98_pred_CP_carc.b <- -145 + 0.23 * (carcass$DXA_lean_carcHalf)
mCP_carcHalf <- lm(chem_CP_carcHalf ~ DXA_lean_carcHalf, data = carcass)
```

__published equation:__

$$CP (g) = -145 + 0.23 \times DXAlean$$
$RMSD=$ `r round(rmse((carcass$chem_CP_carcHalf), Mit98_pred_CP_carc.b))` g

$rCV =$ `r round((rmse((carcass$chem_CP_carcHalf), Mit98_pred_CP_carc.b)*100)/mean((carcass$chem_CP_carcHalf)), 2)`% of the mean CP content of the carcass by chemical analyses

__own equation:__

$\hat{CP_{carc\frac{1}{2}}} =$ `r round(summary(mCP_carcHalf)$coefficients[1,1], 3)` + `r round(summary(mCP_carcHalf)$coefficients[2,1], 3)` $ \times DXAlean  $

$RMSD=$ `r round(rmse((carcass$chem_CP_carcHalf), summary(mCP_carcHalf)$coefficients[1,1] + summary(mCP_carcHalf)$coefficients[2,1] * carcass$DXA_lean_carcHalf))` g 

$rCV =$ `r round((rmse((carcass$chem_CP_carcHalf), summary(mCP_carcHalf)$coefficients[1,1] + summary(mCP_carcHalf)$coefficients[2,1] * carcass$DXA_lean_carcHalf)*100)/mean((carcass$chem_CP_carcHalf)), 2)`% of the mean CP content of the carcass by chemical analyses

$R^2 = $ `r round(summary(mCP_carcHalf)$adj, 2)`

#### 2. Water

```{r}
Mit98_pred_Water_carc.b <- 150 + 0.73 * (carcass$DXA_lean_carcHalf)
mWATER_carcHalf <- lm(chem_water_carcHalf ~ DXA_lean_carcHalf, data = carcass)
```

__published equation:__

$$Water (g) = 150 + 0.73 \times DXAlean$$

$RMSD=$ `r round(rmse((carcass$chem_water_carcHalf), Mit98_pred_Water_carc.b))` g

$rCV =$ `r round((rmse((carcass$chem_water_carcHalf), Mit98_pred_Water_carc.b)*100)/mean((carcass$chem_water_carcHalf)), 1)`% of the mean water content of the carcass by chemical analyses

__own equation:__

$\hat{Water_{carc\frac{1}{2}}} =$ `r round(summary(mWATER_carcHalf)$coefficients[1,1], 3)` + `r round(summary(mWATER_carcHalf)$coefficients[2,1], 3)` $ \times DXAlean $  
$RMSD=$ `r round(rmse((carcass$chem_water_carcHalf), summary(mWATER_carcHalf)$coefficients[1,1] + summary(mWATER_carcHalf)$coefficients[2,1] * carcass$DXA_lean_carcHalf))` g

$rCV =$ `r round((rmse((carcass$chem_water_carcHalf), summary(mWATER_carcHalf)$coefficients[1,1] + summary(mWATER_carcHalf)$coefficients[2,1] * carcass$DXA_lean_carcHalf)*100)/mean((carcass$chem_water_carcHalf)), 1)`% of the mean water content of the carcass by chemical analyses 

### c) ...Mitchell, Scholz & Pursel 2003 _Meat Science_ __carc__

+ half carcasses (right) (mean 43 kg)
+ sex not specified
+ chem. analys.
+ % chem - % DXA 
+ Lunar DPXL instrument -  pencil-beam 

#### 1. Lipid percent

```{r}
carcass$sumComponents_CarcHalf <- (carcass$chem_water_carcHalf + carcass$chem_ash_carcHalf + carcass$chem_CP_carcHalf + carcass$chem_lipid_carcHalf) 
# percentages: 100% is sum of components chemical, in DXA this is already provided
carcass$category <- as.factor(carcass$category)

Mit03_pred_Fat_carc.d <- 13.5 + 0.582 * (carcass$DXA_fatPerc_carc)
carcass$percLipid_carcHalf <- (carcass$chem_lipid_carcHalf/(carcass$sumComponents_CarcHalf))*100 

mLipidPerc_carc <- lm(percLipid_carcHalf ~ DXA_fatPerc_carc, data = carcass)
```

__published equation:__

$$\%Lipid = 13.5 + 0.582 \times \% DXAfat $$

$RMSE=$ `r round(rmse(carcass$percLipid_carcHalf, Mit03_pred_Fat_carc.d), 1)` % 

__own equation:__

$ \% Lipid = $ `r round(summary(mLipidPerc_carc)$coefficients[1,1], 3)` + `r round(summary(mLipidPerc_carc)$coefficients[2,1], 3)` $ \times \%Fat $

$RMSE=$ `r round(rmse(carcass$percLipid_carcHalf, summary(mLipidPerc_carc)$coefficients[1,1] + summary(mLipidPerc_carc)$coefficients[2,1] * carcass$DXA_fatPerc_carc), 1)` % 

$R^2 = $ `r round(summary(mLipidPerc_carc)$adj, 2)`

#### 2. CP + water percent

```{r}
Mit03_pred_Lean_carc.d <- 25.2 + 0.584 * (carcass$DXA_leanPerc_carc)
carcass$percLeanTissue_carcHalf <- ((carcass$chem_CP_carcHalf + carcass$chem_water_carcHalf)/(carcass$sumComponents_CarcHalf)) * 100

mLeanTissPerc_carc <- lm(percLeanTissue_carcHalf ~ DXA_leanPerc_carc, data = carcass)
```

__published equation:__

$$\%(water + CP) = 25.2 + 0.584 \times \% Lean $$

$RMSE=$ `r round(rmse(carcass$percLeanTissue_carcHalf, Mit03_pred_Lean_carc.d), 1)` % 

__own equation:__

$\% Lean \; tissue \; (water + CP)= $ `r round(summary(mLeanTissPerc_carc)$coefficients[1,1], 3)` + `r round(summary(mLeanTissPerc_carc)$coefficients[2,1], 3)` $\times \%Lean$

$RMSE=$ `r round(rmse(carcass$percLeanTissue_carcHalf, summary(mLeanTissPerc_carc)$coefficients[1,1] + summary(mLeanTissPerc_carc)$coefficients[2,1] * carcass$DXA_leanPerc_carc), 1)` %

$R^2 = $ `r round(summary(mLeanTissPerc_carc)$adj, 2)`

## Figure 6

```{r}
equation.b1=function(x){-145+0.23*x} # published
equation.b2=function(x){coef(mCP_carcHalf)[1]+coef(mCP_carcHalf)[2]*x} # own

pb1 <- ggplot(carcass, aes(x=DXA_lean_carcHalf, y=chem_CP_carcHalf)) + 
  geom_point(shape=19, size=3, col="black") + 
  stat_function(fun=equation.b1,geom="line", color="darkred", linetype=2) + # published
  geom_smooth(method='lm', formula= y~x, color="black", linetype=1, size=.5) + # own
  stat_function(fun=equation.b2,geom="line", color="black", linetype=1) + # own
  ylim(0, 8500) + xlim(0, 40000) +
  ylab("CP, g") + xlab("Lean tissue by DXA, g") + 
  theme_bw(base_size = 15, base_family = "Arial") +
  theme(panel.grid.major = element_blank(), panel.grid.minor = element_blank()) +
  theme(legend.position="none")

equation.b3=function(x){150+0.73*x} # published 
equation.b4=function(x){coef(mWATER_carcHalf)[1]+coef(mWATER_carcHalf)[2]*x} # own

pb2 <- ggplot(carcass, aes(x=DXA_lean_carcHalf, y=chem_water_carcHalf)) +
  geom_point(shape=19, size=3, col="black") + 
  stat_function(fun=equation.b3,geom="line", color="darkred", linetype=2) + # published
  geom_smooth(method='lm', formula= y~x, color="black", linetype=1, size=.5) + # own
  stat_function(fun=equation.b4,geom="line", color="black", linetype=1) + # own
  ylim(0, 30000) + xlim(0, 40000) +
  ylab("Water, g") + xlab("Lean tissue by DXA, g") + 
  theme_bw(base_size = 15, base_family = "Arial") +
  theme(panel.grid.major = element_blank(), panel.grid.minor = element_blank()) +
  theme(legend.position="none")

equation.c1=function(x){13.5+0.582*x} # published 
equation.c2=function(x){coef(mLipidPerc_carc)[1]+coef(mLipidPerc_carc)[2]*x} # own

pc1 <- ggplot(carcass, aes(x=DXA_fatPerc_carc, y=lipid_perc_carc, color = category)) +
  geom_point(shape=19, size=3) + 
  scale_color_manual(values=c("#d2cccc", "#636161", "#000000")) +
  stat_function(fun=equation.c1,geom="line", color="darkred", linetype=2) + # published
  geom_smooth(method='lm', formula= y~x, color="black", linetype=1, size=.5) + # own
  stat_function(fun=equation.c2,geom="line", color="black", linetype=1) + # own
  ylim(0, 35) + xlim(0, 25) +
  ylab("Lipid %") + xlab("Fat % by DXA") + 
  theme_bw(base_size = 15, base_family = "Arial") +
  theme(panel.grid.major = element_blank(), panel.grid.minor = element_blank()) +
  theme(legend.position="none")

equation.c3=function(x){25.2+0.584*x} # published 
equation.c4=function(x){coef(mLeanTissPerc_carc)[1]+coef(mLeanTissPerc_carc)[2]*x} # own

pc2 <- ggplot(carcass, aes(x=DXA_leanPerc_carc, y=lean_perc_carc, colour = category)) +
  geom_point(shape=19, size=3) + 
  scale_color_manual(values=c("#d2cccc", "#636161", "#000000")) +
  stat_function(fun=equation.c3,geom="line", color="darkred", linetype=2) + # published
  geom_smooth(method='lm', formula= y~x, color="black", linetype=1, size=.5) + # own
  stat_function(fun=equation.c4,geom="line", color="black", linetype=1) + # own
  ylim(0, 100) + xlim(0, 100) + # axes here not spanning from zero to highest value - too little variation
  ylab("CP + water %") + xlab("Lean % by DXA") + 
  theme_bw(base_size = 15, base_family = "Arial") +
  theme(panel.grid.major = element_blank(), panel.grid.minor = element_blank()) +
  theme(legend.position="none")
```

```{r eval=FALSE, warning=FALSE, message=FALSE}
multiplot(pb1, pc2, pb2, pc1, cols = 2)

png("Fig6.png", width = 800, height = 665)
multiplot(pb1, pc2, pb2, pc1, cols = 2)
dev.off()
```


# REGRESSION ON PERCENTAGES (see Supplementary Material section 6)

## Table S3

```{r}
carcass$sumComponents_CarcHalf <- (carcass$chem_water_carcHalf + carcass$chem_ash_carcHalf + carcass$chem_CP_carcHalf + carcass$chem_lipid_carcHalf) 

# percentages: 100% is sum of components chemical, in DXA this is already provided
carcass$percLipid_carcHalf <- (carcass$chem_lipid_carcHalf/(carcass$sumComponents_CarcHalf))*100 
carcass$percLeanTissue_carcHalf <- ((carcass$chem_CP_carcHalf + carcass$chem_water_carcHalf)/(carcass$sumComponents_CarcHalf)) * 100

mLipidPerc_carc <- lm(percLipid_carcHalf ~ DXA_fatPerc_carc, data = carcass)
mLeanTissPerc_carc <- lm(percLeanTissue_carcHalf ~ DXA_leanPerc_carc, data = carcass)

tableS3 <- cbind(as.data.frame(c("Intercept", "% lean", "Intercept", "% lipid")),
                               c(paste0(round(as.numeric(coef(mLeanTissPerc_carc)[1]), 3), " (", round(summary(mLeanTissPerc_carc)$coefficients[1,2], 3), ")"),
                                 paste0(round(as.numeric(coef(mLeanTissPerc_carc)[2]), 3), " (", round(summary(mLeanTissPerc_carc)$coefficients[2,2], 3), ")"),
                                 paste0(round(as.numeric(coef(mLipidPerc_carc)[1]), 3), " (", round(summary(mLipidPerc_carc)$coefficients[1,2], 3), ")"),
                                 paste0(round(as.numeric(coef(mLipidPerc_carc)[2]), 3), " (", round(summary(mLipidPerc_carc)$coefficients[2,2], 3), ")")
                                ),
                 c(round(summary(mLeanTissPerc_carc)$coefficients[1,4], 3), "< 0.001",
                 round(summary(mLipidPerc_carc)$coefficients[1,4], 3), "< 0.001"),
                 c(round(summary(mLeanTissPerc_carc)$adj.r.sq, 3), NA, round(summary(mLipidPerc_carc)$adj.r.sq, 3), NA),
                 c(round(rmse(carcass$percLeanTissue_carcHalf, summary(mLeanTissPerc_carc)$coefficients[1,1] +
                                       summary(mLeanTissPerc_carc)$coefficients[2,1] * carcass$DXA_leanPerc_carc), 2), NA,
                 round(rmse(carcass$percLipid_carcHalf, summary(mLipidPerc_carc)$coefficients[1,1] + 
                                     summary(mLipidPerc_carc)$coefficients[2,1] * carcass$DXA_fatPerc_carc), 2), NA))
  
names(tableS3) <- c("term (DXA variable)", "estimate (SE)", "P", "R^2", "RMSE")  
write.table(tableS3, file = "tableS3.txt", sep = "\t")
```
````
