## Supplementary Material for "Accuracy of predicting chemical body composition of growing pigs using dual-energy X-ray absorptiometry"

### 1. Image processing for DXA

Images were processed after scanning using the encore software (version 16). The regions of interest (ROI) for the live pig were placed according to the manufacturer's instruction in a way that was as similar as possible to humans (Figure S1). Artefacts, such as the metallic ear tag and the plastic mask used to keep animals under anaesthesia, were removed during image processing.

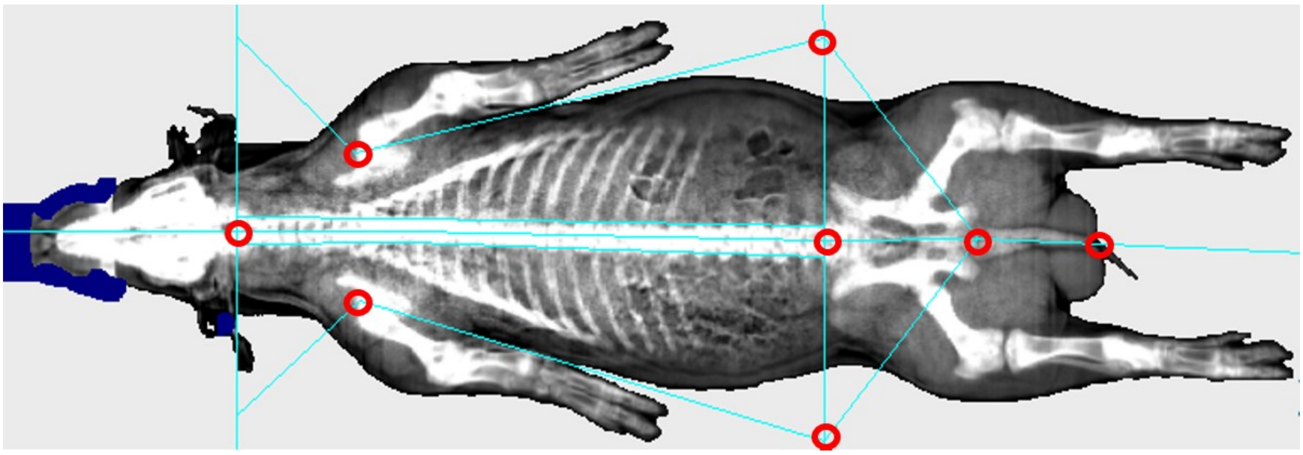

**Figure S1:** Placement of the regions of interest (ROI; areas delimited by turquoise lines) and the reference points (red circles) on scans of live pigs. Artefacts (ear tag and mask) are highlighted in blue.

### 2. Raw data check of chemical analysis data

Chemical data were first checked by subtracting the sums of ash, crude protein, fat and water from the body weight obtained by scales. The mean difference in empty body weight was -57.44 ± 924.49 g and in carcass weight -63.91 ± 408.71 g. In addition, we visually examined the histograms of the proportions of water, ash, CP and lipid of the total sum of these components in connection with the coefficients of variation to check the distribution of the data and (Fig. S1).

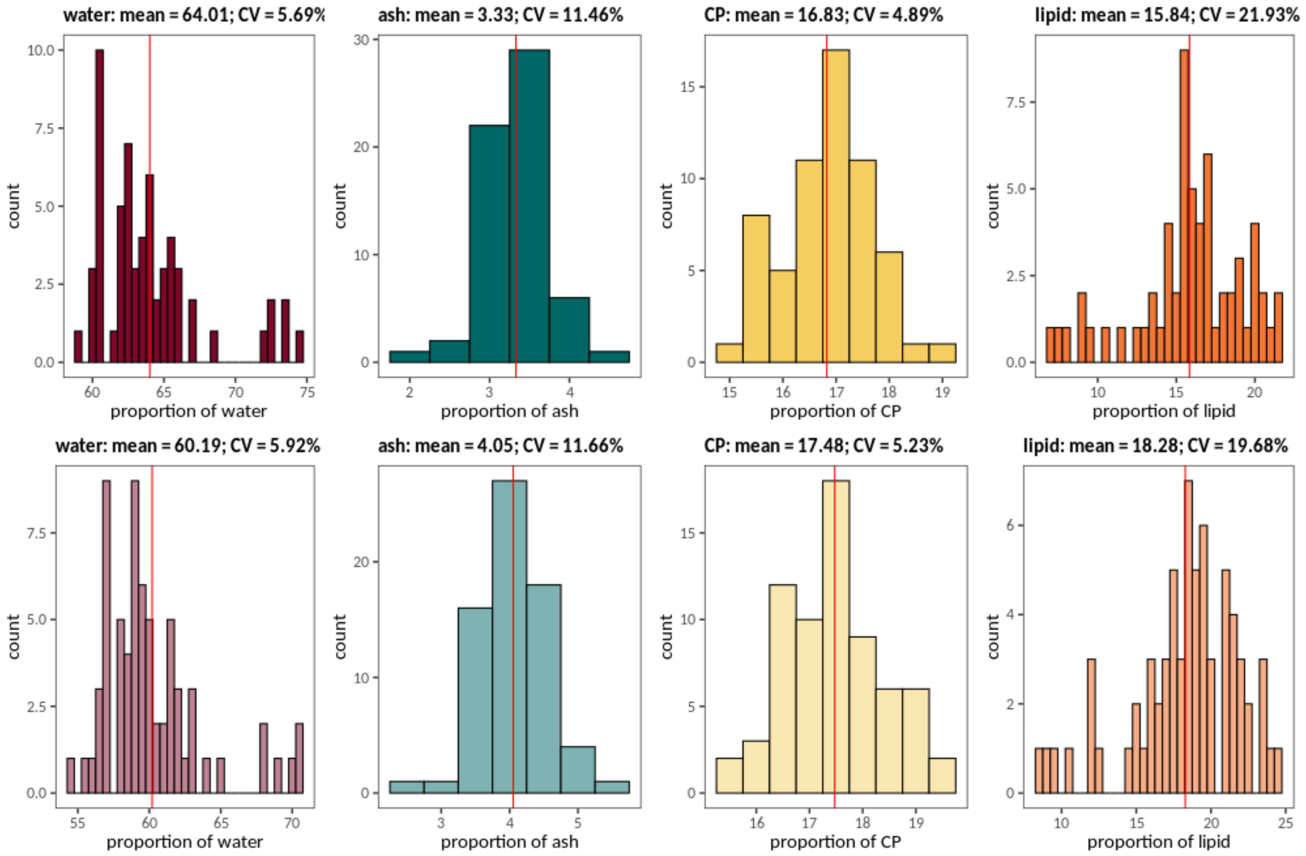

**Figure S2:** Histograms and coefficients of variation (CV) of the proportions of water (red), ash (green), CP (yellow) and lipid (orange) determined by wet chemistry in the empty body (upper row, dark colours) and carcass (lower row, light colours) of entire male pigs. For code and data underlying the figure, see Kasper et al. (2020) and Supplementary Material section 6.

#### 3. Dietary treatments

Data used in this study originates from a so far unpublished experiment in which pigs were fed two grower and two finisher diets differing in their CP content (see Materials and Methods section in the main text for details). In this article, we were not specifically interested in the effect of dietary treatment. It was thus necessary to verify whether dietary treatments affected the estimates of slopes and intercepts of the prediction equations for chemical body composition based on DXA values. Table S1 shows the influence of dietary treatments on intercepts and slopes and the p-values of the treatment term. The treatment effect was only significant in the model of the energy content of the carcass, but the $R^2$-value of the model including the treatment effect was only marginally higher than in the uncorrected model. Intercepts differed slightly between models containing the treatment effect and those that did not, but slopes remained largely unchanged (except for ash and lipid). Therefore, we concluded that, for the purpose of the present study, treatment could be omitted from the regression equations without compromising the estimates of slopes and intercepts.

**Table S1:** Effects of the inclusion of dietary treatment in the prediction equations on intercept, slope(s) and R2. Δintercept: difference in intercept, Δslope: difference in slope, $\Delta R^2$: difference in $R^2$ of corrected model (including treatment effect) minus uncorrected model (without treatment effect). p-value: p-value of the treatment effect when included in the model.

| | empty body[a] | | | | carcass | | | |
|---|---|---|---|---|---|---|---|---|
| | **ΔIntercept** | **ΔSlope** | **ΔR²** | **p-value** | **ΔIntercept** | **ΔSlope** | **ΔR²** | **p-value** |
| weight (g) | 17.963 | 0.000 | 0.000 | 0.051 | 12.997 | 0.000 (mass) | 0.000 | 0.134 |
| energy content (MJ) | 0.318 | 0.000 (lean)<br>-0.001 (fat) | 0.000 | 0.604 | 0.723 | 0.000 (lean)<br>-0.001 (fat) | 0.001 | 0.039 |
| water (g) | -29.298 | -0.001 (lean) | 0.000 | 0.257 | -41.839 | -0.002 (lean) | 0.000 | 0.062 |
| ash (g) | -23.59 | 0.010 (BMC) | 0.000 | 0.387 | 3.438 | -0.012 (BMC)<br>0.001 (lean) | -0.002 | 0.793 |
| Ca (g) | 0.198 | 0.000 (BMC) | -0.003 | 0.985 | 2.622 | 0.001 (BMC) | -0.003 | 0.808 |
| P (g) | 0.911 | -0.008 (BMC)<br>0.000 (lean) | -0.001 | 0.572 | 2.451 | - 0.009(BMC)<br>0.001 (lean) | 0.000 | 0.299 |
| CP (g) | 2.821 | 0.000 (lean) | 0.000 | 0.890 | 3.775 | 0.000 (lean) | 0.000 | 0.849 |
| N (g) | 0.451 | 0.000 (lean) | 0.000 | 0.890 | 0.604 | 0.000 (lean) | 0.000 | 0.849 |
| lipid (g) | 9.145 | 0.011 (fat)<br>-0.003 (lean) | 0.000 | 0.621 | 18.189 | -0.030 (fat)<br>0.011 (lean) | 0.001 | 0.091 |

[a]empty body contents at the time of live DXA scans estimated from empty body contents at slaughter

#### 4. Model diagnostics

Model diagnostics were carried out using the packages car (version 3.0-8; Fox & Weisberg, 2019) and MASS (version 7.3-51.6; Venables & Ripley, 2002) in R (version 3.6.3; R Core Team, 2020). They revealed that some assumptions of linear models were violated. In particular, the assumption of homogeneity of variance was not met judging from visual inspection of residuals vs. fitted values. Variances for larger fitted values were higher than those of lower fitted values in all models. In some of the models[1], we detected statistical outliers, but the assessment of the Cook's distance showed that none of them seemed to have a pronounced influence on the models. Removal of those outliers never resulted in improved model diagnostics. One individual (8729-FE3), which was an outlier in five of the models (ash, Ca and P), was ranked second highest in ash, Ca and P content per g body weight. Another individual (9122-FE3), which was an outlier in the model predicting water from lean mass, had a rather high, but not extreme, lean mass in DXA. Both individuals did not have unusual chemical or DXA values and there was no indication of errors, thus we did not remove these individuals from the dataset. Visual checks revealed that residuals were more or less normally distributed. The lm() procedure in R is considered quite robust, but fitting models with linear least-squares regression in

[1] Body weight and Ca in the carcass, energy, water, ash content in empty body, Ca, P in empty body; empty body ash and lipid contents predicted from carcass

order to derive the parameters of the regression line while violating core assumptions might bear the risk of obtaining unreliable slopes and intercepts for our prediction equations. Thus, we used a "leave-one-out" cross-validation procedure in the package caret (version 6.0-86; Kuhn, 2020) to construct a non-parametric confidence interval for the slopes and intercepts, which are presented in Table 3 in the main text. To obtain the confidence intervals, we computed intercepts and slopes of all models by omitting one row at a time, ordered the estimates according to their value and took the 3$^{rd}$ and 59$^{th}$ and the 3$^{rd}$ and 66$^{th}$ ranked estimates for models of the empty body and the carcass, respectively, as the limits of the confidence intervals. This corresponds to a 93% CI in the empty body and a 94% CI in the carcass.

#### 5. Regional regressions

The main goal of the study was to build calibration equations to relate DXA measurements to chemically determined values of body composition and single nutrients. However, the choice of the range of body weights might affect the slopes and intercepts of those global regression equations. In particular for ash, calcium and phosphorus, the slopes might differ between regressions on a subset of data for each target weight (20, 60 and 100 kg live weight) and the global regression using the full dataset (see Figures 2 and 3 in the main text). To explore this possibility, we compared regression coefficients and $R^2$ between those types of models by building regressions for each target weight category separately (Table S2). It has to be noted that the interpretation of the regional regressions must be treated with caution because the sample sizes are extremely small, especially in the case of the 20 kg category. For most variables, the regional regression lines overlapped reasonably well with the global regression line, both for the empty body (Figure S3) and the carcass (Figure S4), even though the numerical values of intercepts and slopes differed somewhat between the subsets. Thus, it seems that the relationships between chemical measurements and DXA values are mostly linear across weight categories and in agreement with the global one. However, in the case of ash and phosphorus, there were considerable differences between the slopes of the regional models and the global ones. The present data set is not able to clarify, however, whether these deviations are caused by non-linear relationships or by the comparatively high variation in the heavier slaughter weight categories. The latter is supported by the relatively low $R^2$ for the subsets of ash, calcium and phosphorus models, suggesting that more data is needed to fit reliable regression equations for bone minerals.

**Table S2:** Regression equations using the full dataset (across slaughter weight categories) and subsets (target slaughter weight categories of 20, 60 and 100 kg) for empty body and carcass. The sample size (N), the intercept and the slopes (x1: first variable, x2: second variable) of the respective regression, as well as their fit (R2) and error (RMSE, root mean square error, and rCV, the residual coefficient of variation in percent of the chemically determined variable) are given.

| | | | **empty body** | | | | | | | **carcass** | | | | | |
|---|---|---|---|---|---|---|---|---|---|---|---|---|---|---|---|
| | | **N** | **intercept** | **x1** | **x2** | **$R^2$** | **RMSE** | **rCV** | **N** | **intercept** | **x1** | **x2** | **$R^2$** | **RMSE** | **rCV** |
| Weight | full data | 61 | -149.059 | 0.988 | | 1.000 | 150.081 | 0.20% | 61 | 172.744 | 0.991 | | 1.000 | 151.357 | 0.20% |
| | *20 kg* | *6* | *-185.997* | *0.991* | | *0.998* | *32.057* | *0.20%* | *6* | *233.391* | *0.989* | | *0.995* | *45.707* | *0.30%* |
| | 60 kg | 18 | 1309.366 | 0.963 | | 0.999 | 66.787 | 0.10% | 18 | 486.190 | 0.983 | | 0.998 | 95.369 | 0.20% |
| | 100 kg | 37 | 136.093 | 0.985 | | 0.997 | 180.988 | 0.20% | 37 | 1505.524 | 0.973 | | 0.995 | 169.375 | 0.20% |
| Energy content | full data | 61 | -34.940 | 0.007 | 0.038 | 0.995 | 22.904 | 2.90% | 61 | -36.077 | 0.007 | 0.036 | 0.990 | 27.102 | 3.80% |
| | *20 kg* | *6* | *-74.779* | *0.006* | *0.062* | *0.968* | *1.579* | *1.30%* | *6* | *-55.346* | *0.010* | *0.032* | *0.936* | *2.319* | *2.20%* |
| | 60 kg | 18 | 3.856 | 0.005 | 0.041 | 0.832 | 14.239 | 2.70% | 18 | -111.295 | 0.009 | 0.038 | 0.889 | 12.034 | 2.50% |
| | 100 kg | 37 | -113.483 | 0.007 | 0.039 | 0.873 | 26.810 | 2.60% | 37 | 200.683 | 0.004 | 0.033 | 0.776 | 31.637 | 3.50% |
| Water | full data | 61 | 1254.428 | 0.682 | | 0.999 | 578.904 | 1.20% | 61 | 1058.187 | 0.720 | | 0.998 | 545.643 | 1.50% |
| | *20 kg* | *6* | *1939.931* | *0.638* | | *0.972* | *68.884* | *0.50%* | *6* | *1910.808* | *0.643* | | *0.945* | *87.001* | *0.90%* |
| | 60 kg | 18 | -204.679 | 0.712 | | 0.940 | 376.450 | 1.10% | 18 | 3347.429 | 0.656 | | 0.956 | 266.983 | 1.00% |
| | 100 kg | 37 | 36.805 | 0.697 | | 0.934 | 689.194 | 1.20% | 37 | 834.485 | 0.723 | | 0.896 | 650.484 | 1.50% |
| Ash | full data | 61 | 32.300 | 1.322 | | 0.886 | 316.734 | 12.60% | 61 | -33.611 | 0.807 | 0.018 | 0.875 | 311.888 | 12.50% |
| | *20 kg* | *6* | *-436.966* | *2.395* | | *0.548* | *19.939* | *3.60%* | *6* | *-224.067* | *0.759* | *0.037* | *0.574* | *18.409* | *3.40%* |
| | 60 kg | 18 | 817.101 | 0.745 | | 0.249 | 106.052 | 5.80% | 18 | 862.581 | 0.642 | -0.001 | 0.154 | 115.293 | 6.40% |
| | 100 kg | 37 | 552.332 | 1.105 | | 0.252 | 394.956 | 12.50% | 37 | -498.296 | 0.831 | 0.024 | 0.244 | 378.821 | 12.40% |
| Calcium | full data | 61 | -2.807 | 0.407 | | 0.829 | 123.372 | 16.20% | 61 | 19.401 | 0.377 | | 0.799 | 132.045 | 16.50% |
| | *20 kg* | *6* | *-41.879* | *0.476* | | *0.015* | *10.144* | *6.50%* | *6* | *138.946* | *0.059* | | *-0.242* | *13.880* | *8.40%* |
| | 60 kg | 18 | 64.351 | 0.362 | | 0.209 | 56.709 | 10.10% | 18 | 112.608 | 0.313 | | 0.190 | 59.434 | 10.20% |
| | 100 kg | 37 | 19.144 | 0.397 | | 0.222 | 153.189 | 16.00% | 37 | 280.189 | 0.275 | | 0.154 | 156.998 | 16.10% |
| Phosphorus | full data | 61 | -9.819 | 0.145 | 0.003 | 0.893 | 52.199 | 12.00% | 61 | -6.388 | 0.109 | 0.004 | 0.863 | 56.532 | 13.00% |
| | *20 kg* | *6* | -169.630 | 0.626 | 0.000 | 0.378 | 5.429 | 5.70% | *6* | -95.492 | 0.212 | 0.008 | 0.438 | 5.407 | 5.80% |
| | 60 kg | 18 | 137.288 | 0.271 | -0.004 | 0.368 | 23.665 | 7.50% | 18 | 114.153 | 0.234 | -0.004 | 0.319 | 26.668 | 8.50% |
| | 100 kg | 37 | -91.840 | 0.142 | 0.004 | 0.203 | 64.257 | 11.80% | 37 | -15.870 | 0.102 | 0.005 | 0.136 | 67.408 | 12.70% |
| Crude protein | full data | 61 | -784.767 | 0.200 | | 0.990 | 454.075 | 3.60% | 61 | -482.745 | 0.230 | | 0.983 | 476.853 | 4.40% |
| | *20 kg* | *6* | *494.815* | *0.131* | | *0.957* | *17.617* | *0.70%* | *6* | *365.097* | *0.159* | | *0.912* | *27.658* | *1.30%* |
| | 60 kg | 18 | -2439.291 | 0.232 | | 0.615 | 376.780 | 4.10% | 18 | -625.328 | 0.233 | | 0.497 | 430.488 | 5.60% |
| | 100 kg | 37 | -250.492 | 0.194 | | 0.663 | 513.707 | 3.20% | 37 | -1292.663 | 0.243 | | 0.598 | 523.128 | 3.90% |
| Nitrogen | full data | 61 | -125.563 | 0.032 | | 0.990 | 72.652 | 3.60% | 61 | -77.239 | 0.037 | | 0.983 | 76.296 | 4.40% |
| | *20 kg* | *6* | *79.170* | *0.021* | | *0.957* | *2.819* | *0.70%* | *6* | *58.415* | *0.025* | | *0.912* | *4.425* | *1.30%* |
| | 60 kg | 18 | -390.287 | 0.037 | | 0.615 | 60.285 | 4.10% | 18 | -100.053 | 0.037 | | 0.497 | 68.878 | 5.60% |
| | 100 kg | 37 | -40.079 | 0.031 | | 0.663 | 82.193 | 3.20% | 37 | -206.826 | 0.039 | | 0.598 | 83.700 | 3.90% |
| Lipid | full data | 61 | -294.210 | 0.986 | 0.042 | 0.984 | 690.750 | 5.50% | 61 | -538.430 | 0.933 | 0.049 | 0.970 | 828.339 | 7.00% |
| | *20 kg* | *6* | *-1882.493* | *1.509* | *0.074* | *0.989* | *18.146* | *1.20%* | *6* | *-1402.659* | *0.717* | *0.152* | *0.966* | *33.371* | *2.40%* |
| | 60 kg | 18 | 1054.591 | 1.041 | 0.014 | 0.679 | 511.951 | 6.30% | 18 | -1806.169 | 0.907 | 0.088 | 0.705 | 481.901 | 6.40% |
| | 100 kg | 37 | -5400.279 | 1.070 | 0.089 | 0.879 | 757.841 | 4.60% | 37 | 5355.090 | 0.860 | -0.033 | 0.756 | 960.044 | 6.40% |

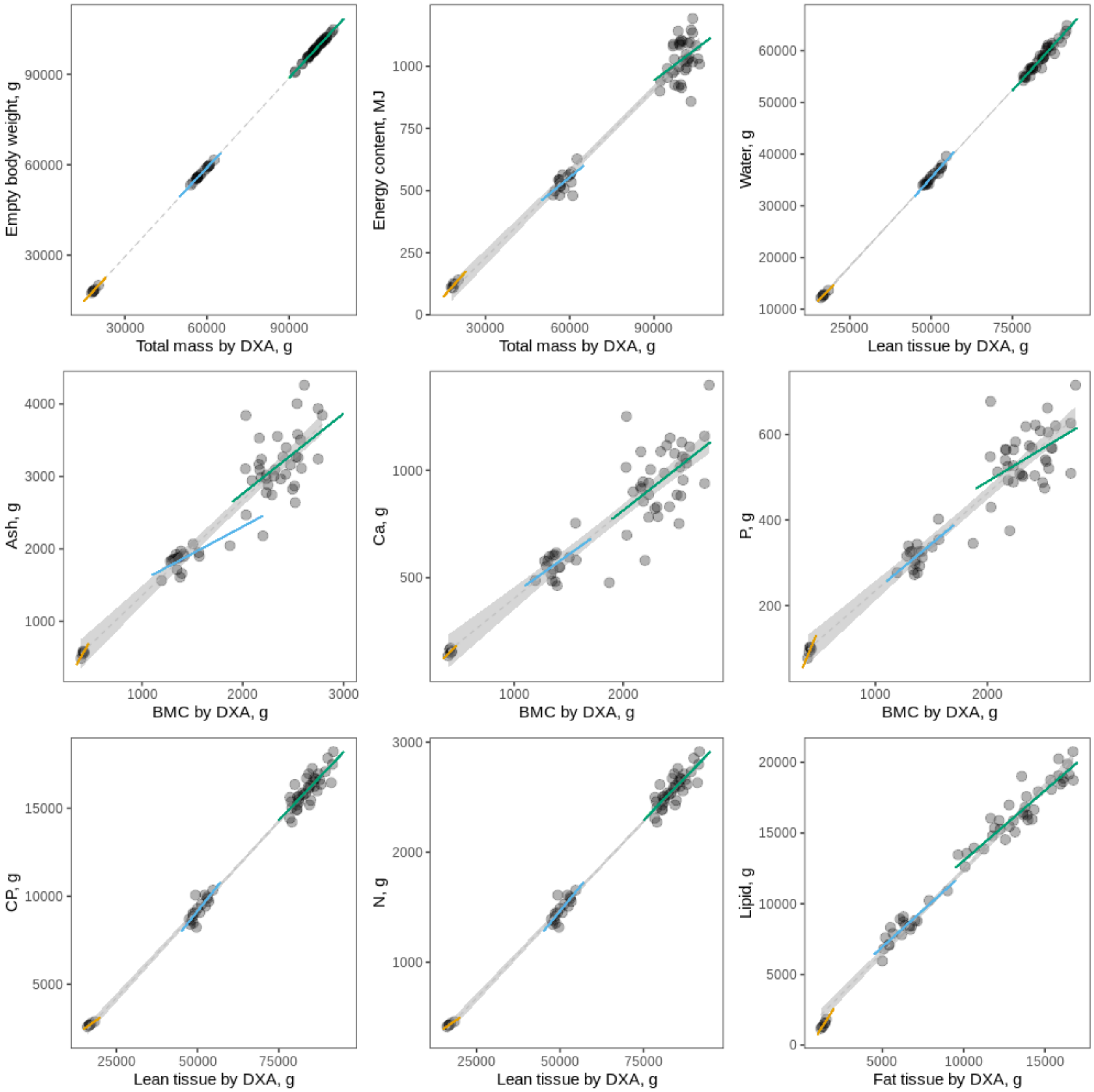

**Figure S3:** Global and regional regression equations in the empty body of weight and chemical composition according to DXA values in growing pigs. Dots represent individual data points, and the grey dotted line the global regression line and the 95% confidence bands shaded in grey. Yellow, blue and green solid lines represent the regression lines of the 20kg, 60 kg and 100 kg target weight category, respectively. For energy, the regression lines for reduced models including only total mass by DXA (instead of lean and fat from DXA) are shown. Also, for P and lipid, only the lines of the reduced regressions including BMC and fat by DXA, respectively, are shown.

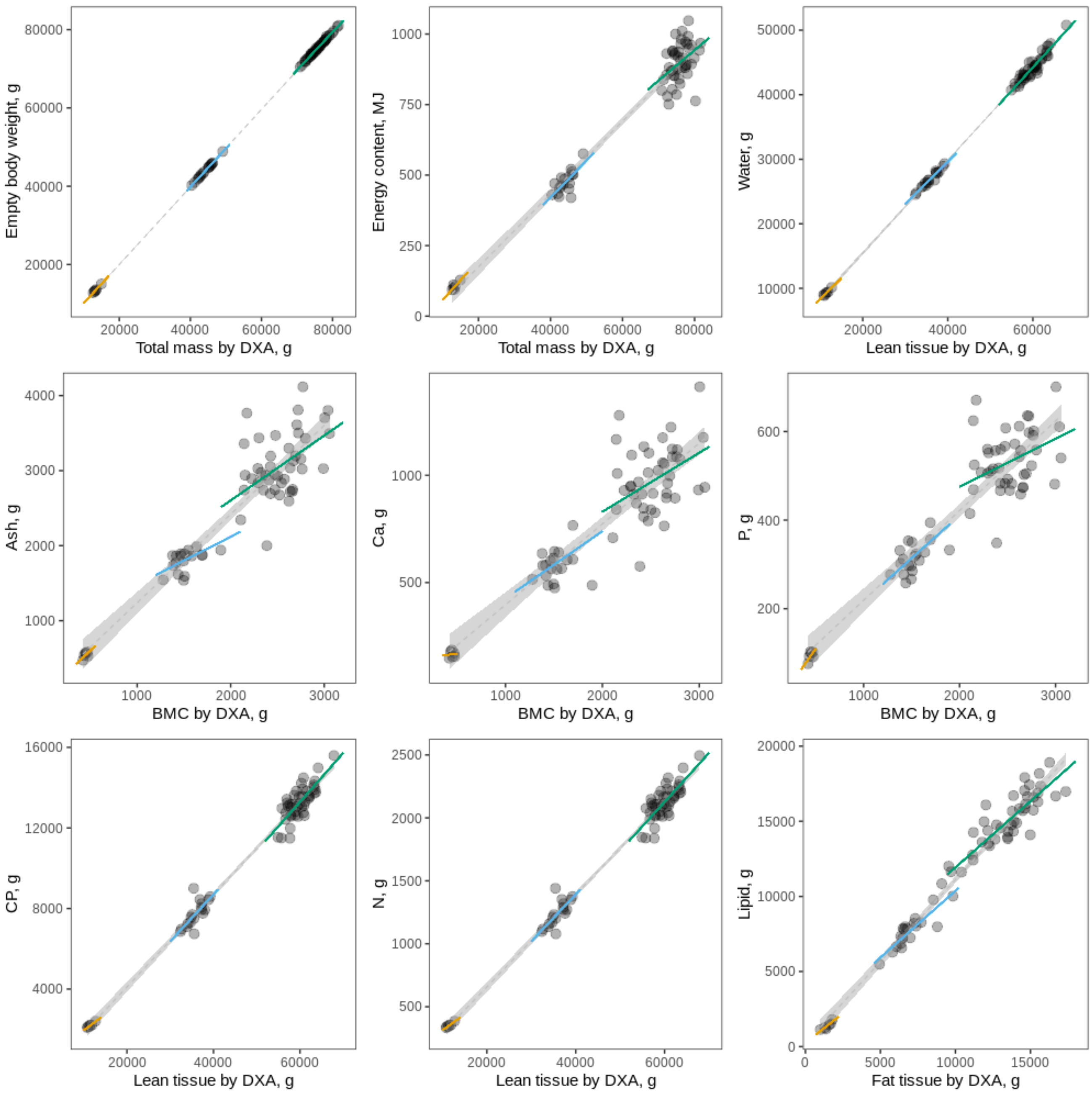

**Figure S4:** Global and regional regression equations in the carcass of weight and chemical composition according to DXA values in growing pigs. Dots represent individual data points, and the grey dotted line the global regression line and the 95% confidence bands shaded in grey. Yellow, blue and green solid lines represent the regression lines of the 20kg, 60 kg and 100 kg target weight category, respectively. For energy, the regression lines for reduced models including only total mass by DXA (instead of lean and fat from DXA) are shown. Also, for ash, P and lipids, only the lines of the reduced regressions including BMC and fat by DXA, respectively, are shown.

#### 6. Regressions on percentages

For a comparison with the percentages of lean tissue mass and lipid in the carcass halves by (Mitchell et al., 2003), we built separate prediction equations. We regressed the percentage of lean tissue by DXA on the percentage of lean tissue mass, which was calculated as the sum of CP and water divided by sum of CP, lipid, water and ash of the half carcasses. The percentage of fat tissue mass by DXA was regressed on the percentage of lipid content in the half carcasses (lipid divided by sum of CP, lipid, water and ash). Table S2 shows the prediction equations derived for % lean $tissue_{DXA}$ and % $lipid_{DXA}$.

**Table S3:** Summary of model estimates of percentages in the half-carcasses including their standard error, their p-value, $R^2$ and RMSE from back-prediction of chemical values.

| chemical variable (predicted variable) | term (DXA variable) | estimate (SE) | p-value | $R^2$ | RMSE |
|---|---|---|---|---|---|
| **% lean tissue (CP + water)** | Intercept | -7.414 (4.501) | 0.104 | 0.842 | 1.40 |
| **(% $Lean_{DXA}$)** | % lean | 1.061 (0.056) | < 0.001 | | |
| **% lipid** | Intercept | 0.565 (0.957) | 0.557 | 0.841 | 1.41 |
| **(% $lipid_{DXA}$)** | % fat | 1.074 (0.057) | < 0.001 | | |

#### 7. Code for analyses in R

*[see Kasper_etal_DXA.Rmd as a separate file]*
